# Monitoring correlates of SARS-CoV-2 infection in cell culture using two-photon microscopy and a novel fluorescent calcium-sensitive dye

**DOI:** 10.1101/2022.09.12.506773

**Authors:** Domokos Máthé, Gergely Szalay, Levente Cseri, Zoltán Kis, Bernadett Pályi, Gábor Földes, Noémi Kovács, Anna Fülöp, Áron Szepesi, Polett Hajdrik, Attila Csomos, Ákos Zsembery, Kristóf Kádár, Gergely Katona, Zoltán Mucsi, Balázs József Rózsa, Ervin Kovács

## Abstract

The organism-wide effects of viral infection SARS-CoV-2 are well studied, but little is known about the dynamics of how the infection spreads in time among or within cells due to the scarcity of suitable high-resolution experimental systems. Two-photon (2P) imaging combined with a proper subcellular staining technique has been an effective tool for studying mechanisms at such resolutions and organelle levels. Herein, we report the development of a novel calcium sensor molecule along with a 2P-technique for identifying imaging patterns associated with cellular correlates of infection damage within the cells. The method works as a cell viability assay and also provides valuable information on how the calcium level and intracellular distribution are perturbed by the virus. Moreover, it allows the quantitative analysis of infection dynamics. This novel approach facilitates the study of the infection progression and the quantification of the effects caused by viral variants and viral load.

## Introduction

The current global epidemic — caused by the novel severe acute respiratory syndrome coronavirus 2 (SARS-CoV-2) — has highlighted the lack of in vivo experimental techniques for commensurable methods that could enable the monitoring of viral infections at cellular or subcellular levels^1^. In its first two years (as of 02/09/2022), the pandemic caused more than 600 million confirmed cases and 6.4 million deaths^2^. Based on current trends, the disease is not expected to disappear anytime soon due to the appearance of successful virus variants that increasingly evade existing immunization, like the delta or the world-wide surge of different omicron variants^3^. Nonetheless, even without acute infections, clinical and experimental evidence shows that long-term effects of SARS-CoV-2 infection, sometimes referred to as post-acute sequelae of COVID-19 or, more colloquially, the long COVID or post-COVID syndrome are affecting many people and are scarcely scrutinized. Most frequently, such damage occurs to the cardiovascular system (mainly the endothelium and the heart), the autonomic and the central nervous system, lung, kidneys and the neuroimmune network system^4^. However, the precise cellular mechanisms or even the mere level of cellular damages are yet to be measured and understood in a quantitative way. Moreover, several studies indicate that post-COVID symptoms occur in almost two-thirds of the patients even after a non-critical course of the disease^5^. With increasing proportions of infected people suffering from post-COVID and repeated reinfections, and the need to provide further therapies against these, from a translational medical perspective a quantitative cellular harm assay remains a necessity. It is to be established as a routine way to quickly determine robust, standardizable cellular correlates of either SARS-CoV-2 infection or correlates of cellular alterations in putative models of post-COVID syndromes. This “damage assay should be performed quantitatively on relevant models, such as human tissue organoids or certain cell cultures widely used for SARS-CoV-2 studies.

We considered that a common pathway of harm in SARS-CoV-2 infection is cellular calcium ion homeostasis disruption. Since the start of research on SARS-CoV-2 viral entry, multiple mechanisms have been identified. Cellular membrane surface structures such as angiotensin-convertase enzyme 2 (ACE2), neuropilin-1, Ca^2+^-dependent C-type lectins like dendritic cell (DC) or lymphocytic (LC)- specific ICAM-3–grabbing nonintegrin (DC-SIGN and LC-SIGN respectively), cluster of differentiation (CD)-147 protein, Syndecan-1-4, Toll-Like Receptor 4, Limphocyte Function Associated Antigen-1 (LFA-1), sphingolipids, heparan sulphates are the currently known facilitators of different cell entry mechanisms for different SARS-CoV-2 variants with different emphasis^6,7^. It is also known now that multiple different cell types (not just ACE2 expressing ones) are infected via this multitude of entries^8,9^. Interestingly, while some of them already are calcium dependent proteins, most of post-entry events converge in some form of calcium-dependent effector mechanism^10,11^. Two of the most important effectors of harm, cell-to-cell spread via cellular fusion by S2 and fusogenic peptide hexamer formation, or ACE2 downstream action via the calcineurin pathway^12,13^ are both dependent on Ca^2+^ ions. Other results also highlight a connection between calcium homeostasis and SARS-CoV-2 infection.^3,14^ It has been well established previously for many viruses that calcium is essential for viral entry into host cells, viral gene expression, processing of viral proteins, viral maturation and release^14,15^. Thus, these viruses either create their own calcium channel proteins, also known as viroporins, or hijack host-encoded cellular calcium channels to increase intracellular Ca^2+^ ion concentration. This can occur both by calcium influx from the extracellular space and release from the endoplasmic reticulum^15^. Besides the infection effector process, SARS-CoV and also SARS-CoV-2 host cell entry is also shown to be calcium dependent^16^. Both „early” (direct fusion with the cell membrane) and „late” (endocytotic) entry pathways require the presence of Ca^2+^ ion to stabilize the conformation changes of the fusion protein — previously cleaved from S (spike) protein — that is responsible for the fusion of viral and cell membranes^17,18^. Host cell proteases (TMPRSS2, furin etc.) are required to cleave spike protein for the formation of the functional fusogenic elements (fusion peptide, FP) that trigger viral fusion with the host cell membrane^19^. Ca^2+^ binding to the negatively charged residues of FPs seems essential to promote the fusion process. Similarly, high endosomal Ca^2+^ level is necessary in the endocytic vesicles for viral fusion and release of RNA into the cytosol^19^. It has been shown for SARS-CoV, that the viral entry is strictly dependent on the Ca^2+^ ion concentration and is inhibited by calcium chelators such as BAPTA-AM, which act in endosomes^20^. Recent studies with S protein pseudotyped lentiviruses showed that the depletion of extracellular or intracellular Ca^2+^ ion concentration drastically decreased SARS-CoV-2 viral entry^11,21^. Furthermore, although clinical observations are still controversial, based on in vitro studies L-type calcium channel blockers are proposed as potential therapeutics in Covid-19^22,23^. It has also been highlighted that calcium ion plays a crucial role in several steps of viral replication and protein synthesis in the cell, at a molecular level^24,25^. As free calcium concentration is typically low in the cytoplasm (∼100 nM), but it is stored in large amounts in intracellular vesicles, endoplasmatic reticulum (ER) and Golgi lumens (0.1–1 mM)^15^, a Ca^2+^ sensor with prime cell membrane permeability is needed to be able to closely monitor these processes in the living cells. Studies revealed that the viroporins of coronaviruses, such as ORF3a or most notably the E protein, can transport Ca^2+^ ions into the cytoplasm, activate the NLRP3 inflammasome and are involved in virulence^14,26^. These data undoubtedly reveal the role of the calcium ion in the course of COVID-19 as a disease and immunity, too^13^.

Tracking and visualizing the spread of SARS-CoV-2 infection in cellular and organoid systems have been indispensable in several important studies.^27^ In antiviral drug efficacy experiments, real-time quantitative PCR has been the go-to method of infection monitoring^28^. While this method can indicate the overall viral RNA concentration in the extracellular medium, it does not provide any information about the state of individual cells. To visualize the infection in individual cells and thus allow microscopic imaging, fluorescent or immunofluorescent (IF) staining has been applied in cell cultures and organoids^28–30^ However, IF staining requires fixation of the cells which prevents continuous monitoring of the infection process. Furthermore, the meticulous sample preparation and the expensive staining reagents limit its practical applicability. In their interesting approach to develop a novel cell culture system model, Ju et al. replaced the sequence encoding the nucleocapsid (N) protein with that of green fluorescent protein (GFP)^31^. This genetic modification not only decreased the biosafety level (BSL) of the model system but also allowed the fluorescent detection and imaging of the infected cells. Nevertheless, this staining approach involving genetic engineering remains limited to such model systems, and even there, the GFP sequence is deleted from the genome over time as demonstrated by serial passage experiments^31^.

Herein, we report a straightforward downstream-effector-oriented procedure that exploits the disturbance of cellular Ca^2+^ physiology by SARS-CoV-2 through the application of a novel Ca^2+^ sensitive dye. The dye is aimed to be more sensitive and less disturbing to the cellular environments than previously applied ones. Most of the known fluorescent Ca^2+^ ion sensors, especially those with BAPTA chelator functionality^32^, suffer from certain limitations in terms of their cellular applications. First, the widely used acetoxymethyl ester (AM) protected derivatives need to be applied at high concentrations (in 1–2 mM) to generate the active, deprotected form at suitable concentrations for the assay measurements. The deprotection of the AM ester releases a toxic byproduct, formaldehyde, in substantial concentration during its hydrolysis, which may lead to cell damage or death, thus perturbing the cell viability assay^33^. Second, these commonly used dyes are practically membrane-impermeable in their unprotected forms^34^, such as Oregon Green (2’,7’-difluorofluorescein)^34^. To avoid these issues of the AM esters and to find a good compromise between activity and cell internalization, we aimed to design a novel Ca^2+^ ion sensitive sensor molecule, **B**APTA-**E**nabled di**E**thyl-**F**luorescein – **C**alcium **P**robe (BEEF-CP). This compound features two apolar and electron donating ethyl groups on the xanthene moiety, presumably contributing to its internalization and eliminating the need for esters that generate cytotoxic compounds upon activation^33^. Using fluorescence information in a previously-not-achieved high-sensitivity manner, one might be able to collect relevant data about many stages of the infection process. The infection of individual cells and cell cultures indicated by the fluorescent turn-on of the dye is monitored by high-resolution two-photon (2P) microscopy, which is capable of extracting information with high temporal and spatial resolution from different planes of cell cultures or even three-dimensional organoids.

## Results

### The calcium-sensitive fluorescent dye readily enters the cell

BEEF-CP is an analogue of the common Ca^2+^ sensitive dyes, Calcium Green and Oregon Green that features apolar and electron-donating substituents, namely ethyl groups instead of halogen substituents on the fluorescein moiety (Fig. 1a). Fluorescein-based fluorophores are well-known for their non-fluorescent lactone and fluorescent zwitterion tautomerism, which is heavily influenced by the substituents on the xanthene ring, pH and other factors^32^. At the same time, the pH and Ca^2+^ dependent fluorescence of BAPTA-based Ca^2+^ indicators is a key factor in their applicability. At low or high pH values, these dyes can give false negative or false positive signals, respectively. The Ca^2+^ turn-on fluorescence experiments of BEEF-CP demonstrated low background and high enhancement in the biologically relevant pH range of 6.0 8.0, and consequently confirmed its suitability for in vitro measurements (Supplementary Fig. S16).

**Fig. 1:**
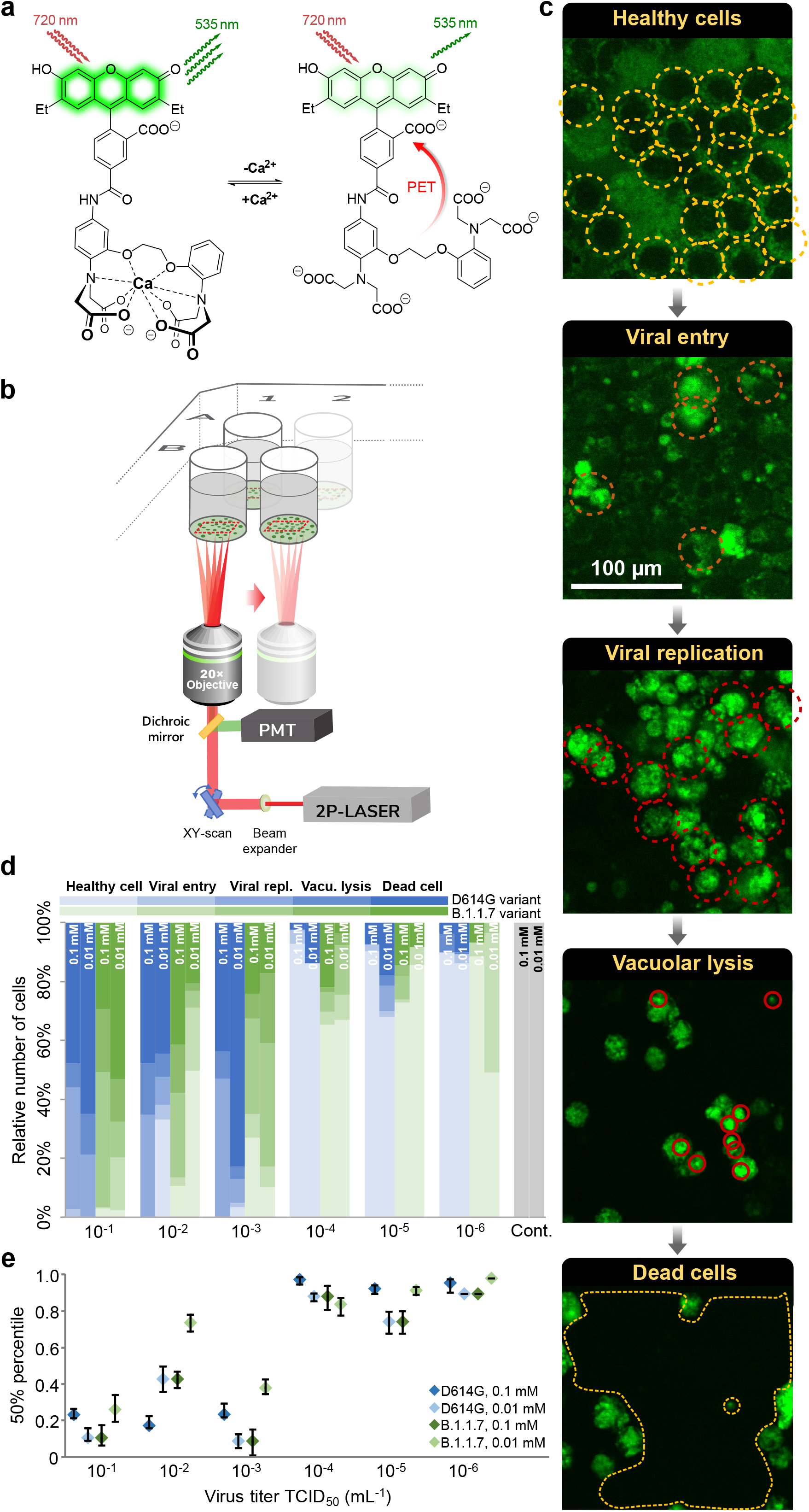
Experimental design and manual analysis of infection in Vero e6 cells using SARS-CoV-2 variants D614G and B.1.1.7. **a,** Fluorescence enhancement of the BEEF-CP dye upon Ca^2+^ binding is attributed to the alleviation of the photoinduced electron transfer (PET) quenching. Its low background fluorescence and high two-photon cross-section (TPCS) along with its advantageous cell internalization, make BEEF-CP an appropriate candidate for cytosolic Ca^2+^ imaging with 2P microscopy. **b,** Experimental design with FemtoViruScope. A pulsed Ti:Sa laser source was used for excitation at 700 nm wavelength, laser light was diffracted with a pair of galvo mirrors for scanning the region of interest. A high-numerical-aperture objective (Olympus 20x) was used to obtain subcellular resolution. Samples were prepared in 96-well bioassay plates and transferred directly to the microscope. Samples were moved semi-automatically to locate the different wells in the field of view of the microscope. **c**, Exemplary images from samples where most cells correspond to a particular stage of the infection: (See Methods for more details on the characterization criteria) **All** *healthy cells*, with no staining in the nucleus, indicated with yellow circles. *Cells in the viral entry phase* have increased signal level and their nuclei show staining. Cells in this stage are indicated with orange circles. *Cell in the viral replication phase*, labeled with red circles, show increased fluorescence level throughout the entire cell. *Cells in the vacuolar lysis phase* show irregular cell shapes as they start degrading. Vacuoles can be distinguished as smaller bright regions with high fluorescence intensities. *Dead cells* are detached from the surface. The number of the dead cells are estimated by the surface region that is not covered with any other cell. **d,** The relative number of cells in different stages of the infection 48 hours post infection for the two variants at two different dye concentration at various virus titer. Data from wells with the same virus titer, variant, and dye concentration are averaged. Statistical data for the individual cell categories are shown in Supplementary Fig. S3. **e,** 50th percentile values, i.e. the percentage of cells under the median level of infection, for the data displayed in panel **d**. Statistical analysis of the manual cell sorting show that i : ratio of healthy cells are significantly higher with lower virus titer, regardless of the virus variant or dye concentration; ii: for high infection levels (TCID_50_ >10^−3^ mL^−1^) the proportion of dead cells is higher for the D614G variant.

The spectroscopic characterization of BEEF-CP involved one-photon (1P) and 2P fluorescent measurements and the quantum yield (QY) determination in the presence and absence of Ca^2+^ ions. The measured fluorescence intensity at pH 7.2 showed a single-step enhancement upon Ca^2+^ addition in accordance with the formation of a single monometallic complex. The fluorescence enhancement was found to be around threefold in the biologically relevant Ca^2+^ concentration range, from 10 nM to 1 M (Supplementary Fig. S15). The dissociation constant of the Ca^2+^ complex (K_D_ = 175.6 nM) is comparable with previously synthesized molecular probes for in vivo calcium level determination. ^35,36^ The background fluorescence of BEEF-CP is not zero in the absence of Ca^2+^, which provides appropriate contrast to visualize even the healthy cells with low level of Ca^2+^ ion during the cell viability assay. Furthermore, the high-slope region of the Ca^2+^-dependent fluorescent curve is adequately positioned to distinguish between healthy and malfunctioning cells.

The further applicability of BEEF-CP in cellular systems was tested by flow cytometry measurements on HEK-293 cell line. Stained cells showed an increased mean fluorescent intensity by around one order of magnitude compared to the control level (Supplementary Fig. S12). In a parallel experiment, the stained HEK cells were treated with ethylene glycol-bis(*β*-aminoethyl ether)-*N,N,N′,N′*-tetraacetic acid (EGTA) prior to the analysis, which resulted in a somewhat lower mean fluorescent intensity. In contrast, cells stained in the presence of ionomycin using otherwise identical procedure exhibit substantially higher mean fluorescent intensity. The ionomycin releases exclusively the intracellular Ca^2+^ storage, which proves that the dye is internalized by the cells as it responds to the cytosolic Ca^2+^ levels.

### Subcellular dye localization is characteristic and reproducibly different in uninfected and infected cell cultures

To perform high-resolution quantitative analysis of the level of SARS-CoV-2 infection we used 2P imaging. This imaging technique has a high, subcellular spatial resolution (∼350 nm), and virtually no out-of-focus signal from non-studied cells or compartments, above or below of the imaging plane, which ensures that the detected signal originates only from the cells of interest. Samples were prepared in a 96-well bio-assay plate with a clear, flat bottom for good optical access. One to three high-resolution, 2P, full-field, 300×300 μm large image were acquired with a FemtoViruScope (Femtonics Ltd, Budapest) at 700 nm wavelength (see Supplementary Fig. S2 for wavelength selection) from one or more representative area within each sample, see Methods for details.

Acquired images were analyzed with two complementary methods (manual and automatic). During manual evaluation, the cells were sorted according to their morphology (Fig. 1b-d), while using the automatic analysis, based on statistical methods, consisted of the classification of the full images based on a set of image parameters (Fig. 2). Both methods take the different features of the acquired raw images into account for the analysis, and for both, we investigated the fine details of the images (at cellular or single-pixel level) for designing a quantitative measure correlating with the initial challenge infection level. Two variants of concern of SARS-CoV-2, namely D614G and B.1.1.7 were studied. These caused pathophysiologically and clinically distinguishable disease patterns in the clinic, so the putative infection-quantitation and infection-assay properties of BEEF-CP would be tested as the distinguishing features of cellular Ca^2+^ related effects of these two variants. Different cellular features such as the state of infection and the apoptotic-like state were visible and distinguishable on the images of Vero E6 cells infected with the two SARS-CoV-2 variants (Fig. 1b). The investigations were done in a phase when still living cells (both healthy and infected) adhered to the plate surface. Cell compartments, such as cell membrane, nucleus, and nucleolus were visible and distinguishable in the samples. With increasing virus titer, the mean fluorescence of infected cells increased significantly as this caused more Ca^2+^ to enter the cytoplasm. When the infecting viral titer increased even further, the morphology of cells changed, and dead cells detached from the wells plate; thus, the number of these cells could be estimated by the ratio of unpopulated areas at the plates’ surface. Due to the subcellular resolution of this cell viability assay, we could detect the formation of several syncytial formations, as giant multinucleated cells in the most infected cultures, which agrees with the earlier observation that SARS-CoV-2 virus causes this type of cellular fusion^37^ (See Supplementary Fig. S11).

**Fig. 2:**
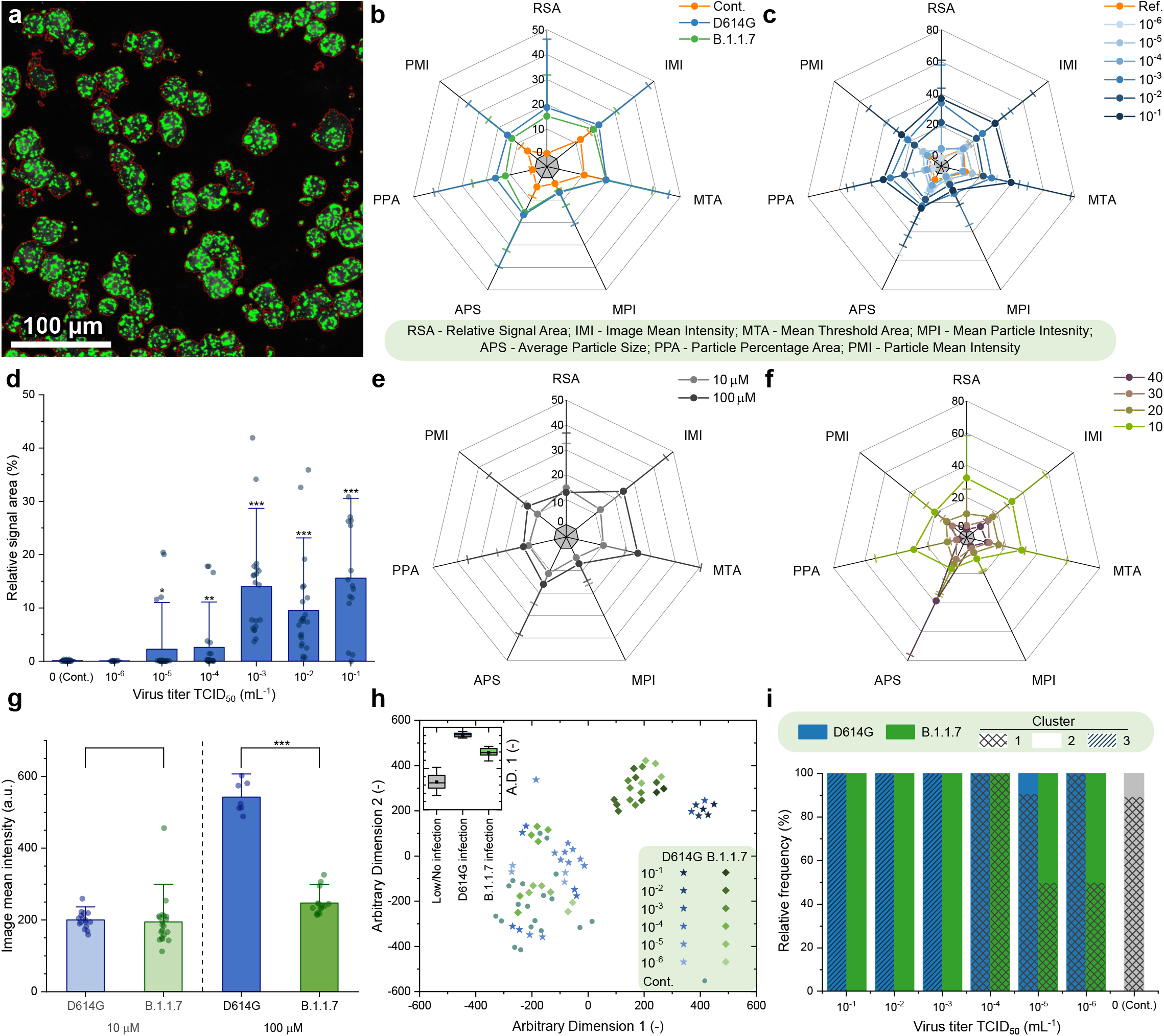
Automatic analysis of the 2P microscopy images of infected cells using SARS-CoV-2 variants D614G and B.1.1.7. **a**, Red contours indicate the areas automatically defined as particles during this analysis. **b,c,e,f**, Spider charts showing the effects of (**b**) variant, (**c**) virus titer, (**e**) dye concentration and (**f**) photomultiplier tube (PMT) relative voltage on the range-normalized values of 7 different image parameters obtained from 2P microscopy images. **d**, The image parameter called ‘relative signal area’ shows a significant increase at virus titers higher than TCID_50_ 10^−3^ mL^−1^. **g**, The image parameter called ‘image mean intensity’ for the two studied variants at high virus titer (TdD_50_ > 10^−3^ mL^−1^) at 10 μΜ and 100 μΜ dye concentration, **h**, T-distributed stochastic neighbor embedding (t-SNE) 2D plot obtained from all the seven image parameters recorded at 100 μΜ dye concentration shows three clusters depending on the virus variant and titer. Inset shows that the three clusters, namely no or low infection, infection with D614G variant, and infection with B.1.1.7 variant are clearly separated along the first dimension. **i**, Classification of the images corresponding to different variants at various virus titer in three clusters by seven-dimensional Gaussian mixture model clustering. (Error bars show standard deviation; significance levels as: * = p ≤ 0.1; ** = p ≤ 0.05; *** = p≤0.01.)

### Manual evaluation

During the manual evaluation cells where manually localized on the recorded images, then the individual cells were visually categorized into five groups, representing different stages of the infection, see Fig. 1c and Methods for more details on cell categories. The individual images from different wells were subjected to a single-blind evaluation by human operator. The cells were classified into four categories (Fig. 1c), while the fifth category, cell death, was determined by the absence of living cells, estimated from the surface area that was not covered with cells, as shown in Fig. 1c. The ratio of the cells belonging to each category was calculated for the different virus variants, virus titer and dye concentration showing an increasing number of cells in the healthy or initial infection categories with the virus titer decreasing (Fig. 1d). The 50th percentile values were calculated for each virus titer by averaging the result from all wells infected with the given titer. For this calculation, cells were sorted by the severity of their infection from healthy to dead cells, and the position of the median was calculated as the ratio of all the cells. The 50th percentile values also show a clear trend with increasing viral load. Furthermore, there is a clear jump in the severity of the infection in the cell culture between TCID_50_ virus titer 10^−4^ and 10^−3^ mL^−1^.

### Automatic analysis

An ImageJ^38^ macro was created to identify particle objects based on an intensity threshold (Fig. 2a), and to extract image data from the fluorescence microscopy images in the form of a few numerical parameters. The ImageJ macro is available in the Supplementary Information. Seven parameters characterizing the detected particles were defined in each image, more specifically i: Relative signal area, ii: Image mean intensity, iii: Mean of the threshold area, iv: Maximum particle intensity, v: Average particle size, vi: Particle percentage area, and vii: Particle mean intensity.

The input images correspond to experimental points that can be described by two infection-related variables, namely virus titer and virus variant and by two measurement-related variables, namely BEEF-CP sensor concentration and photomultiplier tube (PMT) voltage. Analysis of the image parameters as a function of the infection-related and measurement-related variables can yield information about the efficiency and robustness of the automatic image analysis, respectively. Fig. 2b–c, 2e–f show that the image parameters vary according to different acquisition settings, i.e., virus concentration, virus type, dye concentration and PMT voltage. Importantly, certain image parameters such as ‘Relative signal area’ (Fig. 2d) can be a good indicator of the viral infection, as it shows increasing parameter value with increasing virus titer. However, images of the two variants did not show clear separation by any parameter values without eliminating the variation caused by the measurement-related variables.

Other prevalent Ca^2+^ sensors, such as OGB-1 AM (Oregon Green 488 BAPTA-AM), are used at a high concentration, around 1 mM (ranging 0.9-1.4 mM^39–42^). We used BEEF-CP at lower concentrations, at 0.1 mM and 0.01 mM. Both concentrations were suitable to differentiate between little or no infection and a high level of infection using automatic image analysis. Moreover, differentiation of the variants responsible for the infection became possible when the images obtained with higher dye concentration (0.1 mM) were analyzed in those cases when the well was highly infected (TCID_50_ > 10^−3^ mL^−1^) (Fig. 2g). Therefore, for further analysis, only those images were considered that were acquired from samples stained with 0.1 mM dye concentration.

A cluster analysis that considers all seven parameters was run on all the images acquired using 0.1 mM dye concentration. Two-dimensional visualization of the seven-dimensional clustering (Fig. 2h) shows three separated clusters; see Methods for the detailed analysis. A three-component gaussian mixture model cluster analysis was performed on the image parameters. Fig. 2i shows that the three obtained clusters agree well with the three distinct states of the wells, i.e. little or no infection, highly infected with the D614G virus variant, and highly infected with the B.1.1.7 variant.

### Quantitative image analysis shows a viral variant-dependent separation and a clear infection threshold of the applied virus titers

The manual and automatic analyses both showed a difference between the two tested virus variants, with D614G having a significantly higher infection rate. This finding complements earlier results that found a higher infection rate for D614G than B.1.1.7 in HeLa ACE2 and similar rates in HEK-293T cells^43^. There is a very clear threshold found over 10^−3^ mL^−1^ TCID_50_ virus titer, where instead of occasional virus infection spread, we saw reliably escalated infection at all imaging sites. Furthermore, the automatic analysis revealed other novel findings. First, it showed a clear threshold for the dye titer, showing that at least 0.1 mM dye concentration is required for the significant differentiation between variants (Fig. 2g). The results suggest that the fluorescent intensity of infected cells — related to the image mean intensity — is delimited by the dye concentration at 0.01 mM and by the Ca^2+^ concentration at 0.1 mM dye load. Second, the cluster analysis of the data could distinguish three separate groups. Among these was the non-infected group, the images falling into this cluster showed similar characteristics to the control measurements. However, more interestingly, the remaining images were almost perfectly separated by the virus variant used for the infection (Fig. 2h). These results demonstrate that this method is capable of quantifying the infection level and can also differentiate the two applied variants, D614G and B.1.1.7.

## Discussion

This study presents a general concept of quantitative calcium-based viral cellular harm assay, here applied on SARS-CoV-2 viral variants. For quantitative analysis of SARS-CoV-2 infection, we chose to monitor the Ca^2+^-dependent cellular mechanism previously described in the literature as it strongly correlates to the infection rate. For monitoring the intrinsic concentration of Ca^2+^, a novel cell membrane permeable internalized, two-photon active Ca^2+^ selective sensor molecule was designed and synthesized, which allows the measurement of both extra and intracellular Ca^2+^ concentration, without chemical compromises. BEEF-CP as a Ca^2+^-selective sensor exhibits several advantages for practical applicability, such as i) appropriate cell internalization^44^; ii) the effective concentration of the dye is 10-100 times lower (0.01 0.1 mM) compared to other widely used compounds (>1 mM); and iii) is free from toxic leaving groups hence infection processes are not affected by the chelation of Ca^2+^ using the novel dye, unlike in the case of working concentrations for BAPTA-AM; iv) appropriate spectrophysical characteristics and sufficient pH insensitivity around the physiological pH.

ViruScope has provided high resolution and extreme sensitivity by fast 2P fluorescence imaging. The selection of the 2P imaging method was indicated by its fine z-sectioning ability compared to single-photon imaging, as shown in Supplementary Fig. S1. As we used the fluorescent intensity profile of the images acquired for quantitative analysis, it was imperative to measure only a single cellular layer without contamination from the out-of-focus cells or compartments. This was achieved by the means of 2P imaging. Furthermore, it should be noted that the proposed imaging method can be directly applied to tissue slices or organoids, opening new ways to test the effects of SARS-CoV-2 infections in more complex systems as well. The high-resolution detection of the Ca^2+^ level within the infected cells requires a sensor molecule with appropriate 2P sensitivity and adequate cell internalization. The developed sensor, BEEF-CP meets these criteria owing to its diethylfluorescein fluorophore. The fluorescent dye concentration influences the information content of the images; 0.1 mM — unlike 0.01 mM — proved to be sufficient to determine not only the level of infection but also the virus variant. Importantly, neither applied dye concentrations induced cell death in the control wells. The incubation time (60 minutes) was sufficiently short for rapid imaging of the fast dynamics of the infection.

Two different methods, with comparable results, have been used to evaluate imaging data. The first method allowed, by manual cell counting, to quantify the number of cells in different stages of the infection such as i) healthy cells, ii) initial infection of the cells, iii) infected cells, iv) infected vacuoles, v) dead cells (as the absence of cells). The second method automatically extracted image parameters from the image files to compare the different infection states without needing manual categorization. With this methodology that works in BSL3 environment, we demonstrated that the increase of the intracellular Ca^2+^ ion concentration is a good indicator of the infection progress as the concentration of the calcium changes significantly inside the cell during virus infection. Correlates of infection in relationship to intracellular and fusogenicity-related calcium concentration changes might offer a prospective and reliable way of infection and therapy monitoring in vitro.

In conclusion, we propose a new calcium-sensitive dye for correlates of SARS-CoV-2 infection determination, quantification and monitoring in two- and three-dimensional cell cultures using 2P microscopy. This molecule is at least one order of magnitude more sensitive than previously applied calcium dyes. Our BSL-3 2P microscopy measurements have also established that the applied concentrations of the dye do not interfere with viral replication and viral fusion events. We postulate that this method thus ensures proper monitoring of the full viral entry spectrum of events. At the same time, it also enabled us to distinguish intracellular details of cell damage, such as vacuole and apoptotic body formation. Using clustering analysis, we could use the 2P microscopy calcium fluorescence images for the distinction of two different viral variants in cell cultures. Using this 2P microscopy method, we could also establish correlates of infection related to the initial viral multiplicity of infection numbers. Further research may utilize this technique to study and compare the effects of antiviral drug candidates on the infection in cell cultures.

## Methods

### Two-photon (2P) imaging

All experiments were performed on a FemtoViruScope (Femtonics Ltd., Budapest, Hungary). Laser pulses were generated by a Mai Tai HP laser (SpectraPhysics, Santa Clara, CA), the laser wavelength was tuned between 750 and 850 nm, depending on the experiment. Laser intensity was controlled using a Pockels-cell (PC) electro-optical modulator (model 350–80 LA; Conoptics) after beam expansion (1:2, Thorlabs). For excitation and signal collection, a XLUMPLFLN20XW lens (Olympus, 20x, NA 1.0) was used, separated using a dichroic mirror (700dcxru, Chroma Technology) before the two-channel detector unit, which was sitting on the objective arm (travelling detector system) as described in detail elsewhere^16,45^. Emitted light entering to the detectors were filtered with emission filters, ET520/60 for the green and ET605/70 for the red channel (Chroma Technology, Bellow Falls,VT). Fluorescence signal was collected to GaAsP photomultiplier tubes (PMT) fixed on the objective arm (H7422P-40-MOD, Hamamatsu). A set galvanometric mirrors was used to acquire full field images. All acquired images were recorded at 1000×1000 pixels resolution, spatial resolution was 0.36 μm/pixel. Single pixel integration time was 6.67 μs, resulting a 6.7 s image acquisition time. The PMT’s maximal amplification voltage (∼300 V) and the imaging laser power (∼50 mW) were kept constant through all measurement and for all wavelength for a more reliable comparison between the acquired fluorescent images. Dynamic range of the PMT is 16bit, we selected the PMT’s gain to obtain most of the range but have a maximum of 50% of the pixels dark on the dimmest and no saturated pixel on the brightest image.

We acquired a series of images using different stimulation wavelengths to calibrate the optimal laser wavelength. To compensate for the change of the output laser power experienced when tuning the output of the pulse laser, the PC amplification was aligned individual at every different wavelength tested, to keep to laser intensity reaching the sample constant at ∼50mW power. The amplification levels were calibrated prior to the acquisition with an intensity meter placed under the objective. Note, that the transmission of the objective is also dependent on the laser intensity; i.e. measuring the intensity under the objective also eliminates this error, besides the laser output modulation. Furthermore, not just the transmission but the material dispersion of the objective is wavelength dependent. Temporarily elongated laser impulses can also affect the 2P efficacy. To minimize this effect, we used glass stand chamber well plate, where the effect of the material dispersion is minimal. The maximum calculated elongation of the pulse, caused by the material dispersion, when tuning the laser from 750 to 850 nm is 19 fs which. Due to the quadratic nature of the 2P power on the excitation, and because the full pulse length is still below 200 fs at 850 nm, pulse elongation has less than 1% effect on the stimulation quantum efficacy. This is much lower than the effects we demonstrated and falls within its noise level, therefore we took no count for this effect during the analysis of the wavelength optimization.

One to three high-resolution image is acquired from every well. The stage was moved between the plates automatically, with the know horizontal and vertical distances between the wells; however, the field of view and the focus were selected manually. Images were saved as multichannel tiffs (for the dual wavelength detection). Image acquisition was performed with a custom Matlab code.

### Virus categories during manual analysis

Imaged cells were manually localized on the recorded images, and then individual cells were visually categorized into five groups according to the following criteria:

a. <u>Healthy</u>: Healthy cells display a pale, low level fluorescence, due to low Ca^2+^ level and their shape is regular. The distribution of cells is homogenous, their arrangement on the plate is normal.
b. <u>Viral entry</u> : These cells clearly show bright spot(s) in certain areas localized near the membrane, indicating an initial infection accompanied by the increase of the intracell Ca^2+^ level.
c. <u>Viral replication:</u> These cells exhibit bright fluorescence, due to the increased Ca^2+^ level and signs of irregularities in their whole shape, which remains compact. Their mean brightness is at least 4-times higher compared to healthy cells.
d. <u>Vacuolar lysis</u>: Dying cells displaying bright fluorescence in their integrity and showing obvious disintegration were selected in this category. Due to the apoptosis-like disintegration of cells, bright fragments appear nearby the original cell, which were considered part of the initial cell and not counted separately.
e. <u>Dead cells</u>: Frames in samples with high viral titer contained significantly more dark areas. Cells disappeared from the spots where they were initially cultured. The difference between the mean cell count in non-infected frames and the actual cell count of the frame was considered the number of dead cells.

Cells belonging into each category were counted manually by the operator performing the analysis without direct information about the image position on the well-plate, i.e. without information about the virus titer, variant or dye concentration corresponding to the image.

### Automatic analysis

An ImageJ^38^ macro was created to load PMT normalized images sequentially, run a despeckle filter to remove noise, run a threshold command to select the signal, measure the mean, max and relative area of the selection, measure the mean intensity of the whole image (including empty space) and finally create a selection based on the threshold. To quantitate particle texture and roughness, a copy was made, and a gaussian blur was applied. The converted image set was then subtracted from the original to enhance any variations in the image. Threshold areas in these images were then selected as before to create a second mask which is again analyzed. Seven parameters characterizing the detected particles on the images were defined for each image, more specifically i: Relative signal area, ii: Image mean intensity, iii: Mean of the threshold area, iv: Maximum particle intensity, v: Average particle size, vi: Particle percentage Area, and vii: Particle mean intensity. T-distributed stochastic neighbor embedding (t-SNE) 2D plot was obtained in MATLAB using the built-in “tsne” function with default random number generation, a perplexity value of 10 and exaggeration value of 50 on all seven parameters of images acquired with 0.1 mM dye concentration. Gaussian mixture model and k-means clustering analyses were performed in the seven-dimensional space determined by the seven image parameters. The number of clusters was set to 3. For the gaussian mixture fitting, the regularization parameter value was set to 1.

## Data Availability

The authors declare that all the data supporting the findings of this study are available within the paper and Supplementary Information. Raw microscopy images are available as Source Data. Additional raw data are available from the corresponding authors upon request.

## Author Contributions

DM, GS, ZK, BP, GF, NK, PH, ÁZ, KK carried out the biological tasks and microscopy; GS, GF, LC performed the data analysis; EK and ZM were responsible for the chemical synthesis and characterization; AF carried out the TPCS measurement; AC performed UV-Vis and fluorescent spectroscopy; ÁS carried out the flow cytometry measurements; EK, DM, BR supervised the research project; GK, BR provided the technical background for 2P imaging; DM, GS, LC, MZ and EK wrote, reviewed and edited the manuscript.

## Acknowledgements

We thank Balàzs Chiovini, Linda Sulcz-Judák and Imre G. Csizmadia for critical evaluation of the manuscript. The authors are grateful to Miklós Dékány (Richter Gedeon Pic.) for his help with HRMS and MS-MS measurements. DM thanks Zoltán Kaleta for his valuable suggestions. Miklós Sz. Kellermayer is gratefully acknowledged for scientific guidance and for setting up the physical virology research line at the Department of Biophysics and Radiation Biology, Semmelweis University.

## Supplementary Information

Experimental procedures, spectral data, NMR spectra, supplementary figures of biological investigations, numerical data corresponding to the presented graphs are provided as Supplementary Information.

## Supplementary Information for

### Supplementary Methods

#### Viral isolate and viral infection gradient plate

D614G mutant variant of the ancestral Wuhan-Hu-1 SARS-CoV-2 virus, isolated from a Hungarian patient in 4 July 2020. SARS-CoV-2 B1.1.7 (Alpha) variant isolated from a Hungarian patient in October 2020. All experiments with the application of infective material were performed under Biosafety Level 3 (BSL-3) conditions at the National Biosafety Laboratory, National Public Health Center (Budapest, Hungary). One day prior to infection Vero E6 cells were plated in 96-well flat bottom tissue culture plates (TPP, Switzerland) in vaccine production-serum free medium (VP-SFM) until each well reached 80% confluence.

The Vero E6 cells for isolation were maintained in DMEM (Lonza, Switzerland) medium supplemented with 5% OptiClone fetal bovine serum (FBS, Euroclone, Italy) and Cell Culture Guard (PanReacApplichem, Germany). The viral titers were determined by the median tissue culture infectious dose assay (TCID50) method and calculated using the public domain code TCID_50_ calculator v2.1 [Binder M. TCID50 Calculator (v2.1-20-01-2017_MB) [(accessed on 10 June 2020)]; available online: https://www.klinikum.uni-heidelberg.de/fileadmin/inst_hygiene/molekulare_virologie/Downloads/TCID50_calculator_v2_17-01-20_MB.xlsx].

The basic viral titer of the stock solution was calculated as lx 10^−1^ ml^−1^. Subsequently, a tenfold serial dilution was performed using the stock solution, i.e. from 10^−2^ until 10^−6^ TCID_50_, each in a final volume of 100 microliters. This volume was supplemented to a final volume of 200 microliters with VP-SFM and TCID_50_ added to the plate wells for the viral infection calcium imaging tests in triplicate. A positive control triplicate using 10^−1^ TCID_50_ and a negative uninfected control triplicate were also plated.

The plates were incubated in a humidified 37 °C incubator at 5% CO_2_ for forty-eight hours to allow for the cytopathogenic effects of viral replication to complete.

For imaging, the fluorescent dye solution was added to each well in 10 microliters of volume in a solution of 10 or 100 times diluted original stock (1 mg of dye dissolved in 1 mL of ethanol, which was diluted with 9 mL of distilled water). The supernatant replaced after 60 minutes incubation prior to the imaging.

#### Flow cytometry measurements

The flow cytometry measurement was carried out on a BD-FACSVerse instrument (BD company) including three laser sources and 8 channel in FICS channel. The HEK-293 cell line was implemented according to the generic vendor protocol at 37 °C in humidified atmosphere with 5% CO_2_. The base medium for this cell line is D-MEM with 4.5 g L^1^ glucose. To make the complete growth medium, fetal bovine serum was added to a final concentration of 10%.

For calcium efflux measurement we used a cell suspension with 1 × 10^6^ cell in 1 mL HBSS (Thermo Fisher) solution. Cells were loaded with BEEF-CP dye final concentration 10 μΜ or without dye at HBSS as untreated control. Cells were incubated for 45 minutes at 37 °C in a shaking incubator (500rpm) in the dark. Cells were washed twice with DMEM with 2% FBS and resuspended in HBSS. For recovering, the cells were stored in the dark at room temperature until about 0.5–1 hour. The samples were measured with a 3 laser, 8 channel BD FACSVerse instrument.

The baseline fluorescence was determined with untreated cells. For maximal calcium flux ratio, cells were treated with 1 gg/ml ionomycin (Thermo Fisher) for 5 minutes and measured. For negative control cells were inhibited with EGTA. Positive control: Ionomycin, 1 gg mL^−1^ final concentration. Negative control: EGTA 8 mM final concentration. The data were analyzed with BD FACSuite Software.

### Supplementary Figures

**Figure S1.**
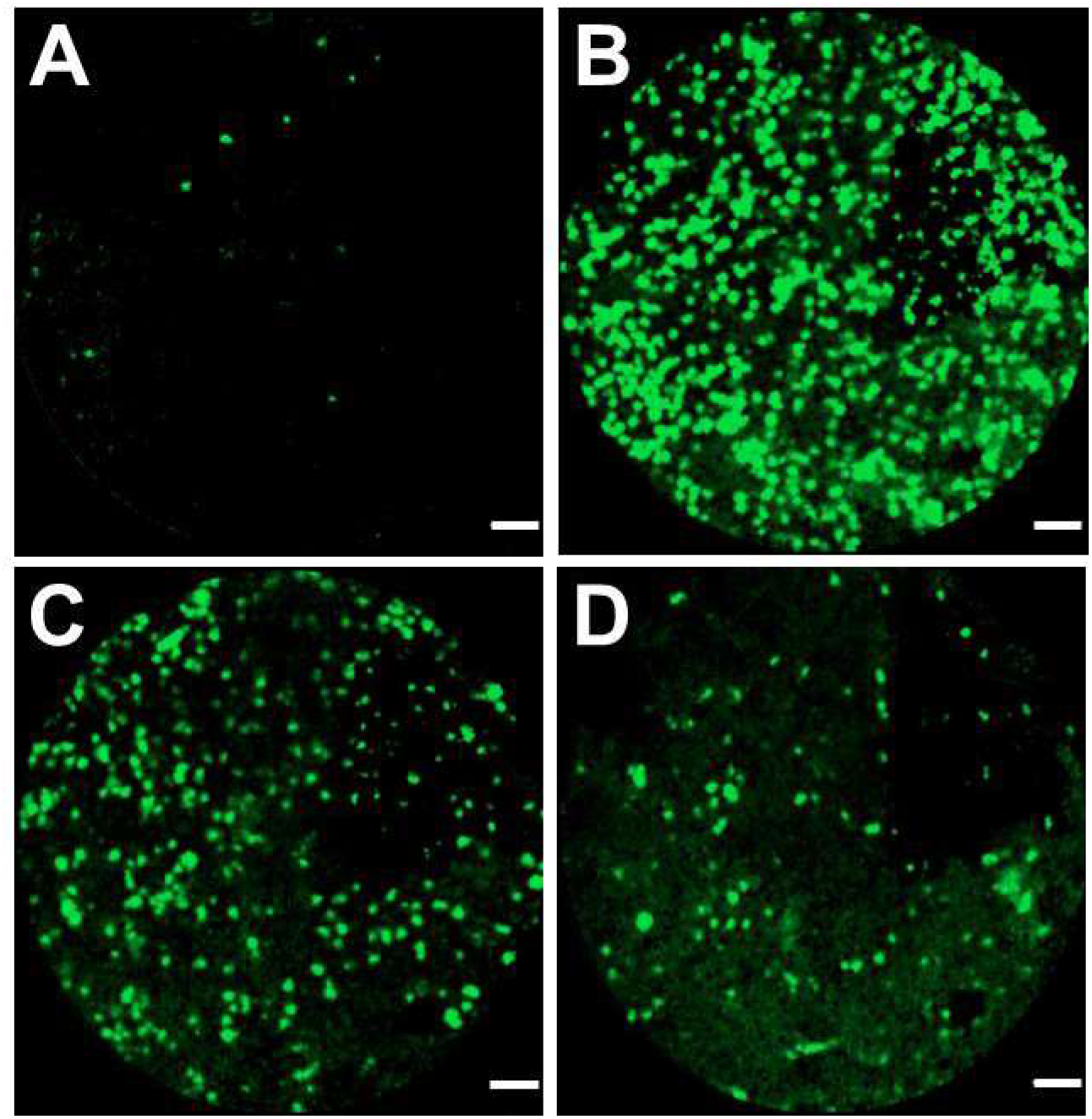
**Images acquired with a Cellvizio confocal laser scanning fiber optic endomicroscope** using the 488 nm excitation. Panels (A) and (B) show the cellular calcium response in bright green fluorescence using the novel calcium sensing dye, while (C) and (D) show the use of the dye in normal non-infected cell culture. Besides the obvious viral titer dependency of the readout calcium light response, the depth dependent changes in resolution and cellular image blurring are also emphasized when this non-3D method of imaging is applied in real-life 3D cell culture conditions. Scale bars 50 gm.

**Figure S2.**
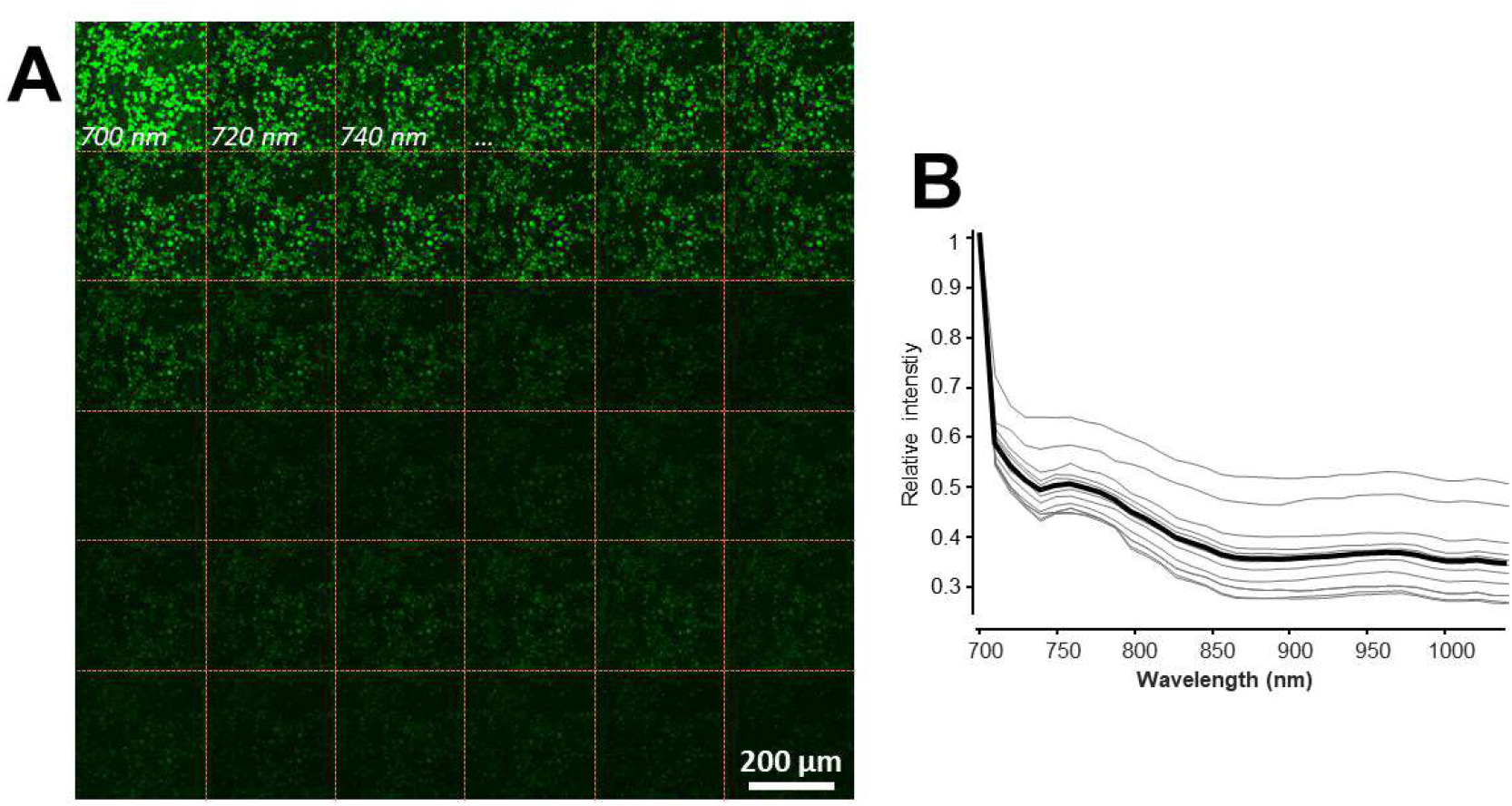
**Wavelength selection.** (A) ‘Wavelength stack’, images acquired from the same imaging region without changing the laser intensity or PMTs’ setting. (B) Averaged two-photon fluorescent signal as a factor of the wavelength, as in (A). Data was pulled from hand selected image areas contains distinguishable structures. Gray lines indicate individual measurements, from different well-plates, intensity is normalized to the one measured at 700 nm. Black line indicates average of 10 measurements with a clear stimulation peak at 700 nm. This 700 nm excitation wavelength was used for all further measurements.

**Figure S3.**
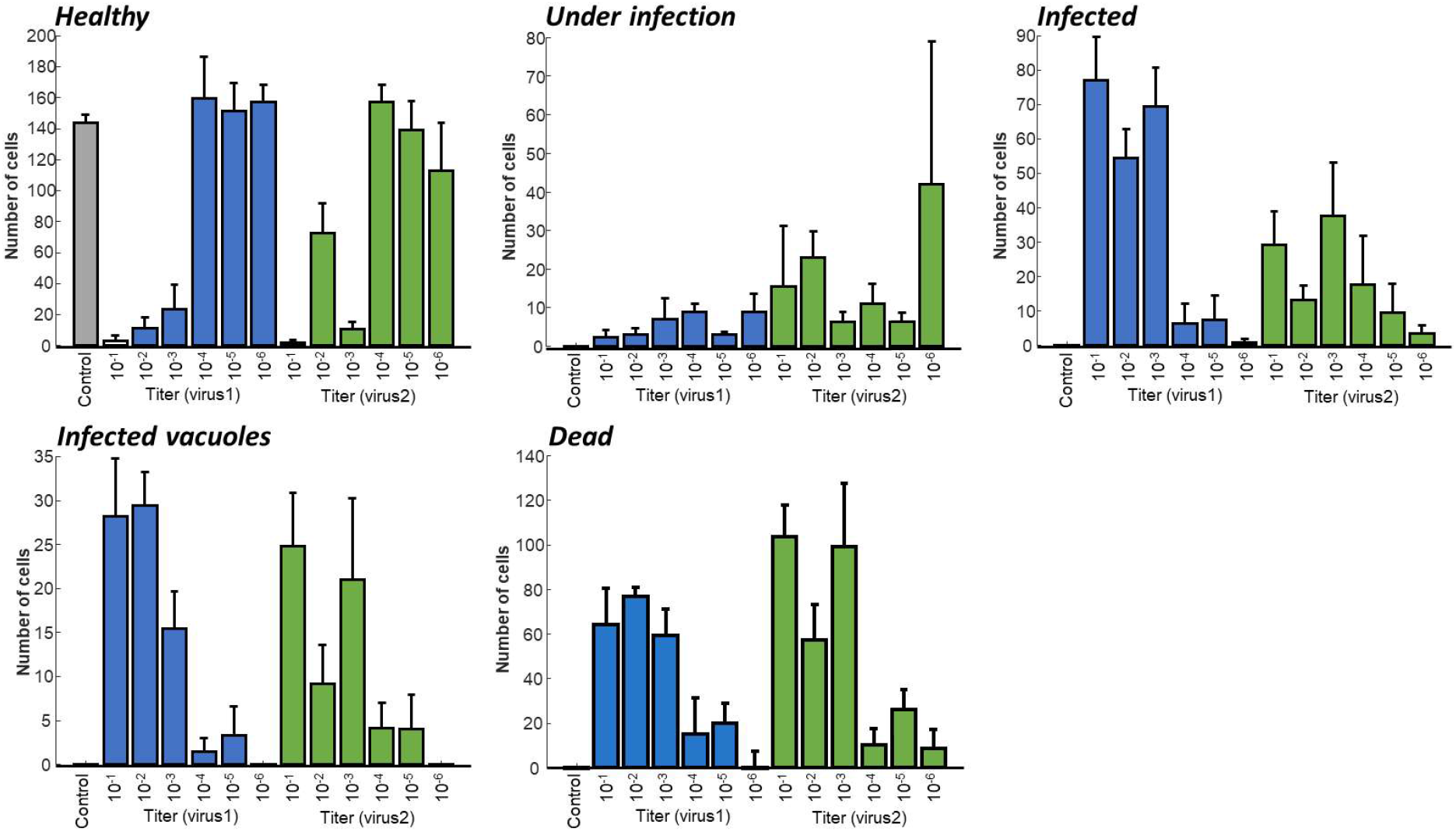
**Distribution of cells in different infection state as the function of the virus titer.** All graphs show the average number of cells found in the different cell categories per image. If multiple images were taken with the same condition the data is averaged. Error bars show ± S.E.M.

**Figure S4.**
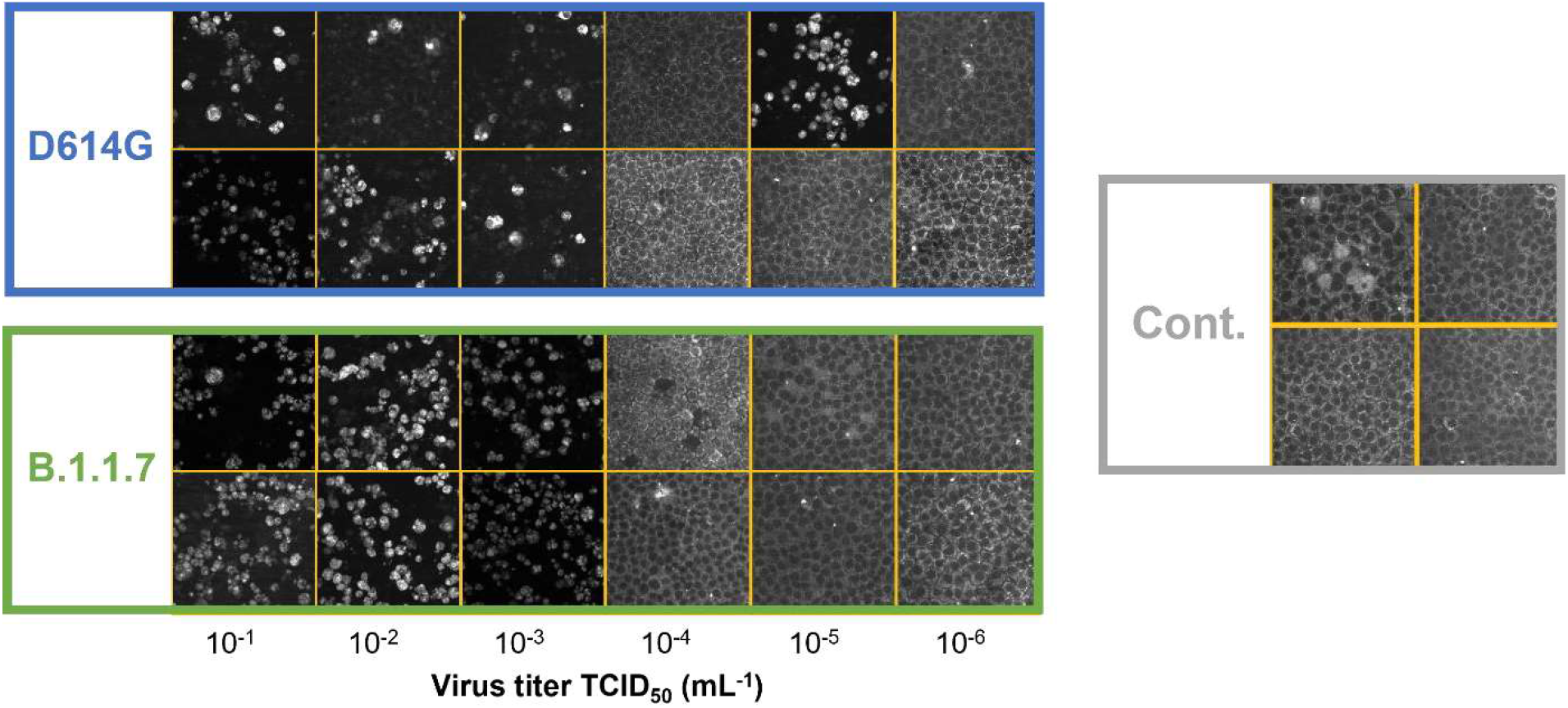
**Exemplary images for the different level infection.** Representative images about how the applied virus titer influences the appearance of the labeled cell culture during two-photon imaging. Blue square shows the samples labeled with the D614G variant, green square for the B.1.1.7 variant. Gray square shows the Control images.

**Figure S5.**
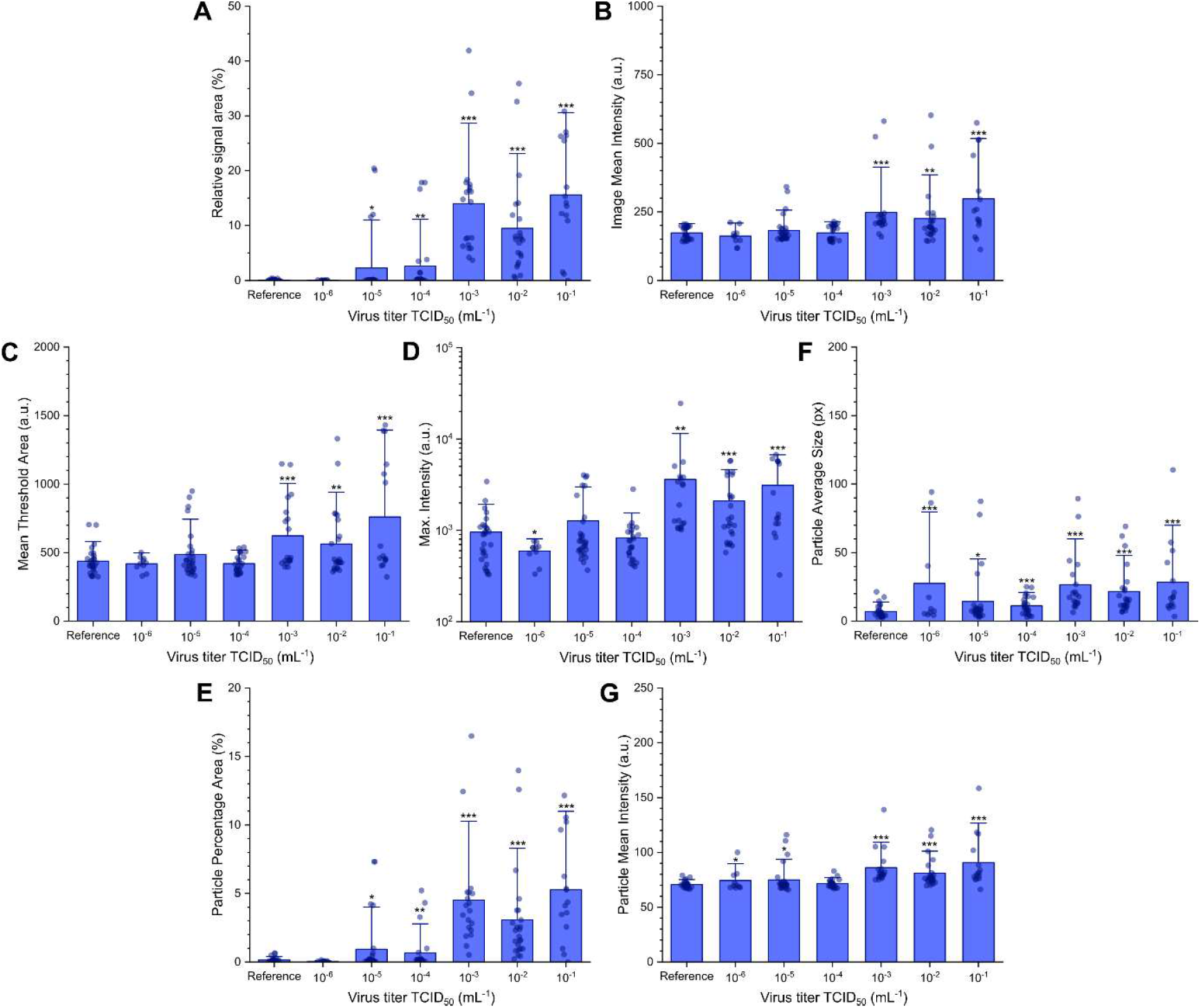
**Image parameters from the automatic image analysis plotted against virus titer (TCID_50_).** The automatic analysis determined seven different parameters for each image. Dots represent the measurements from the individual images, while blue bars show the averages for the given measurement conditions. (Error bars show standard deviation; significance levels as: * = p ≤ 0.1; ** = p ≤ 0.05; *** = p ≤ 0.01).

**Figure S6.**
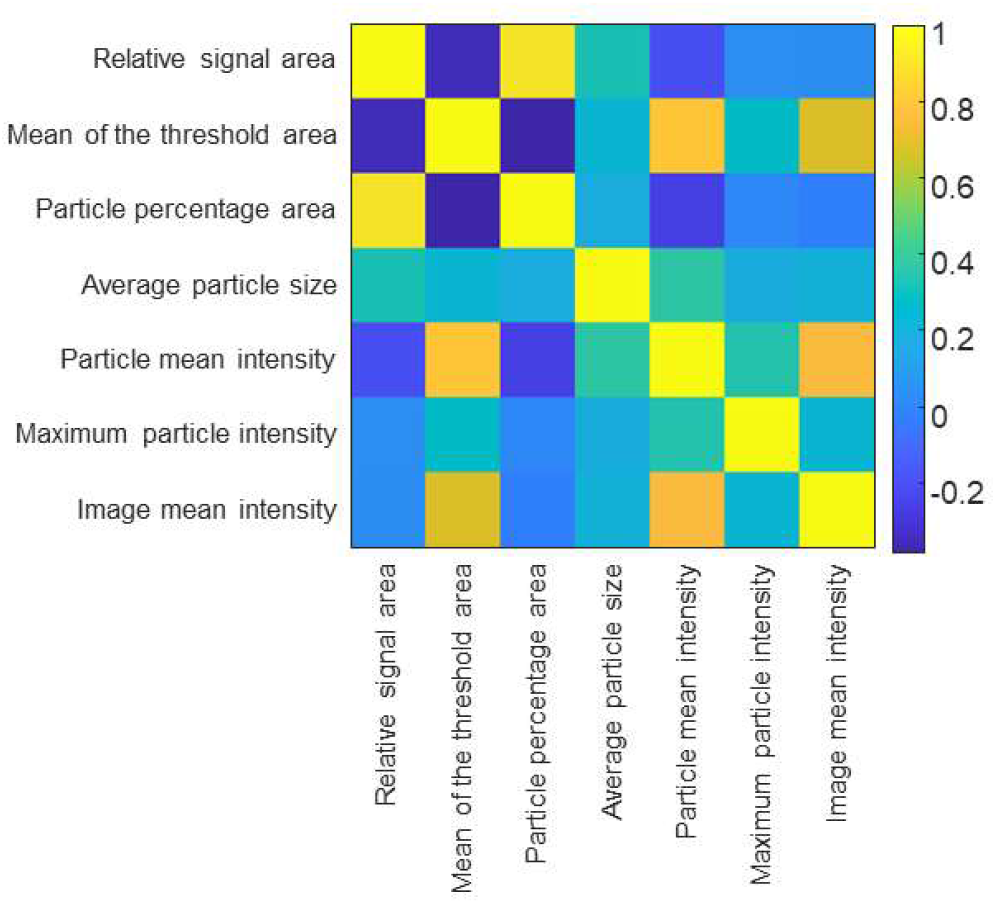
**Cross-correlation between the parameters.** Cross-correlation between all the seven parameters. None of the correlation coefficient besides the center axes was higher than 0.8, therefore all parameters could be used during the cluster analysis.

**Figure S7.**
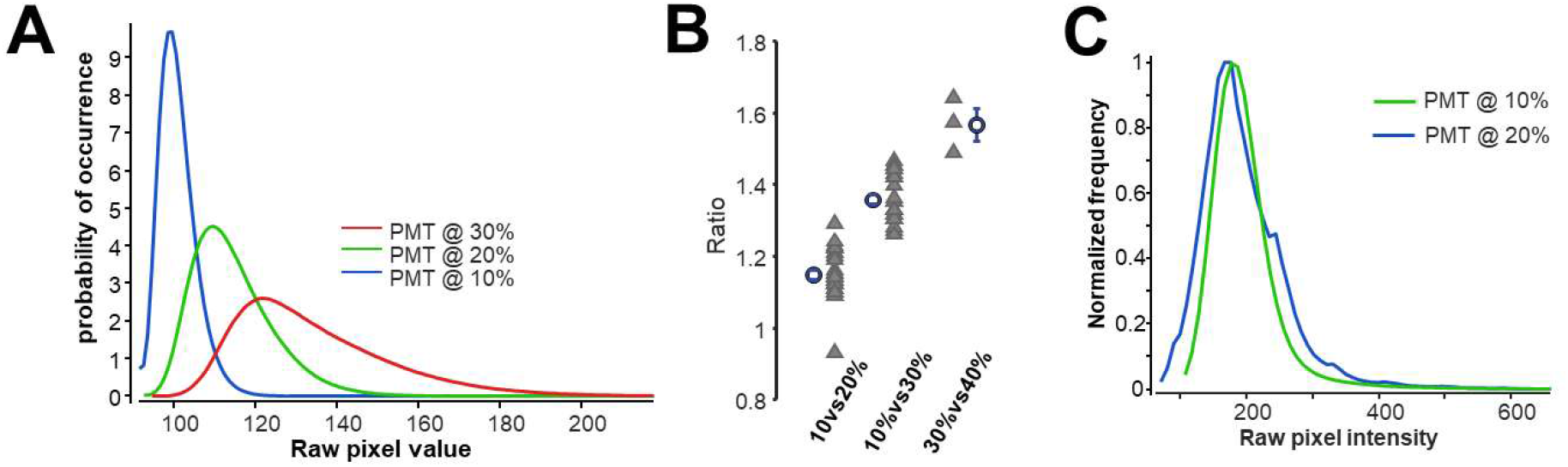
**Normalization for different PMT voltages.** (A) Exemplified pixel intensity histograms for an imaging region that have measured with three different PMT voltage. For comparison between images measured at different PMT voltage, images at 20% and 30% are normalized to the level of the images which were measured. For this normalization an offset and a division have been used. The offset and division coefficient were determined so that the curves have similar mean and deviation. Coefficient from multiple image duplets and triplets were averaged. This average used for all the images. (B) Division ratios for the different PMT voltage differences. Triangles shows the individual calculations while circles show the averages. Error bars represented as S.E.M. (C) Show of an example of for the normalized histogram. Curves drawn from the images acquired with virus titer of 10^−2^ mL^−1^ TCID_50_ for the B.1.1.7 variant. For 10% PMT case data is pulled from the original figure while the image acquires at 20% PMT was normalized as above before the histogram is calculated.

**Figure S8.**
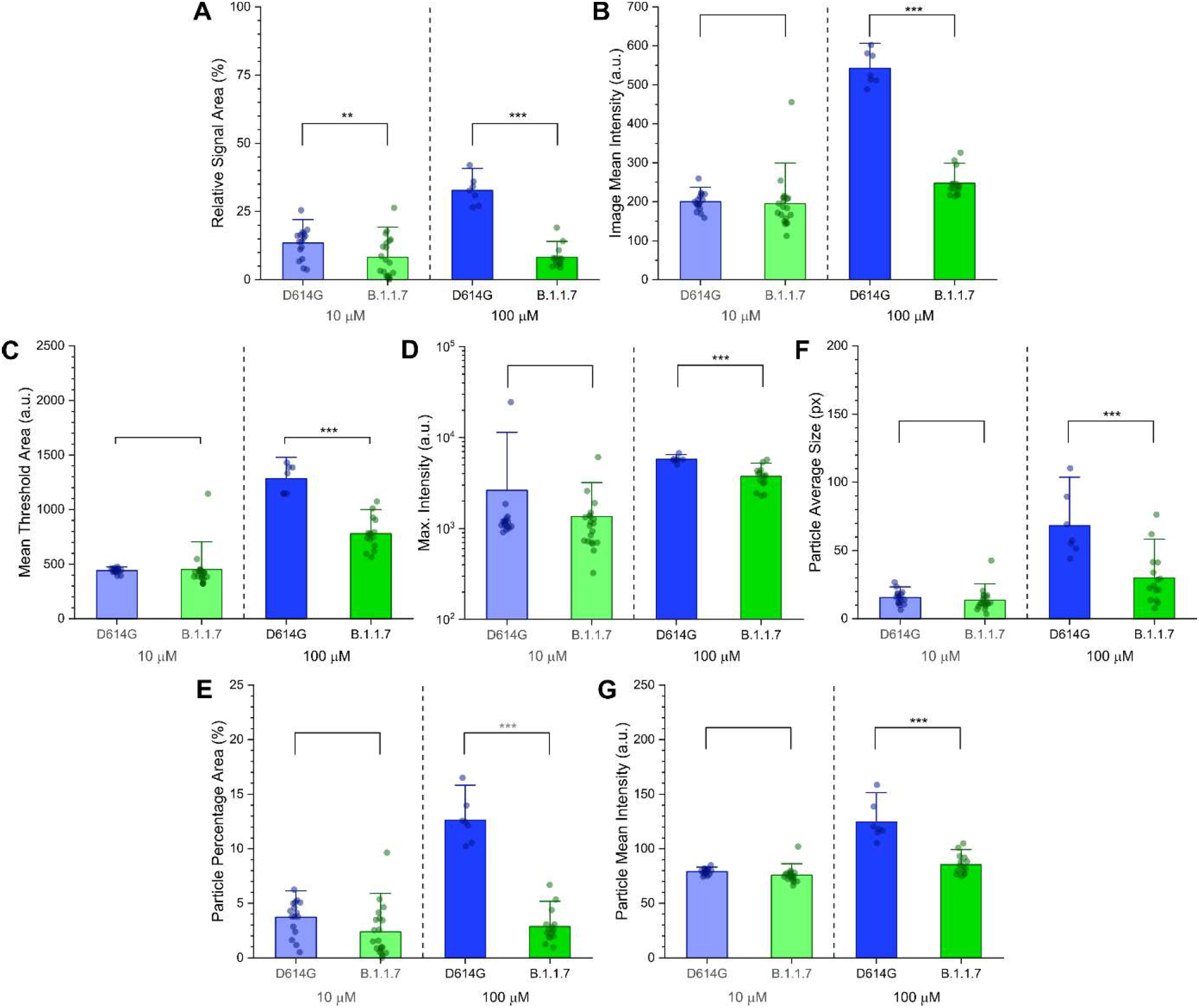
**Image parameters from the automatic image analysis plotted against the virus variant** at different dye concentrations (i.e. 10 μΜ and 100 μΜ). Only data points corresponding to virus titer of IO^−3^ mL^−1^ TCID_50_ and higher were considered. Dots represent the measurements from the individual images, while bars show the averages for the given measurement conditions. (Error bars show standard deviation; significance levels as: * = p ≤ 0.1; ** = p ≤ 0.05; *** = p ≤ 0.01).

**Figure S9.**
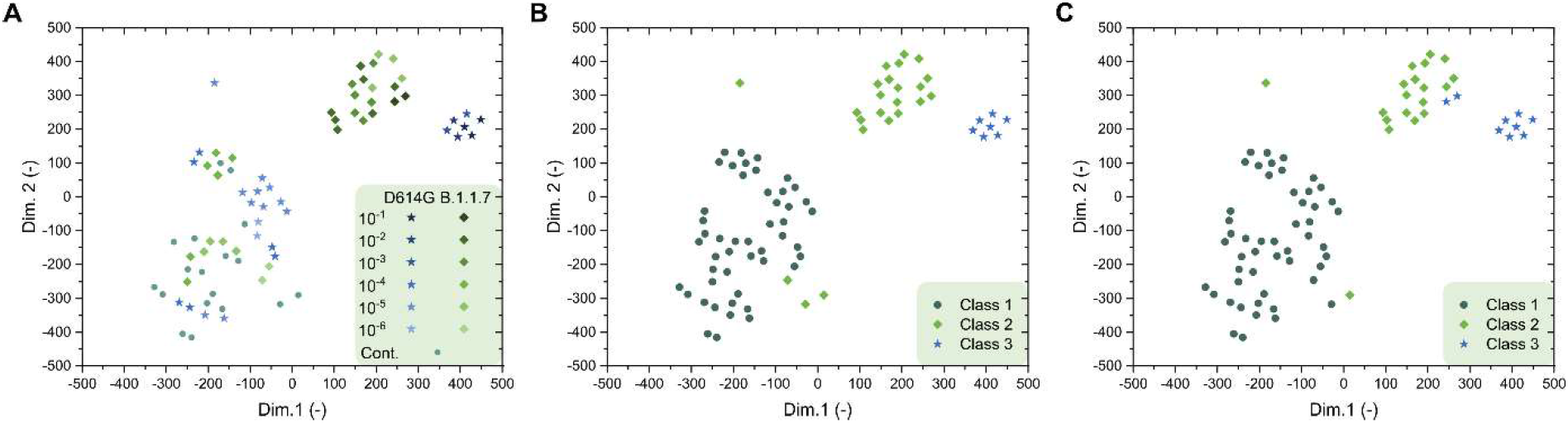
**T-distributed stochastic neighbor embedding (tsne) 2D** plots obtained from all the seven image parameters recorded at 100 μΜ dye concentration. (A) Tsne plot with virus titer TCID_50_ (ml^−1^) values and virus variant for each point. (B) Tsne plot showing the three classes established by seven-dimensional gaussian mixture model clustering. (C) Tsne plot showing the three classes established by seven-dimensional k-means clustering. For both clustering methods, the number of clusters was set to three.

**Figure S10.**
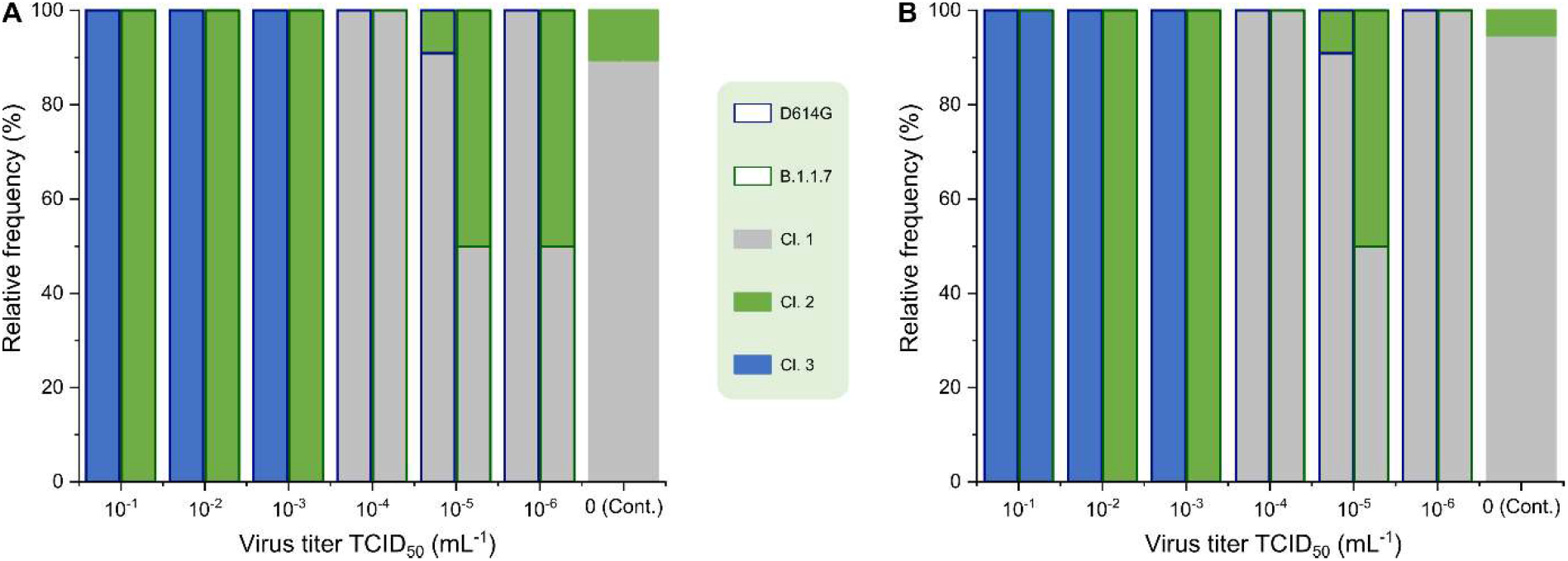
**Classification of the images corresponding to different variants at various virus titer in three clusters by seven-dimensional clustering.** (A) Gaussian mixture model clustering. (B) K-means clustering.

**Figure S11.**
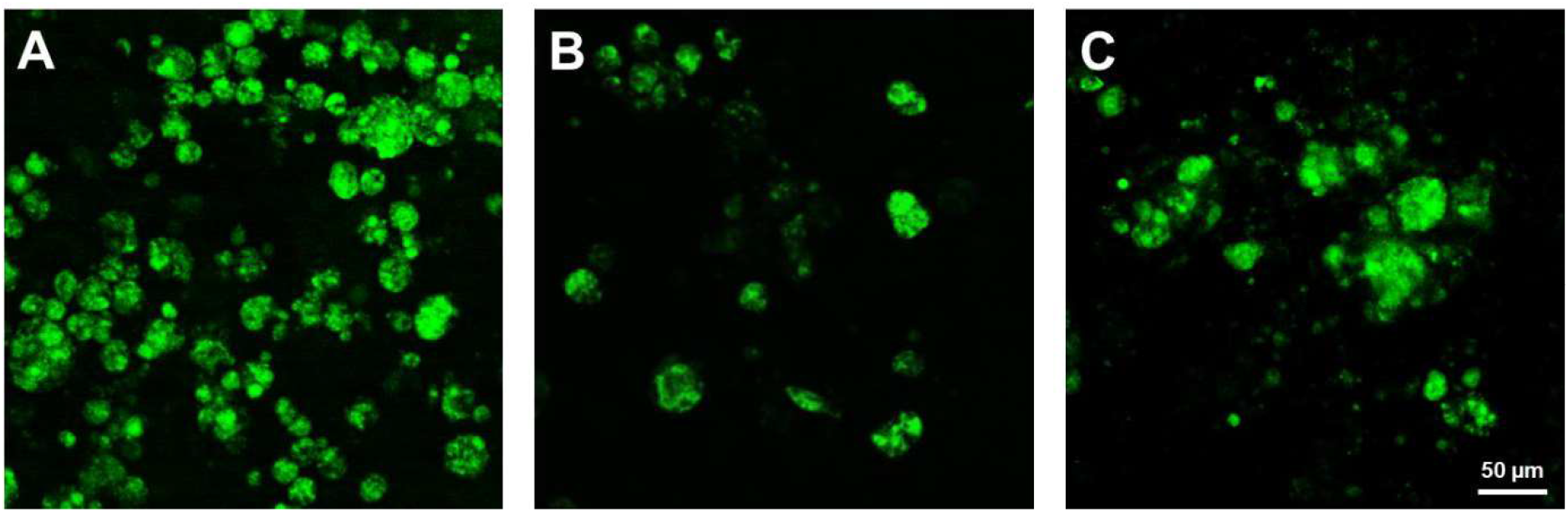
**Syncytium formation.** (A-C) Examples for syncytium formation with different syncytium sizes, assumably with increasing number of contributing cells.

**Figure S12.**
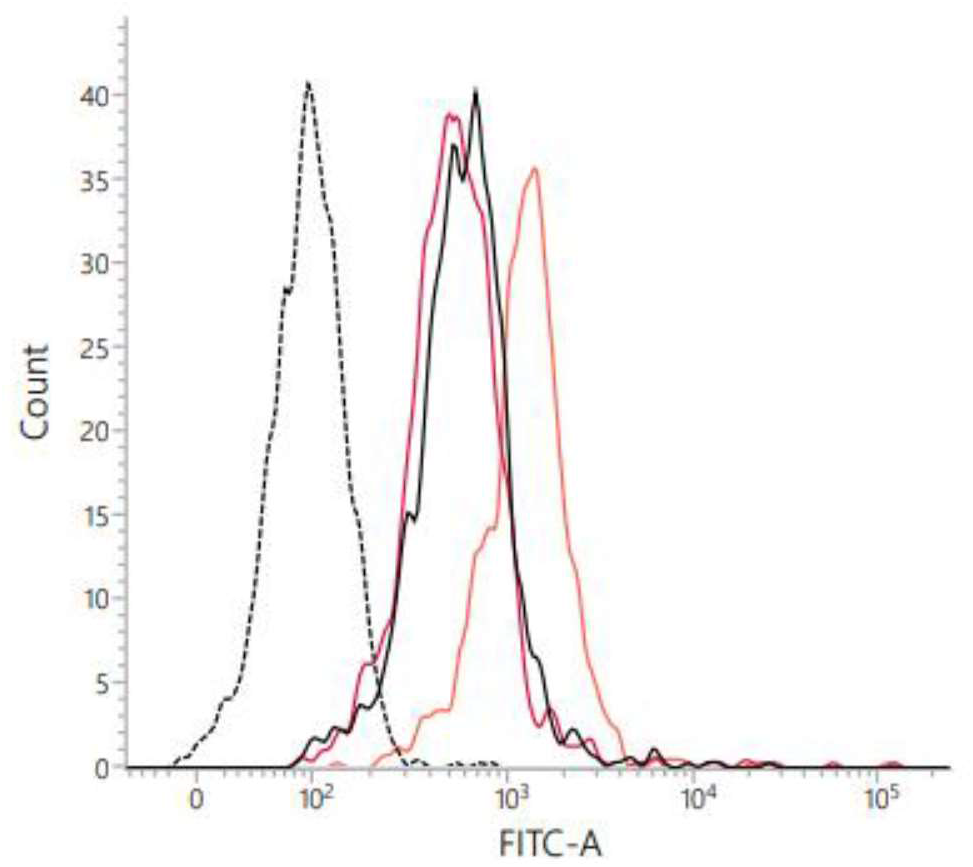
**Results of the flow cytometry measurement.** Dotted black line: untreated cells; Solid black line: BEEF-CP treated cells; solid orange line: ionomycin + BEEF-CP treated cells; solid red line: EGTA + BEEF-CP treated cells.

### Supplementary Tables

**Table S1.** **Tabulated summary of the flow cytometry measurements.**

|  | Fluorescence Intensity Mean (a.u.) |  |  | Ratio (-) |  |  |
| --- | --- | --- | --- | --- | --- | --- |
|  | Untreated | +Ionomycin | +EGTA | Ionomycin/<br>Untreated | EGTA/<br>Untreated | (Ion./Unt.)/<br>(EGTA/Unt.) |
| HEK-293 control | 108 | 189 | 104 | 1.75 | 0.96 | 1.82 |
| HEK-293 + BEEF-CP | 784 | 1601 | 911 | 2.04 | 1.16 | 1.76 |

**Table S2.** **Image parameter values from the manual cell counting analysis.** Each row represents a different image.

| Image ID | Titer<br>TCID<sub>50</sub><br>(mL<sup>-1</sup>) | Variant | Dye<br>conc.<br>(μM) | Health | Initial<br>infection | Infected | Infected<br>vacuoles | Sum | Dead |
| --- | --- | --- | --- | --- | --- | --- | --- | --- | --- |
| 001 | 10<sup>-1</sup> | D614G | 100 | 0 | 0 | 73 | 20 | 93 | 73 |
| 003 | 10<sup>-1</sup> | D614G | 100 | 0 | 0 | 83 | 15 | 98 | 68 |
| 004 | 10<sup>-1</sup> | D614G | 100 | 0 | 0 | 51 | 15 | 66 | 100 |
| 005 | 10<sup>-1</sup> | D614G | 10 | 0 | 0 | 65 | 35 | 100 | 66 |
| 006 | 10<sup>-1</sup> | D614G | 10 | 1 | 16 | 48 | 32 | 97 | 69 |
| 007 | 10<sup>-1</sup> | B.1.1.7 | 100 | 0 | 0 | 68 | 20 | 88 | 78 |
| 008 | 10<sup>-1</sup> | B.1.1.7 | 100 | 0 | 0 | 25 | 22 | 47 | 119 |
| 009 | 10<sup>-1</sup> | B.1.1.7 | 100 | 0 | 0 | 19 | 30 | 49 | 117 |
| 010 | 10<sup>-1</sup> | B.1.1.7 | 10 | 21 | 34 | 8 | 1 | 64 | 102 |
| 014 | 10<sup>-1</sup> | B.1.1.7 | 10 | 3 | 0 | 46 | 46 | 95 | 71 |
| 013 | 10<sup>-1</sup> | B.1.1.7 | 10 | 20 | 0 | 10 | 30 | 60 | 106 |
| 016 | 10<sup>-2</sup> | D614G | 100 | 0 | 0 | 45 | 40 | 85 | 81 |
| 017 | 10<sup>-2</sup> | D614G | 100 | 0 | 0 | 77 | 21 | 98 | 68 |
| 019 | 10<sup>-2</sup> | D614G | 10 | 3 | 1 | 51 | 32 | 87 | 79 |
| 021 | 10<sup>-2</sup> | D614G | 10 | 4 | 7 | 31 | 34 | 76 | 90 |
| 023 | 10<sup>-2</sup> | D614G | 10 | 6 | 7 | 69 | 20 | 102 | 64 |
| 024 | 10<sup>-2</sup> | B.1.1.7 | 100 | 0 | 5 | 31 | 26 | 62 | 104 |
| 027 | 10<sup>-2</sup> | B.1.1.7 | 100 | 0 | 13 | 3 | 0 | 16 | 150 |
| 030 | 10<sup>-2</sup> | B.1.1.7 | 100 | 0 | 8 | 16 | 15 | 39 | 127 |
| 033 | 10<sup>-2</sup> | B.1.1.7 | 10 | 11 | 44 | 5 | 5 | 65 | 101 |
| 036 | 10<sup>-2</sup> | B.1.1.7 | 10 | 20 | 39 | 11 | 0 | 70 | 96 |
| 039 | 10<sup>-2</sup> | B.1.1.7 | 10 | 23 | 29 | 14 | 14 | 80 | 86 |
| 040 | 10<sup>-3</sup> | D614G | 100 | 0 | 0 | 64 | 5 | 69 | 97 |
| 041 | 10<sup>-3</sup> | D614G | 100 | 0 | 0 | 85 | 29 | 114 | 52 |
| 042 | 10<sup>-3</sup> | D614G | 100 | 1 | 0. | 97 | 15 | 113 | 53 |
| 043 | 10<sup>-3</sup> | D614G | 10 | 0 | 0 | 82 | 24 | 106 | 60 |
| 045 | 10<sup>-3</sup> | D614G | 10 | 21 | 33 | 18 | 3 | 75 | 91 |
| 048 | 10<sup>-3</sup> | D614G | 10 | 21 | 9 | 70 | 17 | 117 | 49 |
| 050 | 10<sup>-3</sup> | B.1.1.7 | 100 | 0 | 7 | 9 | 10 | 26 | 140 |
| 052 | 10<sup>-3</sup> | B.1.1.7 | 100 | 18 | 0 | 11 | 11 | 40 | 126 |
| 054 | 10<sup>-3</sup> | B.1.1.7 | 100 | 0 | 1 | 22 | 1 | 24 | 142 |
| 056 | 10<sup>-3</sup> | B.1.1.7 | 10 | 21 | 14 | 60 | 53 | 148 | 18 |
| 058 | 10<sup>-3</sup> | B.1.1.7 | 10 | 15 | 10 | 87 | 30 | 142 | 24 |
| 060 | 10<sup>-4</sup> | D614G | 100 | 181 | 9 | 0 | 0 | 190 | 0 |
| 062 | 10<sup>-4</sup> | D614G | 100 | 177 | 16 | 0 | 0 | 193 | 0 |
| 064 | 10<sup>-4</sup> | D614G | 100 | 184 | 4 | 1 | 0 | 189 | 0 |
| 065 | 10<sup>-4</sup> | D614G | 10 | 139 | 5 | 1 | 0 | 145 | 21 |
| 068 | 10<sup>-4</sup> | D614G | 10 | 139 | 5 | 1 | 0 | 115 | 21 |
| 070 | 10<sup>-4</sup> | D614G | 10 | 120 | 6 | 0 | 0 | 190 | 40 |
| 072 | 10<sup>-4</sup> | B.1.1.7 | 100 | 175 | 8 | 0 | 0 | 183 | 0 |
| 074 | 10<sup>-4</sup> | B.1.1.7 | 100 | 170 | 0 | 0 | 0 | 170 | 0 |
| 076 | 10<sup>-4</sup> | B.1.1.7 | 100 | 176 | 4 | 0 | 0 | 180 | 0 |
| 078 | 10<sup>-4</sup> | B.1.1.7 | 10 | 164 | 28 | 1 | 1 | 194 | 0 |

**Table S2.** continued
| **Image ID** | **Titer**<br>**TCID<sub>50</sub>**<br>**(mL<sup>-1</sup>)** | **Variant** | **Dye**<br>**conc.**<br>**(μM)** | **Health** | **Initial**<br>**infection** | **Infected** | **Infected**<br>**vacuoles** | **Sum** | **Dead** |
| --- | --- | --- | --- | --- | --- | --- | --- | --- | --- |
| 080 | 10<sup>-4</sup> | B.1.1.7 | 10 | 138 | 26 | 20 | 7 | 191 | 0 |
| 083 | 10<sup>-4</sup> | B.1.1.7 | 10 | 10 | 14 | 86 | 17 | 103 | 39 |
| 088 | 10<sup>-5</sup> | D614G | 100 | 180 | 2 | 0 | 0 | 182 | 0 |
| 090 | 10<sup>-5</sup> | D614G | 100 | 171 | 5 | 0 | 0 | 176 | 0 |
| 094 | 10<sup>-5</sup> | D614G | 100 | 161 | 5 | 1 | 0 | 167 | 0 |
| 097 | 10<sup>-5</sup> | D614G | 10 | 160 | 4 | 0 | 0 | 164 | 2 |
| 100 | 10<sup>-5</sup> | D614G | 10 | 177 | 2 | 1 | 0 | 180 | 0 |
| 104 | 10<sup>-5</sup> | B.1.1.7 | 100 | 20 | 0 | 43 | 20 | 83 | 83 |
| 105 | 10<sup>-5</sup> | B.1.1.7 | 100 | 163 | 6 | 0 | 0 | 169 | 0 |
| 107 | 10<sup>-5</sup> | B.1.1.7 | 100 | 152 | 4 | 5 | 0 | 161 | 5 |
| 109 | 10<sup>-5</sup> | B.1.1.7 | 10 | 162 | 8 | 0 | 0 | 170 | 0 |
| 112 | 10<sup>-5</sup> | B.1.1.7 | 10 | 163 | 14 | 0 | 0 | 177 | 0 |
| 113 | 10<sup>-6</sup> | D614G | 100 | 166 | 1 | 0 | 0 | 167 | 0 |
| 116 | 10<sup>-6</sup> | D614G | 100 | 161 | 18 | 1 | 0 | 180 | 0 |
| 118 | 10<sup>-6</sup> | D614G | 10 | 180 | 9 | 2 | 0 | 191 | 0 |
| 119 | 10<sup>-6</sup> | B.1.1.7 | 100 | 161 | 5 | 1 | 0 | 167 | 0 |
| 121 | 10<sup>-6</sup> | B.1.1.7 | 100 | 164 | 8 | 1 | 0 | 173 | 0 |
| 118 | 10<sup>-6</sup> | B.1.1.7 | 10 | 153 | 11 | 1 | 0 | 165 | 1 |
| 126 | 0 | N/A | 100 | 168 | 0 | 0 | 0 | 168 | 0 |
| 125 | 0 | N/A | 100 | 173 | 0 | 0 | 0 | 173 | 0 |
| 131 | 0 | N/A | 100 | 162 | 1 | 0 | 0 | 163 | 0 |
| 136 | 0 | N/A | 10 | 163 | 3 | 0 | 0 | 166 | 0 |
| 145 | 0 | N/A | 10 | 154 | 0 | 0 | 0 | 154 | 0 |
| 143 | 0 | N/A | 10 | 162 | 0 | 0 | 0 | 162 | 0 |

**Table S3.** **The summary of the manual cell counting analysis.**

| Titer<br>TCID<sub>50</sub><br>(mL<sup>-1</sup>) | Variant | Dye<br>conc.<br>(μM) | Mean value |  |  |  |  | Standard deviation |  |  |  |  |
| --- | --- | --- | --- | --- | --- | --- | --- | --- | --- | --- | --- | --- |
|  |  |  | Healthy | Initial<br>infection | Infected | Infected<br>vacuoles, | Dead | Healthy | Initial<br>infection | Infected | Infected<br>vacuoles, | Dead |
| 10<sup>-1</sup> | D614G | 100 | 0.0 | 0.0 | 69.0 | 16.7 | 80.0 | 0.0 | 0.0 | 16.4 | 2.9 | 17.2 |
|  |  | 10 | 0.5 | 8.0 | 56.5 | 33.5 | 67.2 | 0.7 | 11.3 | 12.0 | 2.1 | 2.1 |
|  | B.1.1.7 | 100 | 0.0 | 0.0 | 37.3 | 24.0 | 104.3 | 0.0 | 0.0 | 26.7 | 5.3 | 23.1 |
|  |  | 10 | 14.7 | 11.3 | 21.3 | 25.7 | 92.7 | 10.1 | 19.6 | 21.4 | 22.8 | 19.2 |
| 10<sup>-2</sup> | D614G | 100 | 0.0 | 0.0 | 61.0 | 30.5 | 104.7 | 0.0 | 0.0 | 22.6 | 13.4 | 53.2 |
|  |  | 10 | 4.3 | 5.0 | 50.3 | 28.7 | 77.3 | 1.5 | 3.5 | 19.0 | 7.6 | 13.1 |
|  | B.1.1.7 | 100 | 0.0 | 8.7 | 16.7 | 13.7 | 126.7 | 0.0 | 4.0 | 14.0 | 13.1 | 23.0 |
|  |  | 10 | 18.0 | 37.3 | 10.0 | 6.3 | 94.0 | 6.2 | 7.6 | 4.6 | 7.1 | 7.6 |
| 10<sup>-3</sup> | D614G | 100 | 0.3 | 0.0 | 82.0 | 16.3 | 74.2 | 0.6 | 0.0 | 16.7 | 12.1 | 31.8 |
|  |  | 10 | 14.0 | 14.0 | 56.7 | 14.7 | 66.3 | 12.1 | 17.1 | 34.0 | 10.7 | 21.8 |
|  | B.1.1.7 | 100 | 6.0 | 2.7 | 14.0 | 7.3 | 135.7 | 10.4 | 3.8 | 7.0 | 5.5 | 8.7 |
|  |  | 10 | 18.0 | 12.0 | 73.5 | 41.5 | 20.7 | 4.2 | 2.8 | 19.1 | 16.3 | 4.2 |
| 10<sup>-4</sup> | D614G | 100 | 180.7 | 9.7 | 0.3 | 0.0 | 0.0 | 3.5 | 6.0 | 0.6 | 0.0 | 0.0 |
|  |  | 10 | 132.7 | 5.3 | 0.7 | 0.0 | 27.0 | 11.0 | 0.6 | 0.6 | 0.0 | 11.0 |
|  | B.1.1.7 | 100 | 173.7 | 4.0 | 0.0 | 0.0 | 0.0 | 3.2 | 4.0 | 0.0 | 0.0 | 0.0 |
|  |  | 10 | 104.0 | 22.7 | 35.7 | 8.3 | 0.0 | 82.4 | 7.6 | 44.6 | 8.1 | 0.0 |
| 10<sup>-5</sup> | D614G | 100 | 170.7 | 4.0 | 0.3 | 0.0 | 0.0 | 9.5 | 1.7 | 0.6 | 0.0 | 0.0 |
|  |  | 10 | 168.5 | 3.0 | 0.5 | 0.0 | 0.8 | 12.0 | 1.4 | 0.7 | 0.0 | 1.2 |
|  | B.1.1.7 | 100 | 111.7 | 3.3 | 16.0 | 6.7 | 29.1 | 79.6 | 3.1 | 23.5 | 11.5 | 46.4 |
|  |  | 10 | 162.5 | 11.0 | 0.0 | 0.0 | 0.0 | 0.7 | 4.2 | 0.0 | 0.0 | 0.0 |
| 10<sup>-6</sup> | D614G | 100 | 163.5 | 9.5 | 0.5 | 0.0 | 0.0 | 3.5 | 12.0 | 0.7 | 0.0 | 0.0 |
|  |  | 10 | 180.0 | 9.0 | 2.0 | 0.0 | 0.0 | -<sup>a</sup> | -<sup>a</sup> | -<sup>a</sup> | -<sup>a</sup> | -<sup>a</sup> |
|  | B.1.1.7 | 100 | 162.5 | 6.5 | 1.0 | 0.0 | 0.0 | 2.1 | 2.1 | 0.0 | 0.0 | 0.0 |
|  |  | 10 | 153.0 | 11.0 | 1.0 | 0.0 | 0.7 | -<sup>a</sup> | -<sup>a</sup> | -<sup>a</sup> | -<sup>a</sup> | -<sup>a</sup> |
| 0 | N/A | 100 | 167.7 | 1.0 | 0.0 | 0.0 | 0.0 | 5.5 | 0.6 | 0.0 | 0.0 | 0.0 |
|  |  | 10 | 161.7 | 1.7 | 0.0 | 0.0 | 0.0 | 1.5 | 1.5 | 0.0 | 0.0 | 0.0 |

<sup>a</sup>Only one image was acquired with these parameters.

**Table S4.** **Image parameter values from the automatic image analysis.** Each row represents a different image.

| Image ID | Titer TCID<sub>50</sub> (mL<sup>-1</sup>) | Variant | PMT relative voltage (%) | Dye dilution (μM) | Relative signal area (%) | Image mean intensity (a.u.) | Mean of the threshold area (a.u.) | Maximum particle intensity (a.u.) | Average particle size (px) | Particle percentage area (%) | Particle mean intensity (a.u.) |
| --- | --- | --- | --- | --- | --- | --- | --- | --- | --- | --- | --- |
| 001 | 10<sup>-1</sup> | D614G | 10 | 100 | 27.076 | 512 | 1387 | 5735 | 51.5 | 10.536 | 116.9 |
| 003 | 10<sup>-1</sup> | D614G | 10 | 100 | 30.852 | 574 | 1390 | 6737 | 57.5 | 12.159 | 118.2 |
| 008 | 10<sup>-1</sup> | B.1.1.7 | 10 | 100 | 14.156 | 326 | 1074 | 5724 | 29.4 | 5.338 | 88.3 |
| 009 | 10<sup>-1</sup> | B.1.1.7 | 10 | 100 | 10.875 | 295 | 1010 | 5365 | 20.8 | 4.371 | 83.8 |
| 002 | 10<sup>-1</sup> | D614G | 20 | 100 | 26.412 | 513 | 1431 | 5671 | 110.3 | 10.225 | 158.4 |
| 016 | 10<sup>-2</sup> | D614G | 10 | 100 | 35.909 | 602 | 1333 | 5735 | 55.0 | 13.974 | 115.1 |
| 017 | 10<sup>-2</sup> | D614G | 10 | 100 | 32.637 | 488 | 1150 | 5789 | 69.1 | 12.584 | 120.4 |
| 024 | 10<sup>-2</sup> | B.1.1.7 | 10 | 100 | 19.153 | 306 | 783 | 4385 | 24.1 | 6.675 | 88.0 |
| 025 | 10<sup>-2</sup> | B.1.1.7 | 10 | 100 | 5.076 | 213 | 566 | 2272 | 7.7 | 1.859 | 74.6 |
| 028 | 10<sup>-2</sup> | B.1.1.7 | 10 | 100 | 8.040 | 234 | 740 | 4308 | 13.6 | 3.060 | 79.6 |
| 026 | 10<sup>-2</sup> | B.1.1.7 | 20 | 100 | 4.820 | 217 | 596 | 2430 | 22.3 | 1.251 | 76.6 |
| 029 | 10<sup>-2</sup> | B.1.1.7 | 20 | 100 | 8.002 | 246 | 775 | 4077 | 41.5 | 2.542 | 93.3 |
| 027 | 10<sup>-2</sup> | B.1.1.7 | 30 | 100 | 4.419 | 217 | 618 | 2336 | 28.7 | 0.961 | 76.4 |
| 030 | 10<sup>-2</sup> | B.1.1.7 | 30 | 100 | 7.508 | 245 | 787 | 3986 | 62.1 | 2.310 | 101.1 |
| 040 | 10<sup>-3</sup> | D614G | 10 | 100 | 34.151 | 524 | 1148 | 5049 | 44.1 | 12.434 | 105.3 |
| 041 | 10<sup>-3</sup> | D614G | 10 | 100 | 41.923 | 581 | 1142 | 5604 | 89.4 | 16.491 | 138.8 |
| 049 | 10<sup>-3</sup> | B.1.1.7 | 10 | 100 | 5.839 | 220 | 745 | 3132 | 13.2 | 2.317 | 79.3 |
| 051 | 10<sup>-3</sup> | B.1.1.7 | 10 | 100 | 7.570 | 238 | 671 | 3688 | 11.8 | 2.515 | 77.5 |
| 053 | 10<sup>-3</sup> | B.1.1.7 | 10 | 100 | 7.767 | 240 | 904 | 3862 | 21.3 | 3.061 | 82.2 |
| 050 | 10<sup>-3</sup> | B.1.1.7 | 20 | 100 | 5.824 | 231 | 794 | 3226 | 41.3 | 1.964 | 91.9 |
| 052 | 10<sup>-3</sup> | B.1.1.7 | 20 | 100 | 6.503 | 233 | 727 | 3394 | 34.0 | 1.876 | 84.4 |
| 054 | 10<sup>-3</sup> | B.1.1.7 | 20 | 100 | 7.761 | 245 | 925 | 3800 | 76.3 | 2.813 | 105.0 |
| 059 | 10<sup>-4</sup> | D614G | 20 | 100 | 0.136 | 200 | 354 | 480 | 5.0 | 0.116 | 69.7 |
| 061 | 10<sup>-4</sup> | D614G | 20 | 100 | 0.068 | 150 | 394 | 645 | 10.3 | 0.088 | 72.3 |
| 063 | 10<sup>-4</sup> | D614G | 20 | 100 | 0.331 | 179 | 532 | 1206 | 9.3 | 0.184 | 71.3 |
| 071 | 10<sup>-4</sup> | B.1.1.7 | 20 | 100 | 0.011 | 198 | 349 | 470 | 3.0 | 0.057 | 67.7 |
| 073 | 10<sup>-4</sup> | B.1.1.7 | 20 | 100 | 0.022 | 189 | 346 | 441 | 3.5 | 0.051 | 67.4 |
| 075 | 10<sup>-4</sup> | B.1.1.7 | 20 | 100 | 0.134 | 196 | 507 | 962 | 5.5 | 0.155 | 69.2 |
| 060 | 10<sup>-4</sup> | D614G | 30 | 100 | 0.105 | 205 | 341 | 431 | 5.4 | 0.073 | 68.8 |
| 062 | 10<sup>-4</sup> | D614G | 30 | 100 | 0.063 | 150 | 372 | 571 | 15.5 | 0.049 | 73.7 |
| 064 | 10<sup>-4</sup> | D614G | 30 | 100 | 0.325 | 185 | 508 | 1102 | 12.3 | 0.161 | 71.7 |
| 072 | 10<sup>-4</sup> | B.1.1.7 | 30 | 100 | 0.008 | 202 | 338 | 401 | 3.8 | 0.049 | 67.5 |
| 074 | 10<sup>-4</sup> | B.1.1.7 | 30 | 100 | 0.011 | 191 | 341 | 417 | 4.8 | 0.037 | 66.9 |
| 076 | 10<sup>-4</sup> | B.1.1.7 | 30 | 100 | 0.131 | 202 | 462 | 812 | 8.2 | 0.133 | 69.1 |
| 084 | 10<sup>-5</sup> | D614G | 10 | 100 | 0.179 | 171 | 368 | 726 | 3.5 | 0.992 | 71.6 |
| 085 | 10<sup>-5</sup> | D614G | 10 | 100 | 0.090 | 157 | 358 | 551 | 2.7 | 0.465 | 70.2 |
| 090 | 10<sup>-5</sup> | D614G | 10 | 100 | 0.189 | 161 | 494 | 1390 | 3.6 | 0.641 | 71.3 |
| 101 | 10<sup>-5</sup> | B.1.1.7 | 10 | 100 | 11.604 | 243 | 806 | 3111 | 34.6 | 4.227 | 91.7 |
| 102 | 10<sup>-5</sup> | B.1.1.7 | 10 | 100 | 20.042 | 325 | 904 | 3938 | 41.9 | 7.312 | 98.2 |
| 086 | 10<sup>-5</sup> | D614G | 20 | 100 | 0.063 | 172 | 375 | 612 | 4.2 | 0.194 | 68.6 |
| 087 | 10<sup>-5</sup> | D614G | 20 | 100 | 0.117 | 168 | 340 | 447 | 3.9 | 0.067 | 67.7 |
| 091 | 10<sup>-5</sup> | D614G | 20 | 100 | 0.113 | 159 | 546 | 1249 | 6.6 | 0.168 | 69.4 |

Table S4 continued
| Image ID | Titer TCID<sub>50</sub> (mL<sup>-1</sup>) | Variant | PMT relative voltage (%) | Dye dilution (μM) | Relative signal area (%) | Image mean intensity (a.u.) | Mean of the threshold area (a.u.) | Maximum particle intensity (a.u.) | Average particle size (px) | Particle percentage area (%) | Particle mean intensity (a.u.) |
| --- | --- | --- | --- | --- | --- | --- | --- | --- | --- | --- | --- |
| 093 | 10<sup>-5</sup> | D614G | 20 | 100 | 0.075 | 175 | 374 | 619 | 5.7 | 0.047 | 67.4 |
| 103 | 10<sup>-5</sup> | B.1.1.7 | 20 | 100 | 12.014 | 261 | 835 | 3074 | 77.8 | 4.141 | 110.6 |
| 104 | 10<sup>-5</sup> | B.1.1.7 | 20 | 100 | 20.425 | 341 | 949 | 3823 | 87.5 | 7.308 | 116.0 |
| 105 | 10<sup>-5</sup> | B.1.1.7 | 20 | 100 | 0.102 | 190 | 435 | 761 | 8.2 | 0.110 | 69.9 |
| 107 | 10<sup>-5</sup> | B.1.1.7 | 20 | 100 | 0.044 | 186 | 402 | 886 | 7.2 | 0.066 | 71.0 |
| 088 | 10<sup>-5</sup> | D614G | 30 | 100 | 0.048 | 170 | 348 | 485 | 5.8 | 0.222 | 67.5 |
| 089 | 10<sup>-5</sup> | D614G | 30 | 100 | 0.019 | 164 | 330 | 366 | 4.1 | 0.016 | 66.0 |
| 092 | 10<sup>-5</sup> | D614G | 30 | 100 | 0.171 | 160 | 619 | 4018 | 7.5 | 0.151 | 73.4 |
| 094 | 10<sup>-5</sup> | D614G | 30 | 100 | 0.098 | 178 | 359 | 505 | 8.0 | 0.029 | 66.5 |
| 106 | 10<sup>-5</sup> | B.1.1.7 | 30 | 100 | 0.087 | 194 | 398 | 594 | 11.6 | 0.080 | 71.6 |
| 108 | 10<sup>-5</sup> | B.1.1.7 | 30 | 100 | 0.026 | 191 | 383 | 672 | 8.0 | 0.033 | 70.6 |
| 113 | 10<sup>-6</sup> | D614G | 20 | 100 | 0.106 | 207 | 425 | 662 | 4.3 | 0.073 | 68.4 |
| 115 | 10<sup>-6</sup> | D614G | 20 | 100 | 0.060 | 171 | 419 | 645 | 4.7 | 0.142 | 69.4 |
| 114 | 10<sup>-6</sup> | D614G | 30 | 100 | 0.092 | 211 | 398 | 554 | 5.4 | 0.040 | 67.9 |
| 116 | 10<sup>-6</sup> | D614G | 30 | 100 | 0.058 | 171 | 406 | 584 | 9.0 | 0.067 | 69.9 |
| 119 | 10<sup>-6</sup> | B.1.1.7 | 30 | 100 | 0.054 | 161 | 455 | 764 | 17.4 | 0.032 | 72.0 |
| 120 | 10<sup>-6</sup> | B.1.1.7 | 40 | 100 | 0.037 | 161 | 442 | 645 | 40.3 | 0.024 | 79.1 |
| 123 | Reference | N/A | 10 | 100 | 0.133 | 165 | 496 | 1063 | 3.2 | 0.646 | 71.5 |
| 124 | Reference | N/A | 10 | 100 | 0.190 | 155 | 453 | 3415 | 3.4 | 0.630 | 71.6 |
| 129 | Reference | N/A | 10 | 100 | 0.359 | 187 | 435 | 1150 | 3.1 | 0.491 | 71.1 |
| 125 | Reference | N/A | 20 | 100 | 0.093 | 168 | 536 | 906 | 5.5 | 0.167 | 69.6 |
| 126 | Reference | N/A | 20 | 100 | 0.124 | 163 | 411 | 962 | 5.9 | 0.157 | 69.4 |
| 130 | Reference | N/A | 20 | 100 | 0.375 | 189 | 444 | 1117 | 11.2 | 0.160 | 75.5 |
| 131 | Reference | N/A | 20 | 100 | 0.051 | 193 | 408 | 563 | 10.6 | 0.060 | 72.4 |
| 144 | Reference | N/A | 20 | 100 | 0.047 | 200 | 404 | 1044 | 3.6 | 0.121 | 68.2 |
| 146 | Reference | N/A | 20 | 100 | 0.062 | 189 | 702 | 1747 | 4.4 | 0.081 | 69.7 |
| 147 | Reference | N/A | 20 | 100 | 0.016 | 192 | 330 | 361 | 3.2 | 0.093 | 68.0 |
| 150 | Reference | N/A | 20 | 100 | 0.060 | 199 | 457 | 1084 | 5.1 | 0.048 | 70.3 |
| 127 | Reference | N/A | 30 | 100 | 0.075 | 165 | 494 | 708 | 7.9 | 0.124 | 68.4 |
| 128 | Reference | N/A | 30 | 100 | 0.094 | 162 | 425 | 2112 | 7.0 | 0.093 | 72.1 |
| 132 | Reference | N/A | 30 | 100 | 0.381 | 198 | 426 | 962 | 17.6 | 0.101 | 77.2 |
| 133 | Reference | N/A | 30 | 100 | 0.043 | 198 | 392 | 526 | 12.4 | 0.037 | 70.4 |
| 145 | Reference | N/A | 30 | 100 | 0.051 | 204 | 393 | 934 | 5.0 | 0.077 | 68.2 |
| 148 | Reference | N/A | 30 | 100 | 0.051 | 193 | 704 | 1566 | 5.4 | 0.057 | 68.7 |
| 149 | Reference | N/A | 30 | 100 | 0.006 | 196 | 324 | 335 | 3.7 | 0.034 | 66.9 |
| 151 | Reference | N/A | 30 | 100 | 0.079 | 204 | 401 | 901 | 7.4 | 0.030 | 70.5 |
| 004 | 10<sup>-1</sup> | D614G | 10 | 10 | 15.320 | 223 | 476 | 1346 | 13.2 | 4.112 | 80.3 |
| 005 | 10<sup>-1</sup> | D614G | 10 | 10 | 17.002 | 219 | 459 | 1183 | 17.7 | 5.264 | 81.5 |
| 006 | 10<sup>-1</sup> | D614G | 10 | 10 | 25.483 | 259 | 443 | 1183 | 11.7 | 6.250 | 79.3 |
| 007 | 10<sup>-1</sup> | B.1.1.7 | 10 | 10 | 26.296 | 456 | 1144 | 6094 | 42.7 | 9.644 | 101.9 |
| 010 | 10<sup>-1</sup> | B.1.1.7 | 10 | 10 | 1.542 | 158 | 405 | 933 | 9.4 | 0.974 | 75.2 |
| 011 | 10<sup>-1</sup> | B.1.1.7 | 10 | 10 | 12.181 | 254 | 548 | 2577 | 10.9 | 3.461 | 77.7 |
| 014 | 10<sup>-1</sup> | B.1.1.7 | 10 | 10 | 13.541 | 211 | 450 | 1488 | 10.8 | 3.594 | 78.5 |
| 012 | 10<sup>-1</sup> | B.1.1.7 | 20 | 10 | 1.189 | 150 | 409 | 843 | 16.8 | 0.551 | 75.5 |

Table S4 continued
| Image ID | Titer TCID<sub>50</sub> (mL<sup>-1</sup>) | Variant | PMT relative voltage (%) | Dye dilution (μM) | Relative signal area (%) | Image mean intensity (a.u.) | Mean of the threshold area (a.u.) | Maximum particle intensity (a.u.) | Average particle size (px) | Particle percentage area (%) | Particle mean intensity (a.u.) |
| --- | --- | --- | --- | --- | --- | --- | --- | --- | --- | --- | --- |
| 013 | 10<sup>-1</sup> | B.1.1.7 | 20 | 10 | 0.000 | 113 | 323 | 325 | 3.4 | 0.004 | 66.1 |
| 015 | 10<sup>-1</sup> | B.1.1.7 | 20 | 10 | 11.896 | 202 | 455 | 1361 | 17.4 | 2.559 | 75.9 |
| 018 | 10<sup>-2</sup> | D614G | 10 | 10 | 11.899 | 194 | 431 | 1118 | 11.7 | 3.766 | 78.1 |
| 020 | 10<sup>-2</sup> | D614G | 10 | 10 | 7.604 | 182 | 427 | 965 | 10.0 | 2.381 | 77.4 |
| 022 | 10<sup>-2</sup> | D614G | 10 | 10 | 14.106 | 194 | 453 | 1858 | 16.3 | 4.603 | 81.4 |
| 031 | 10<sup>-2</sup> | B.1.1.7 | 10 | 10 | 0.945 | 143 | 375 | 704 | 6.9 | 0.811 | 74.0 |
| 034 | 10<sup>-2</sup> | B.1.1.7 | 10 | 10 | 3.280 | 171 | 387 | 737 | 6.9 | 1.502 | 74.4 |
| 037 | 10<sup>-2</sup> | B.1.1.7 | 10 | 10 | 8.701 | 195 | 428 | 1325 | 9.0 | 2.531 | 76.6 |
| 019 | 10<sup>-2</sup> | D614G | 20 | 10 | 11.167 | 190 | 435 | 1094 | 18.7 | 2.838 | 76.2 |
| 021 | 10<sup>-2</sup> | D614G | 20 | 10 | 6.804 | 174 | 428 | 909 | 15.7 | 1.612 | 74.5 |
| 023 | 10<sup>-2</sup> | D614G | 20 | 10 | 13.932 | 194 | 452 | 1193 | 23.6 | 3.776 | 78.0 |
| 032 | 10<sup>-2</sup> | B.1.1.7 | 20 | 10 | 0.770 | 147 | 371 | 678 | 10.9 | 0.434 | 71.2 |
| 035 | 10<sup>-2</sup> | B.1.1.7 | 20 | 10 | 2.889 | 167 | 389 | 724 | 11.4 | 0.858 | 72.3 |
| 038 | 10<sup>-2</sup> | B.1.1.7 | 20 | 10 | 7.344 | 187 | 436 | 1391 | 14.4 | 1.563 | 74.2 |
| 033 | 10<sup>-2</sup> | B.1.1.7 | 30 | 10 | 0.503 | 145 | 361 | 571 | 11.5 | 0.204 | 69.7 |
| 036 | 10<sup>-2</sup> | B.1.1.7 | 30 | 10 | 2.480 | 166 | 385 | 700 | 11.7 | 0.470 | 70.1 |
| 039 | 10<sup>-2</sup> | B.1.1.7 | 30 | 10 | 6.232 | 183 | 428 | 1127 | 15.7 | 0.967 | 72.4 |
| 042 | 10<sup>-3</sup> | D614G | 10 | 10 | 18.344 | 219 | 446 | 1052 | 13.9 | 5.063 | 79.8 |
| 043 | 10<sup>-3</sup> | D614G | 10 | 10 | 17.480 | 213 | 449 | 1281 | 15.0 | 5.088 | 80.8 |
| 045 | 10<sup>-3</sup> | D614G | 10 | 10 | 4.184 | 169 | 395 | 1041 | 6.5 | 1.162 | 75.1 |
| 047 | 10<sup>-3</sup> | D614G | 10 | 10 | 16.401 | 205 | 465 | 1205 | 17.9 | 4.987 | 81.9 |
| 055 | 10<sup>-3</sup> | B.1.1.7 | 10 | 10 | 14.771 | 209 | 421 | 1902 | 10.8 | 4.630 | 77.9 |
| 057 | 10<sup>-3</sup> | B.1.1.7 | 10 | 10 | 17.851 | 215 | 442 | 1172 | 13.8 | 5.363 | 79.9 |
| 044 | 10<sup>-3</sup> | D614G | 20 | 10 | 16.067 | 204 | 438 | 1011 | 19.5 | 3.751 | 76.3 |
| 046 | 10<sup>-3</sup> | D614G | 20 | 10 | 3.670 | 159 | 395 | 24531 | 10.4 | 0.523 | 84.9 |
| 048 | 10<sup>-3</sup> | D614G | 20 | 10 | 16.181 | 200 | 462 | 1081 | 26.7 | 4.288 | 79.7 |
| 056 | 10<sup>-3</sup> | B.1.1.7 | 20 | 10 | 14.315 | 207 | 423 | 1262 | 17.4 | 3.423 | 75.0 |
| 058 | 10<sup>-3</sup> | B.1.1.7 | 20 | 10 | 16.959 | 210 | 440 | 1074 | 20.5 | 4.133 | 77.5 |
| 081 | 10<sup>-4</sup> | B.1.1.7 | 10 | 10 | 17.838 | 214 | 498 | 2827 | 18.1 | 5.222 | 82.9 |
| 065 | 10<sup>-4</sup> | D614G | 20 | 10 | 0.205 | 152 | 436 | 853 | 7.2 | 0.177 | 70.1 |
| 067 | 10<sup>-4</sup> | D614G | 20 | 10 | 0.102 | 152 | 539 | 1114 | 13.3 | 0.086 | 71.7 |
| 069 | 10<sup>-4</sup> | D614G | 20 | 10 | 3.798 | 145 | 416 | 893 | 17.6 | 0.981 | 75.1 |
| 077 | 10<sup>-4</sup> | B.1.1.7 | 20 | 10 | 0.257 | 142 | 376 | 569 | 9.5 | 0.184 | 70.1 |
| 079 | 10<sup>-4</sup> | B.1.1.7 | 20 | 10 | 1.426 | 138 | 380 | 662 | 8.1 | 0.280 | 71.0 |
| 082 | 10<sup>-4</sup> | B.1.1.7 | 20 | 10 | 17.814 | 207 | 468 | 1077 | 24.5 | 4.321 | 78.5 |
| 066 | 10<sup>-4</sup> | D614G | 30 | 10 | 0.219 | 152 | 420 | 777 | 12.2 | 0.109 | 74.5 |
| 068 | 10<sup>-4</sup> | D614G | 30 | 10 | 0.086 | 152 | 507 | 947 | 20.3 | 0.065 | 71.6 |
| 070 | 10<sup>-4</sup> | D614G | 30 | 10 | 3.503 | 146 | 403 | 772 | 18.5 | 0.689 | 72.7 |
| 078 | 10<sup>-4</sup> | B.1.1.7 | 30 | 10 | 0.159 | 142 | 379 | 528 | 11.4 | 0.077 | 69.5 |
| 080 | 10<sup>-4</sup> | B.1.1.7 | 30 | 10 | 1.440 | 139 | 375 | 645 | 8.4 | 0.160 | 68.6 |
| 083 | 10<sup>-4</sup> | B.1.1.7 | 30 | 10 | 16.653 | 201 | 448 | 1002 | 25.0 | 3.269 | 75.6 |
| 095 | 10<sup>-5</sup> | D614G | 20 | 10 | 0.042 | 150 | 516 | 919 | 7.6 | 0.114 | 71.3 |
| 097 | 10<sup>-5</sup> | D614G | 20 | 10 | 0.093 | 153 | 489 | 856 | 9.6 | 0.086 | 73.0 |
| 098 | 10<sup>-5</sup> | D614G | 20 | 10 | 0.048 | 153 | 445 | 695 | 5.6 | 0.114 | 68.9 |

**Table S4 continued**
| Image ID | Titer TCID<sub>50</sub> (mL<sup>-1</sup>) | Variant | PMT relative voltage (%) | Dye dilution (μM) | Relative signal area (%) | Image mean intensity (a.u.) | Mean of the threshold area (a.u.) | Maximum particle intensity (a.u.) | Average particle size (px) | Particle percentage area (%) | Particle mean intensity (a.u.) |
| --- | --- | --- | --- | --- | --- | --- | --- | --- | --- | --- | --- |
| 109 | 10<sup>-5</sup> | B.1.1.7 | 20 | 10 | 0.025 | 150 | 407 | 546 | 7.6 | 0.056 | 71.1 |
| 111 | 10<sup>-5</sup> | B.1.1.7 | 20 | 10 | 0.065 | 156 | 444 | 764 | 7.7 | 0.100 | 70.7 |
| 096 | 10<sup>-5</sup> | D614G | 30 | 10 | 0.038 | 153 | 468 | 800 | 8.8 | 0.094 | 69.8 |
| 099 | 10<sup>-5</sup> | D614G | 30 | 10 | 0.075 | 153 | 453 | 746 | 11.2 | 0.073 | 70.4 |
| 100 | 10<sup>-5</sup> | D614G | 30 | 10 | 0.039 | 153 | 423 | 615 | 8.8 | 0.048 | 70.2 |
| 110 | 10<sup>-5</sup> | B.1.1.7 | 30 | 10 | 0.019 | 150 | 408 | 2417 | 7.5 | 0.034 | 77.2 |
| 112 | 10<sup>-5</sup> | B.1.1.7 | 30 | 10 | 0.044 | 154 | 428 | 647 | 9.1 | 0.069 | 69.7 |
| 117 | 10<sup>-6</sup> | D614G | 30 | 10 | 0.045 | 118 | 501 | 742 | 94.3 | 0.029 | 100.0 |
| 121 | 10<sup>-6</sup> | B.1.1.7 | 30 | 10 | 0.004 | 147 | 343 | 373 | 7.3 | 0.018 | 68.1 |
| 118 | 10<sup>-6</sup> | D614G | 40 | 10 | 0.039 | 118 | 467 | 647 | 86.3 | 0.026 | 80.4 |
| 122 | 10<sup>-6</sup> | B.1.1.7 | 40 | 10 | 0.000 | 145 | 330 | 332 | 6.3 | 0.017 | 67.6 |
| 134 | Reference | N/A | 10 | 10 | 0.003 | 149 | 330 | 355 | 2.5 | 0.308 | 70.3 |
| 136 | Reference | N/A | 20 | 10 | 0.078 | 152 | 562 | 1341 | 6.4 | 0.149 | 71.3 |
| 138 | Reference | N/A | 20 | 10 | 0.076 | 142 | 405 | 586 | 14.6 | 0.069 | 74.8 |
| 140 | Reference | N/A | 20 | 10 | 0.017 | 150 | 478 | 695 | 4.2 | 0.106 | 68.3 |
| 141 | Reference | N/A | 20 | 10 | 0.016 | 144 | 355 | 424 | 6.1 | 0.050 | 69.8 |
| 137 | Reference | N/A | 30 | 10 | 0.001 | 151 | 327 | 328 | 3.7 | 0.017 | 66.6 |
| 139 | Reference | N/A | 30 | 10 | 0.064 | 142 | 376 | 480 | 21.4 | 0.043 | 78.9 |
| 142 | Reference | N/A | 30 | 10 | 0.014 | 151 | 455 | 624 | 6.5 | 0.050 | 68.3 |
| 143 | Reference | N/A | 30 | 10 | 0.007 | 146 | 350 | 386 | 5.7 | 0.024 | 67.0 |

### ImageJ Script

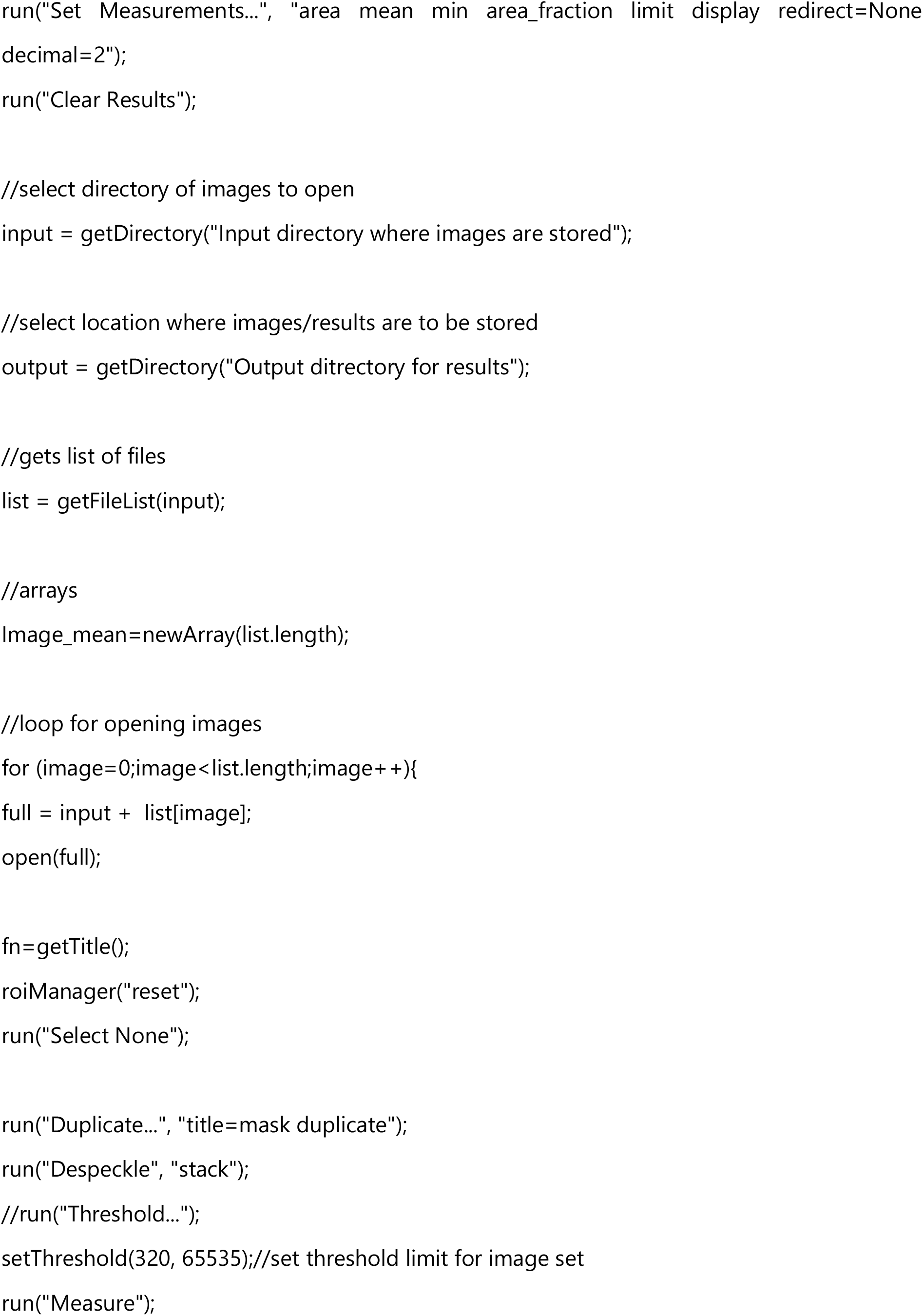
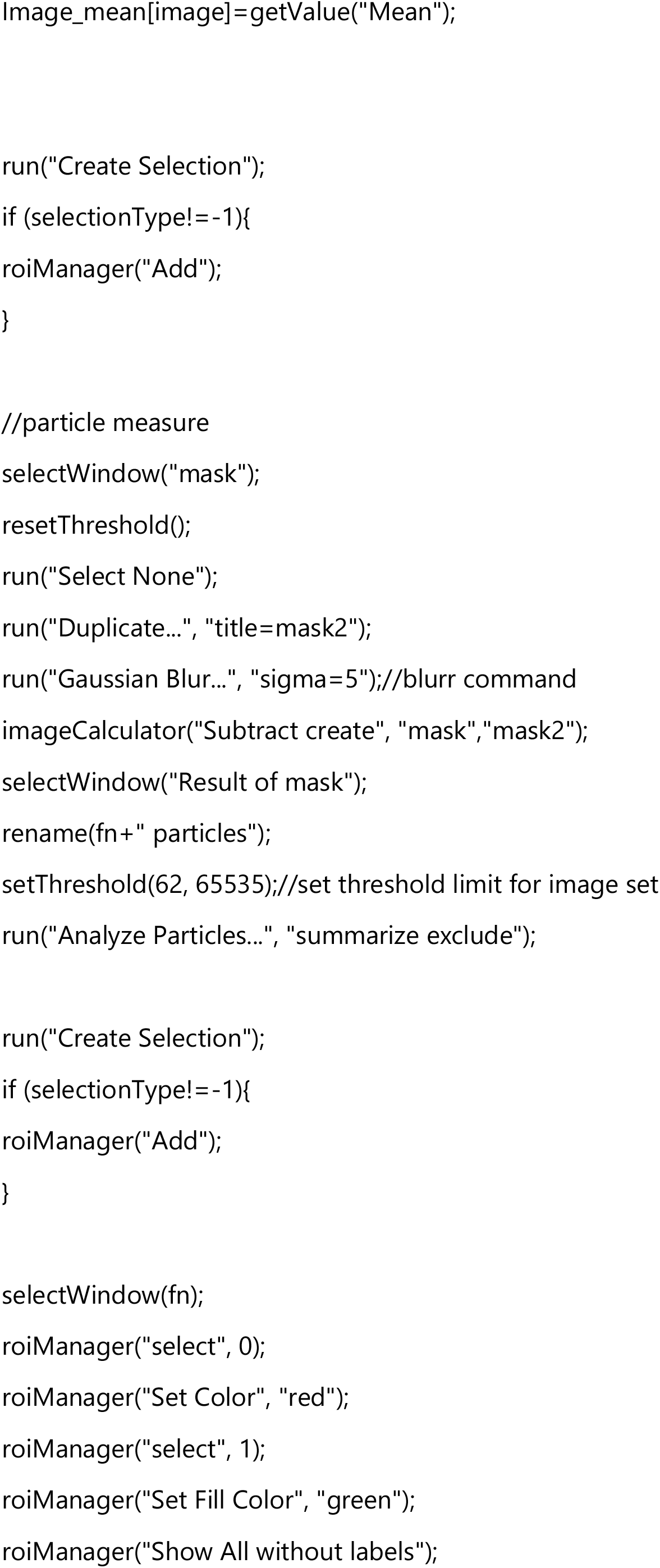
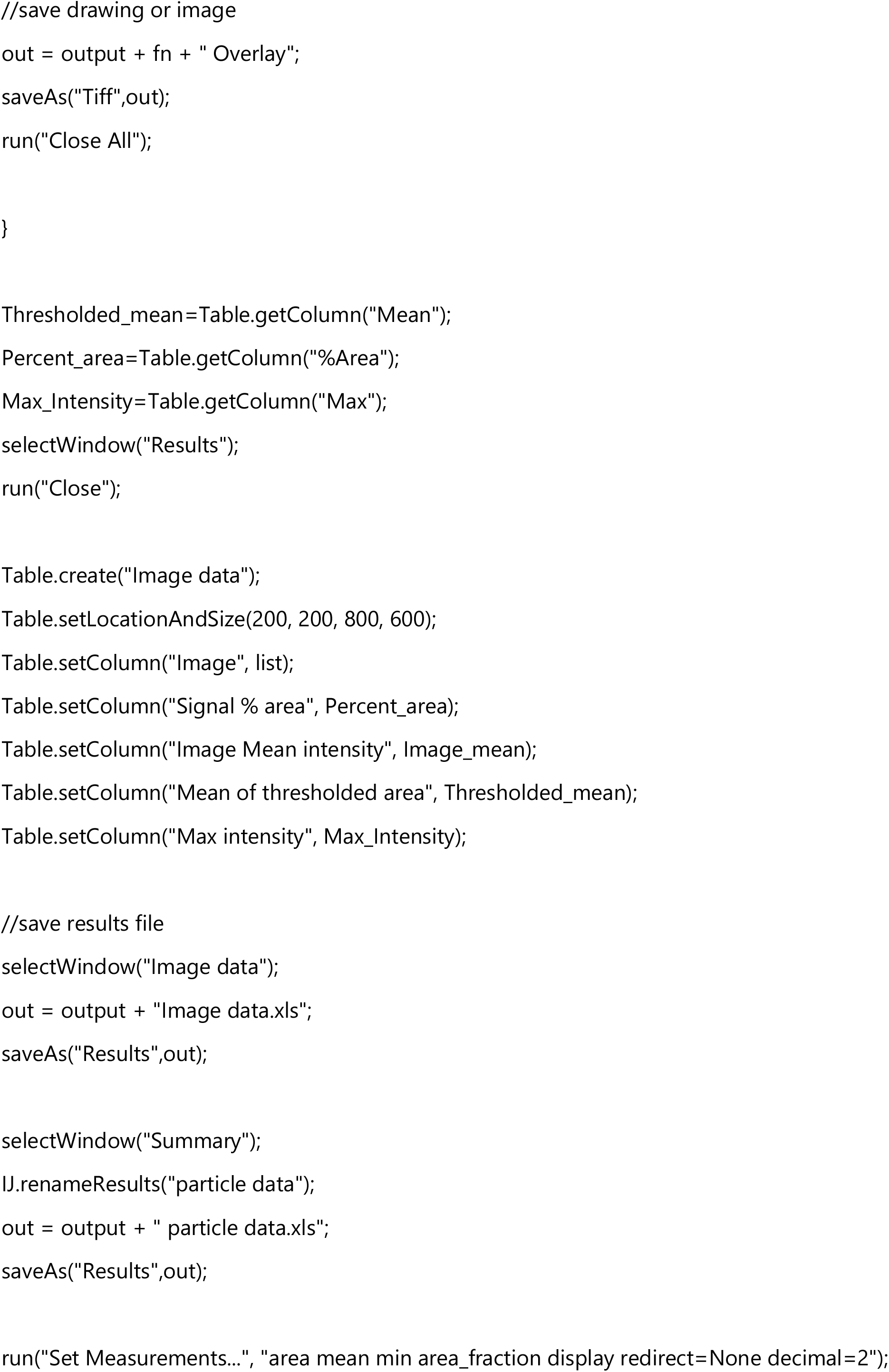

### Synthetic Notes

#### General synthetic methodologies and characterization

All the starting materials, reagents and solvents were purchased from Sigma-Aldrich (Merck) in reagent grade and used as received. NMR solvents were purchased from Eurisotop. Reactions were monitored by LC-MS (Shimazu MS2020, Supelco Ascentis, 2.0 × 50 mm, 2.1 pm C18 column; injection of 1 pi; 5-98% MeCN/H_2_O, linear gradient, with constant 0.1% v/v TFA additive; run of 6 min; flow of 0.8 ml min“^1^; ESI; positive ion mode). Reaction products were purified by gradient elution preparative HPLC (HPLC Gilson 333 instrument, UV detector 220 nm) on a Phenomenex Gemini C18,250×50.00 mm; 10 pm, 110A column using 0.2% v/v TFA in water and acetonitrile as the mobile phase components.

^1^H NMR and ^13^C NMR spectra were recorded at 600/400 MHz, 151/101 MHz on Bruker Avance 600 or 400 spectrometers. All chemical shifts are quoted in parts per million (ppm), measured from the center of the signal except in the case of multiplets, which are quoted as a range. ^1^H NMR and ^13^C chemical shifts are referenced to the residual solvent peak of (CD_3_)_2_SO (^1^H referenced to 2.50 ppm and ^13^C referenced to 39.52 ppm) or CDCl_3_ (^1^H referenced to 7.26 ppm and ^13^C referenced to 77.16 ppm). Coupling constants are given with an accuracy of 0.1 Hz. Splitting patterns are abbreviated as follows: singlet (s), doublet (d), triplet (t), quartet (q), multiplet (m), broad singlet (bs) and combinations thereof. Assignment of spectra was aided by 2D NMR spectroscopy (^1^H-^13^C HSQC and HMBC).

HRMS and MS-MS analyses were performed on a Thermo Velos Pro Orbitrap Elite (Thermo Fisher Scientific) system. The ionization method was ESI operated in positive (or negative) ion mode. The protonated (or deprotonated) molecular ion peaks were fragmented by collision-induced dissociation (CID) at a normalized collision energy of 40-65%. For the CID experiment helium was used as the collision gas. The samples were dissolved in methanol. Data acquisition and analysis were accomplished with Xcalibur software version 4.0 (Thermo Fisher Scientific).

The preparation of the calcium dye was carried out in a five-step synthetic route using 4-ethyl resorcinol, 1,2,4-benzenetricarboxylic anhydride and tetraethyl BAPTA23 in high overall yield.

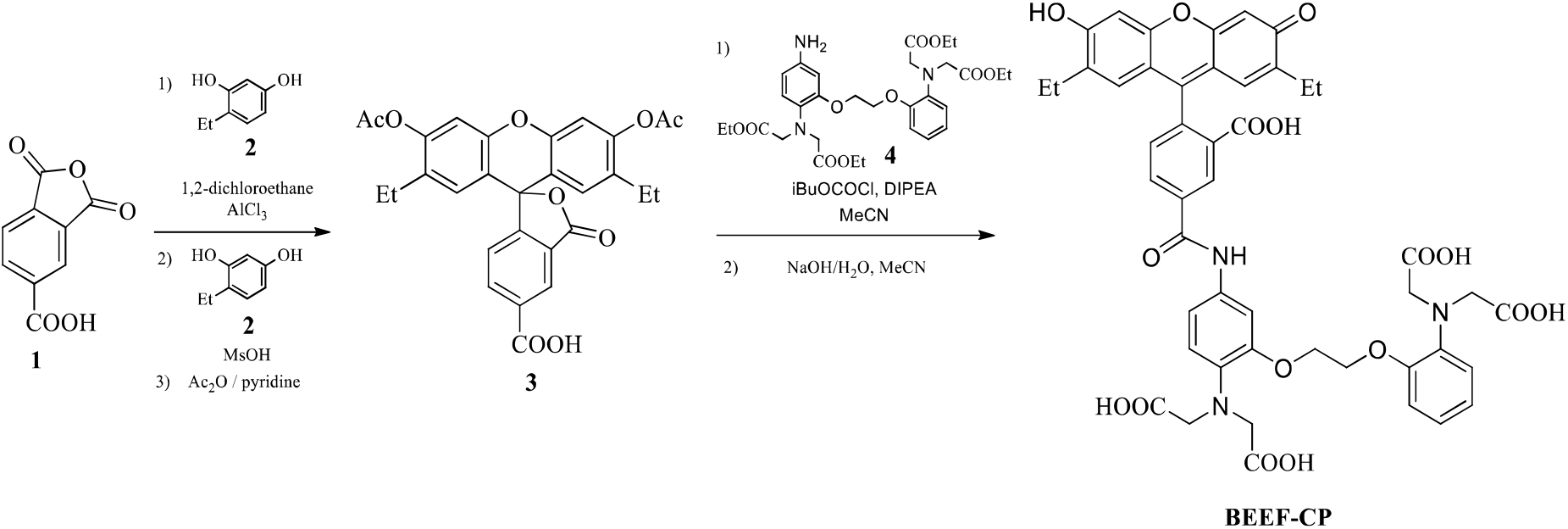

#### Synthesis of ketone 5

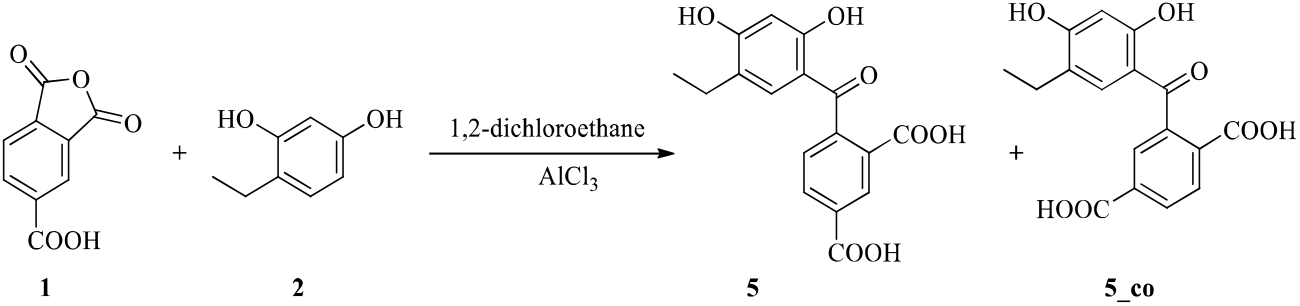

4-Ethylresorcinol (2, 1.38 g, 10.0 mmol, 1.0 equiv.) and 1,2,4-benzenetricarboxylic anhydride (1, 2.11 g, 11.0 mmol, 1.1 equiv.) were dissolved in 1,2-dichloroethane (55.0 mL). AlCl_3_ (7.99 g, 60 mmol, 6.0 equiv.) was added, the mixture was stirred for 72 hours at room temperature. The solvent was evaporated, 100 mL of EtOAc, then 50 mL of aqueous HCl (4M) were added then the layers were separated. The aqueous phase was washed with EtOAc (3 × 100 mL), the combined organic layers was washed with aqueous HCl (1M, 50 mL), brine (50 mL), dried over MgSO_4_. The solvent was evaporated under reduced pressure. The residue (3.51 g) was purified by preparative HPLC (water–acetonitrile–0.1% TFA, using the gradient method). After purification, the fractions were lyophilized. The target isomer (5, 1.49 g, yield 45%) and its regioisomer (**5_co** 1.02 g, yield 31%) were isolated as a yellow powder.

##### Spectroscopic data of 5

_’_H NMR (400 MHz, DMSO-d6) δ 13.26 (s, 2H), 11.86 (s, 1H), 10.62 (s, 1H), 8.50 (d, J = 1.6 Hz, 1H), 8.21 (dd, J = 7.9, 1.7 Hz, 1H), 7.54 (d, J = 7.9 Hz, 1H), 6.80 (s, 1H), 6.39 (s, 1H), 2.34 (q, J = 7.4 Hz, 2H), 0.95 (t, J = 7.5 Hz, 3H). ^13^C NMR (101 MHz, DMSO-d6) δ 198.9,166.0,163.1,162.4,143.8,132.6, 131.8, 130.5, 130.0, 127.9, 122.7, 112.7, 102.2, 21.7, 13.8. HR-MS:(ESI) calcd. for C_17_H_13_O_7_ 329.0667 [M-H]^+^, found 329.06700, Df. = 1.0 ppm. HR-ESI-MS-MS (CID=40%; rel. int. %): 285(2); 191(100) and 147(9).

##### Spectroscopic data of 5_co

^1^H NMR (400 MHz, DMSO-d6) δ 8.15 (dd, J = 8.1, 1.7 Hz, III), 8.06 (d, J = 8.1 Hz, 1H), 7.85 (d, J = 1.6 Hz, 1H), 6.83 (s, 1H), 6.39 (s, 1H), 2.35 (q, J = 7.4 Hz, 2H), 0.96 (t, J = 7.5 Hz, 3H). ^13^C NMR (101 MHz, DMSO-d6) δ 198.7, 166.4, 166.0, 163.0, 162.5, 140.2, 133.8, 133.8, 132.4, 130.2, 130.1, 127.9, 122.6, 112.8, 102.2, 21.7, 13.8.

#### Synthesis of unprotected fluorescein derivative 6

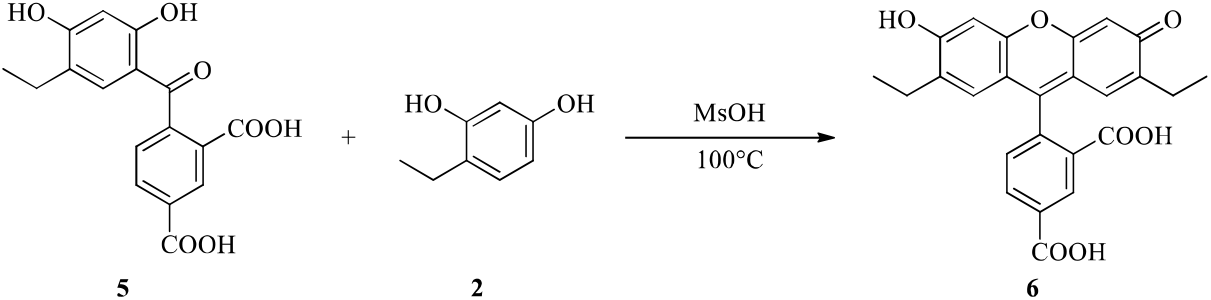

**5** (0.90 g, 2.73 mmol, 1.0 equiv.) and 4-ethylresorcinol (**2**, 628 mg, 3.27 mmol, 1.2 equiv.) were dissolved in methanesulfonic acid (6 mL) and stirred for 2 hours at 100 °C. The mixture was poured carefully to crashed ice (100 g), the formed solid orange precipitate was filtered off and dried. 1.12 g (95%) orange solid product (**6**) was formed and used without further purification.

^1^H NMR (400 MHz, DMSO-d6) δ 10.11 (bs, 1H), 8.42 (s, 1H), 8.29 (dd, J - 8.0, 1.5 Hz, 1H), 7.38 (d, J = 8.0 Hz, 1H), 6.74 (bs, 2H), 6.45 (bs, 2H), 2.36 (q, J = 7.5 Hz, 4H), 0.93 (t, J = 7.5 Hz, 6H). ^13^C NMR (101 MHz, DMSO-d6) δ 167.8, 166.1, 156.1, 150.3, 135.9, 132.7, 129.1, 127.6, 120.3, 102.3, 101.8, 22.4 14.1. HR-MS:(ESI) calcd. for C_25_H_21_O_7_ 433.12818 [M+H]^+^, found 433.12787, Df. = -0.7 ppm. HR-ESI-MS-MS (CID=65%; rel. int. %): 418(100); 405(25); 390(20) and 373(20).

#### Acylation of the fluoresceine derivative 6

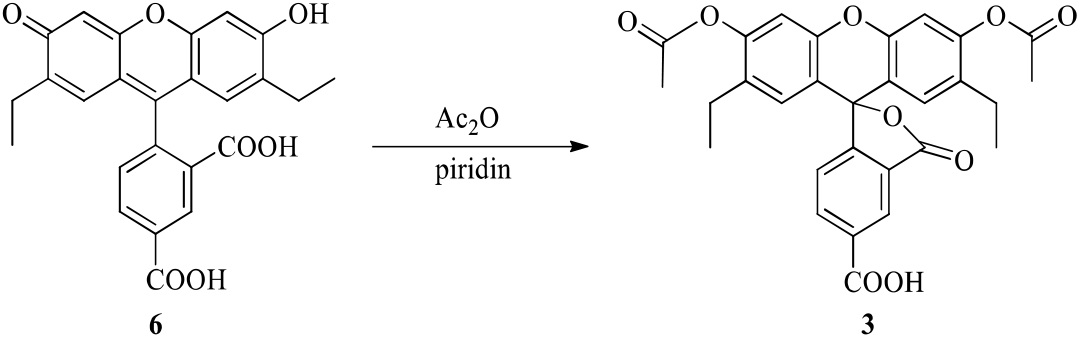

**6** (1.10 g, 2.54 mmol, 1.0 equiv.) was dissolved in acetic anhydride (3.50 mL, 3 equiv.) and pyridine (0.71 mL) was added than the mixture was stirred at 100 °C for 60 minutes. The mixture was cooled down to room temperature than poured carefully to crashed ice (100 g). The product was extracted with EtOAc (3 × 100 mL), the combined organic phases was washed with distilled water (2 × 200 mL), saturated sodium carbonate (2 × 200 mL) aqueous hydrochloric acid (1M, 1 × 50 mL), distilled water (1 × 200 mL) and brine (1 × 100 mL). The solvent was dried over MgSO_4_, evaporated under reduced pressure. 621 mg (47%) product (**3**) was isolated as a yellow solid.

^1^H NMR (400 MHz, DMSO-d6) δ 13.63 (bs, 1H), 8.27 (dd, J = 7.9, 1.2 Hz, 1H), 8.18 (dd, J = 7.9, 0.8 Hz, 1H), 7.78 (dd, J = 1.2, 0.8 Hz, 1H), 7.25 (s, 2H), 6.76 (s, 2H), 2.38 (m, 4H), 2.33 (s, 6H), 0.92 (t, J = 7.5 Hz, 6H). ^13^C NMR (101 MHz, DMSO-d6) δ 168.9, 167.7,166.0,152.4,150.3,148.9,137.5,132.3, 131.3, 128.9, 128.1, 125.7, 124.5, 115.8, 111.2, 81.6, 22.1, 20.6, 14.3. HR-MS:(ESI) calcd. for C_29_H_26_O_9_ 517.14931 [M+H]^+^, found 517.14944.

#### *N*-acylation of amino BAPTA moiety using 3

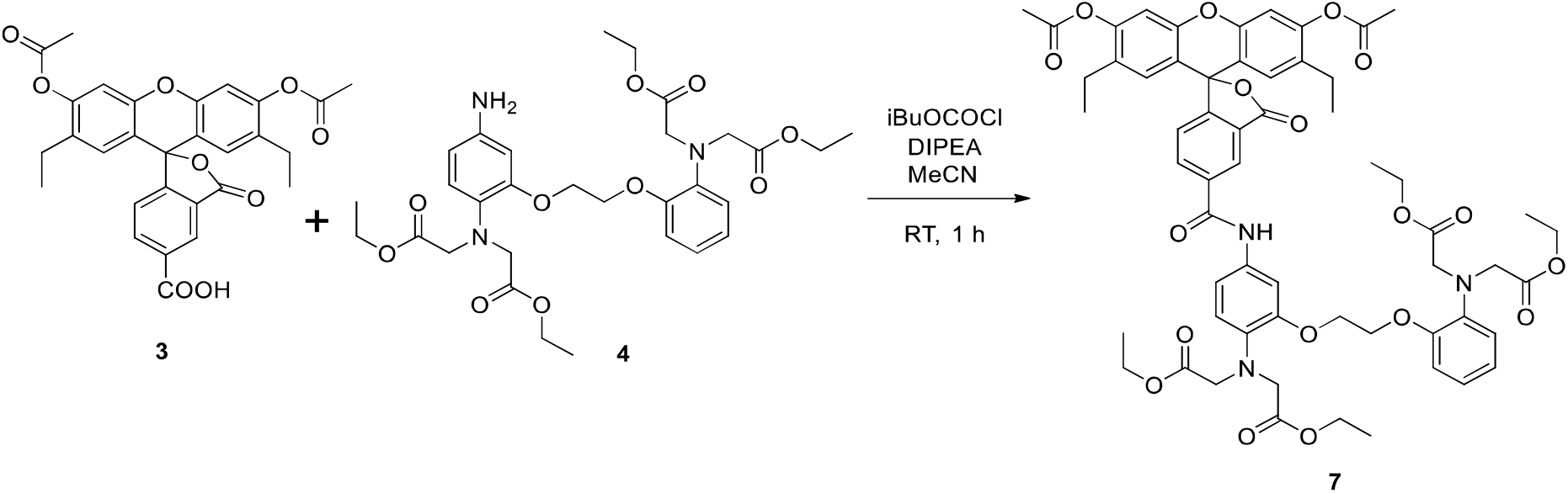

Tetraethyl ester **3** (156 mg, 0.302 mmol, 1.0 equiv.) was dissolved in dry dichloromethane (5 mL) diisopropylethylamine (0.049 mL, 0.605 mmol, 2.0 equiv.) was added. The mixture was cooled down to 0°C. Isobutyl chloroformate (0.038 mL, 0.29 mmol, 0.95 equiv.) was added, the mixture was stirred at room temperature for 60 minutes. 219 mg (0.363 mmol, 1.2 equiv.) of amino BAPTA derivative (**4**)^1^ was added, the mixture was stirred overnight. The mixture was poured into distilled water (40 mL), the product was extracted with EtOAc (6 × 50 mL), the combined organic phases was washed with saturated sodium hydrogen carbonate (1 × 50 mL) and brine (1 × 100 mL). The solvent was dried over MgSO_4_, evaporated under reduced pressure. The residue (405 mg) was purified by preparative HPLC (water-acetonitrile-0.1% TFA, using the gradient method). After purification, the fractions were lyophilized. The product (**7,** 170 mg, yield 51%) was isolated as a yellow powder.

^1^H NMR (400 MHz, CDCh) δ 8.69 (bs, 1H), 8.56 (bs, 1H), 8.58 (s, 1H) 8.29 (dd, *J* = 8.0, 1.7 Hz, 1H), 8.21 (m, 1H), 8.11 - 8.01 (m, 5H), 7.54 - 7.49 (m, 1H), 7.36 - 7.27 (m, 2H), 7.03 (s, 1H), 6.91 - 6.82 (m, 2H), 6.62 (s, 1H), 4.15 - 4.13 (m, 4H), 4.67 (s, 8H), 4.16 (qd, *J* = 7.4 Hz, 3.0 Hz, 8H), 2.45 - 2.36 (m, 4H), 2.36 - 2.32 (m, 6H), 1.25 - 1.12 (m, 12H), 1.01 (t, *J* = 7.5 Hz, 6H). ^13^C NMR (101 MHz, CDCh) δ 172.0, 171.7, 169.1, 168.8, 165.6, 165.6, 155.7, 150.6, 150.5, 150.4, 149.8, 137.4, 135.0, 134.3, 133.8, 133.8, 132.4, 131.4, 130.4, 129.9, 129.8, 129.0, 128.2, 126.7, 124.9, 121.7, 119.4, 119.3, 115.9, 113.2, 111.3, 63.2, 63.1, 61.2, 61.1, 53.9, 23.2, 21.0, 14.3, 14.2, 14.2. HR-MS:(ESI) calcd. for C_59_H_64_N_3_O_18_ 1102.41794 [M+H]^+^, found 1102.41960. HR-ESI-MS-MS (CID=40%; rel. int. %): 1060(25); 1028(100) and 986(8).

#### Hydrolysis of 7 using aqueous sodium hydroxide

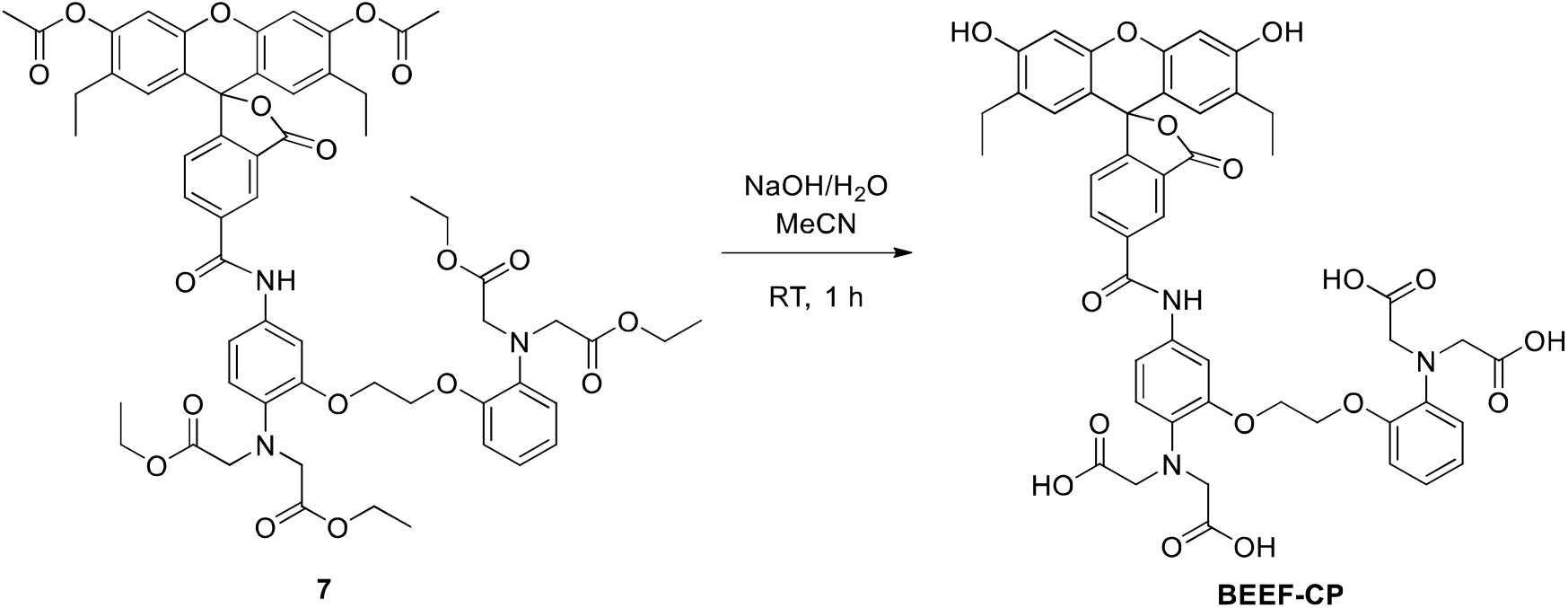

**7** (130 mg, 0.118 mmol, 1.0 equiv.) was dissolved in methanol (2 mL). Sodium hydroxide (50 mg, 125 mmol, 11 equiv.) in water (0.40 mL) was added at 0°C, the mixture was stirred at room temperature for 60 minutes. The mixture was diluted to 5 mL with ethanol/acetonitrile 1/1 and was purified by preparative HPLC (water-acetonitrile-0.1% TFA, using the gradient method). After purification, the combined fractions were lyophilized. The product (**BEEF-CP,** 88 mg, yield 82%) was isolated as a yellow powder.

^1^H NMR (600 MHz, DMSO-d·.) δ 12.38 (bs, 2II), 10.41 (s, 111), 10.12 (bs, 2H), 8.58 (s, 1H) 8.32 (dd, *J* = 8.0, 1.7 Hz, 1H), 7.46 (d, *J* = 2.3 Hz, 1H), 7.40 (d, *J* = 8.1 Hz, 1H), 7.37 - 7.33 (m, 1H), 7.02 - 7.00 (m, *J* = 9.6 Hz, 1H), 6.89 - 6.83 (m, 2H), 6.80 (d, *J* = 8.8 Hz, 1H), 6.78 - 6.75 (m Hz, 1H), 6.72 (s, 2H), 6.40 (s, H), 4.32 - 4.26 (m, 4H), 4.06 (s, 8H), 2.37 (ddp, *J* = 21.7, 14.7, 7.4 Hz, 4H), 0.94 (t, *J* = 7.5 Hz, 6H). ^13^C NMR (151 MHz, DMSO-_de)_ δ 172.4, 168.2, 163.4, 158.2, 158.0, 145.0, 149.5, 149.5, 139.4, 136.6, 135.6, 132.8, 127.3, 121.5, 121.2, 118.5, 118.4, 115.0, 113.5, 108.6, 107.5, 101.8, 67.2, 67.1, 53.5, 22.4, 14.2. HR-MS:(ESI) calcd. for C_47_H_44_N_3_O_16_ 906.27161 [M+H]^+^, found 906.27247. HR-ESI-MS-MS (CID=40%; rel. int. %): 848(100); 816(21); 728(9); 593(9) and 433(3).

### Spectroscopic characterization of BEEF-CP

#### One-photon absorption and emission of BEEF-CP

**UV-Vis absorption spectra** in the wavelength range of 220–700 nm were recorded on a Thermo Scientific Evolution 220 spectrometer in a quartz cuvette (pathlength = 1.0 cm). The general sample preparation protocol involved the dissolution of 1–3 mg of the studied compounds in 50.0 mL MOPS buffer (pH = 7.20) and then a dilution to the suitable concentrations (1.0–8.8× 10^−5^ M) based on the UV-Vis absorption properties of the compounds. The spectrum of the pure solvent was subtracted from the sample spectra. Using the corrected spectra, the molar absorbance (ε) at a specific wavelength value was calculated, and the position of a selected absorption band (*λ*_max_) and the full molar absorption spectra of the compounds were determined.

**Fluorescence emission data** were measured on a Hitachi F-4500 spectrophotometer in a quartz cell with 1.0 cm pathlength. Slit widths were selected to provide 5 nm and 10 nm bandpass for the excitation and emission beams, respectively. Emission data were normalized to the intensity of a probe body, as an external reference, recorded every day. Solvent background spectra (emission spectra or excitation scans) were subtracted from the spectra of samples. Sample solutions were prepared by dilutions from the stock solutions used for the recording of the UV/Vis absorption spectra. The final concentration of the solutions prepared for fluorescence experiments was around 5 μΜ. The compounds were excited at their absorption maxima (*λ*_max_) determined during UV/Vis absorption measurements. Quantum yields were calculated by ratiometric method^2^.

**Figure S13.**
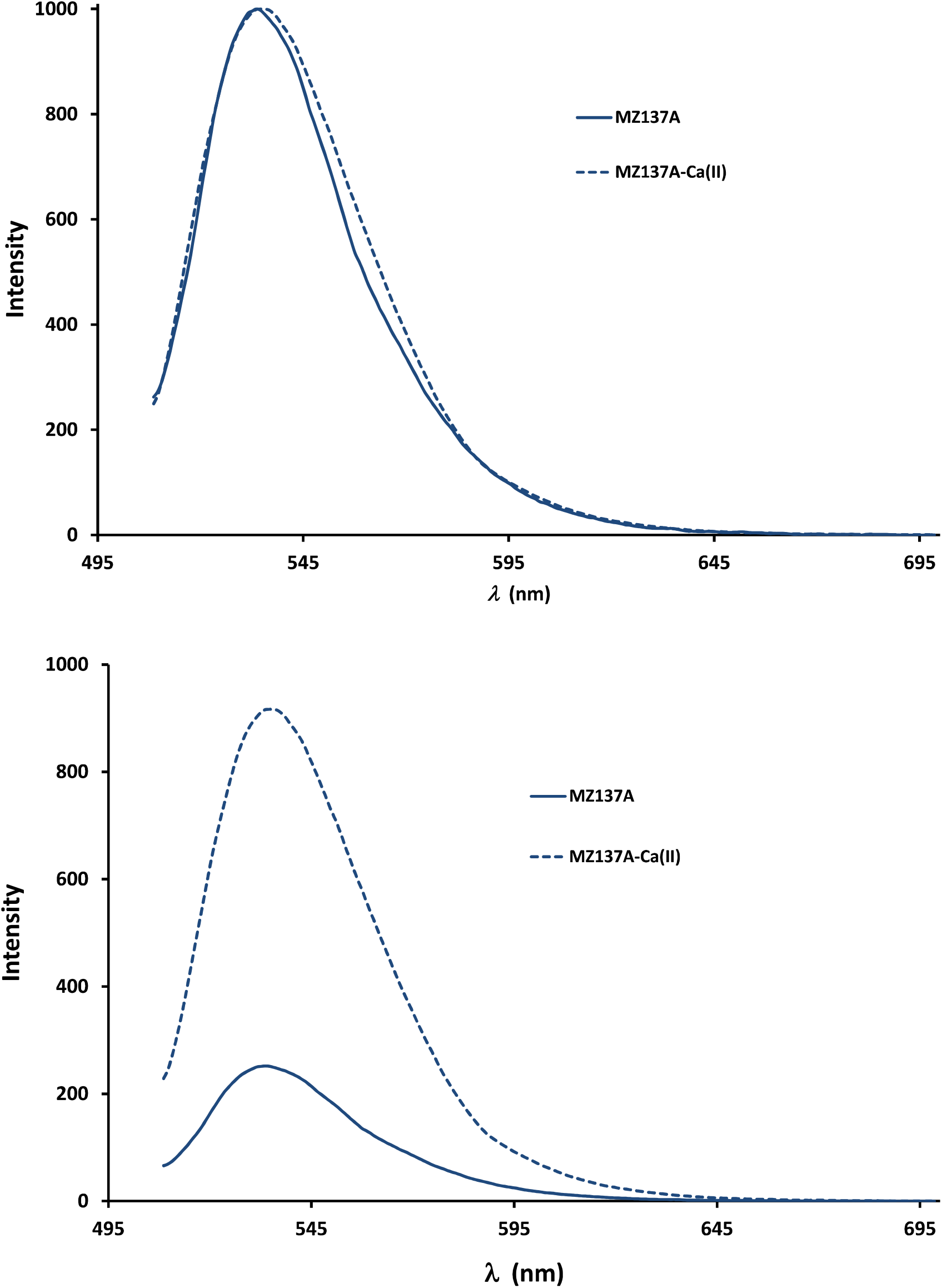
**Comparison of the molar UV-Vis absorption (up) and fluorescence emission (down) spectra** (normalized to c = 1.0 μΜ concentrations) of BEEF-CP in water using MOPS buffer (pH = 7.20) and 100 mM KCl in the presence of EGTA (10.0 mM) without calcium ion (solid line) or in the presence of Ca^2+^ (10.0 mM) (dashed line). Excitation was carried out at or near the absorption maxima.

**Table S5.** Spectroscopic parameters of **BEEF-CP** without Ca^2+^ or in the presence of calcium ion in aqueous media.

| | Absorption | | Emission | | $QY^a$ | $I_{EM,max} - I_{EX}$<br>(nm) |
| --- | --- | --- | --- | --- | --- | --- |
| | $I_{max}$<br>(nm) | $\epsilon$ ($\text{M}^{-1}\text{cm}^{-1}$) | $I_{EM,max}$ (nm) | $I_{max}$ (1<br>mM) | | |
| **BEEF-CP**<sup>b</sup> | 504 | 58600 | 534 | 252 | 0.141 | 30 |
| **[BEEF-CP – $\text{Ca}^{2+}$]**<sup>c</sup> | 504 | 57900 | 535 | 917 | 0.548 | 31 |

<sup>a</sup>Quantum yields were determined using fluorescein in HEPES buffer (pH = 7.4) as a reference.<sup>b</sup>The measurements are carried out in water using MOPS buffer (pH = 7.20) and 100 mM KCl in the presence of EGTA (10.0 mM).<sup>c</sup>The measurements are carried out in water using MOPS buffer (pH = 7.20) and 100 mM KCl in the presence of EGTA (10.0 mM) and $\text{CaCl}_2$ (10.0 mM).

#### Determination of dissociation constant (K_d_) of fluorescent calcium indicator BEEF-CP

Fluorescence-monitored Ca^2+-^titrations were executed by using a solution of the selected dye molecule containing the Ca^2+^-buffer EGTA in a concentration of 0.01 M but no Ca^2+^ ions. These starting solutions were made by the dilution of the stock solution of the dye into ‘Buffer A’. Another solution, containing the dye in the same concentration, was also prepared by using ‘Buffer B’. This latter solution contained the Ca^2+^ buffer EGTA and Ca^2+^ ions at identical, 0.01 M concentration, setting the free Ca^2+^ level to 37 μΜ. For the measurement of the first (0 M) point of the titration curve 2.0 mL of the sample made with ‘Buffer A’ was used. In the titration process various (typically increasing) volume fractions of the sample were replaced by the same volumes of the solution made with ‘Buffer B’. Fine details of this fluorescence titration protocol have been described previously.^3,4^ The recorded spectra are shown in Fig. S14 and the fluorescence intensities versus the log[Ca^2+^]_free_ are shown in Fig. S15. The K_d_ value of **BEEF-CP** determined using this method is 175.6 nM.

**Figure S14.**
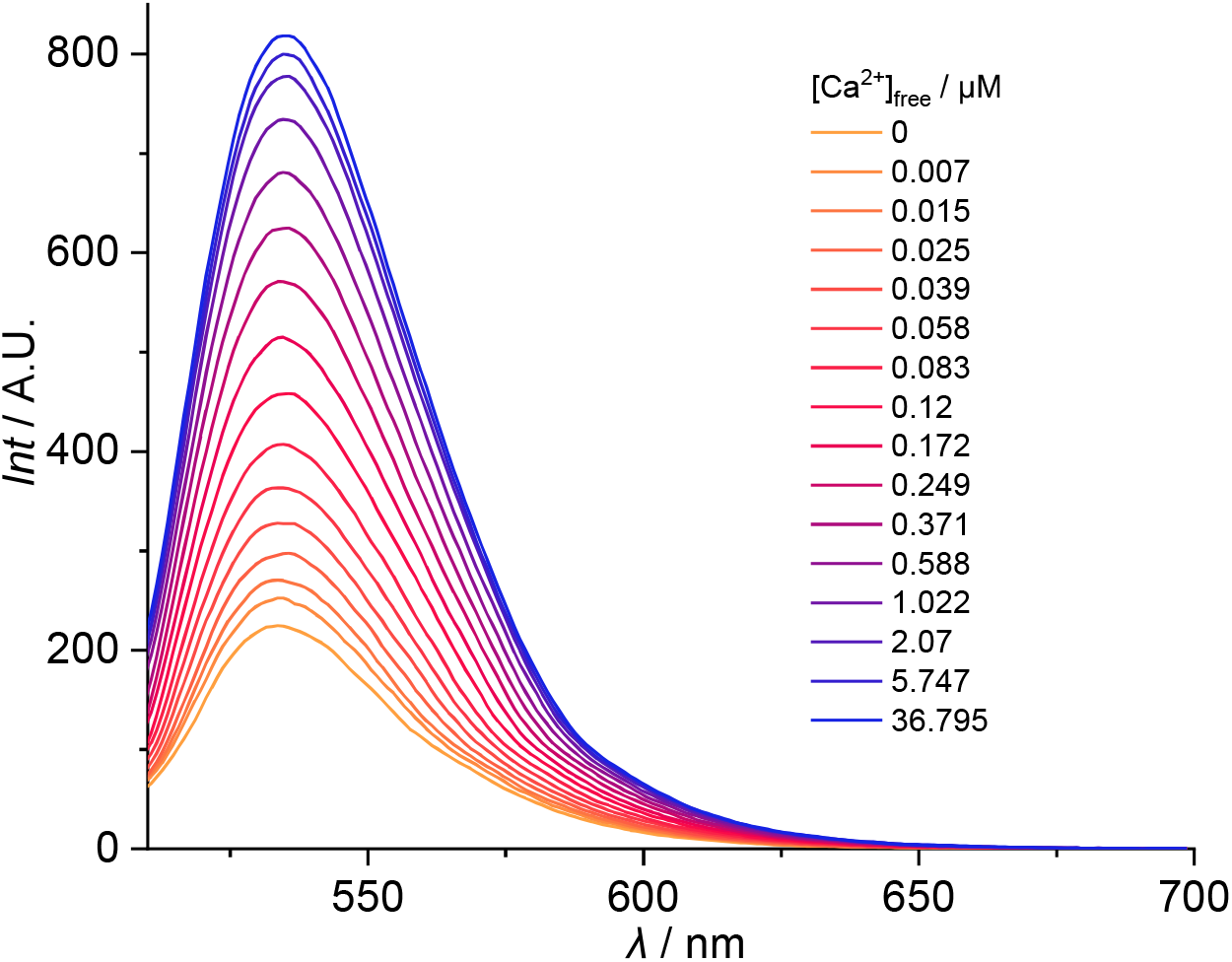
**Emission spectra of BEEF-CP** recorded in the presence of different free Ca^2+^ concentrations in MOPS buffer (pH = 7.2) containing 100 mM KCl.

**Figure S15.**
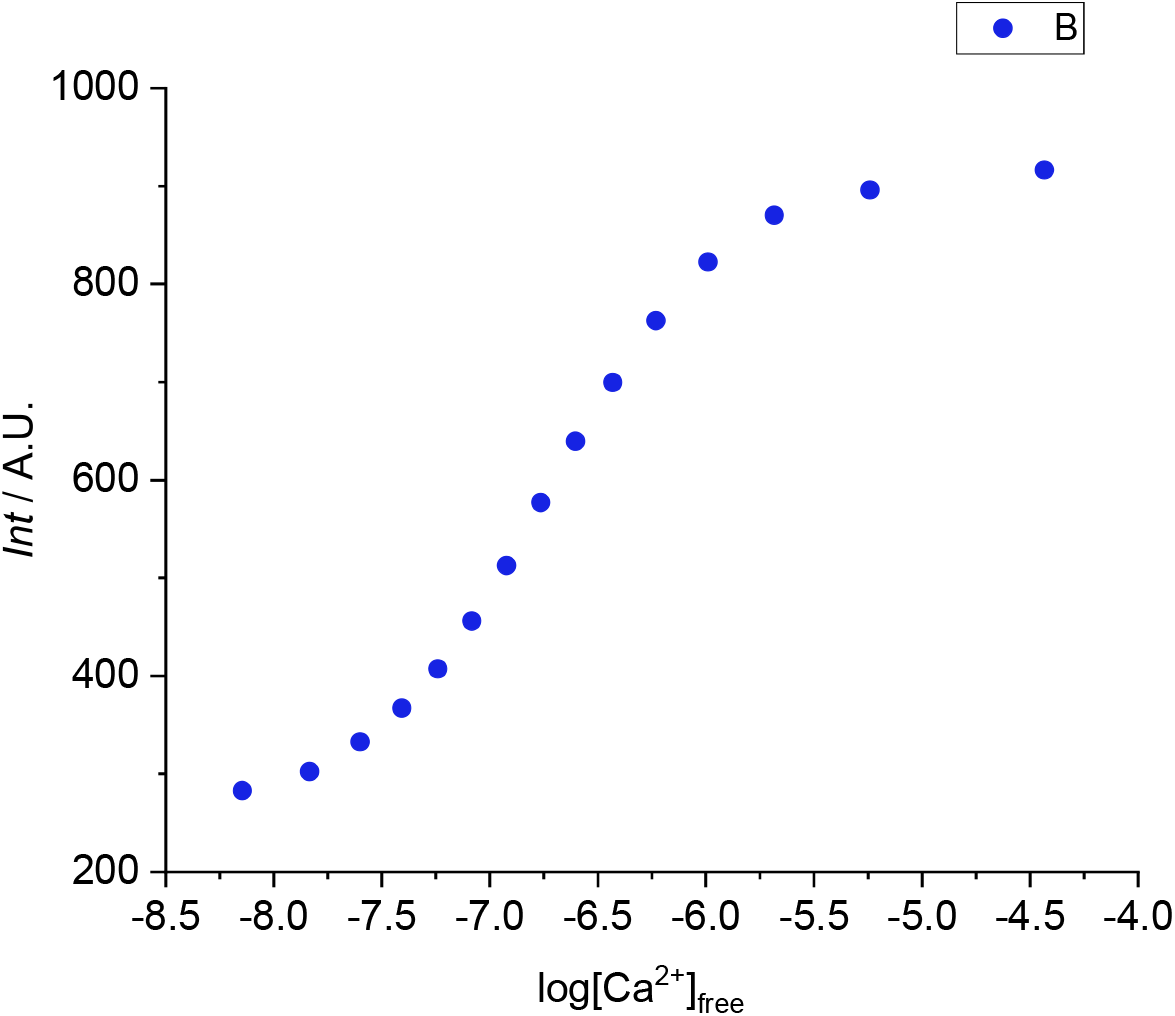
**Calcium titration curves** used for the determination of the dissociation constant (K_d_) constant of the calcium complex of compound BEEF-CP.

#### pH sensitivity of the fluorescent sensor (BEEF-CP)

0.8 mg of **BEEF-CP** was dissolved in DMSO to obtain a stock solution. pH Buffers for the range of 3-9 based on NaOAc/AcOH (pH 3-5) or Na-HEPES/HEPES (pH 6-9) were prepared in the concentration of 5 mM. The ionic strength was set to 50 mM by the addition of KCl and either 1.5 mM EGTA or 1.5 mM CaCl_2_ was dissolved in the solution. The fluorescent spectra of **BEEF-CP** has been recorded using a Shimadzu 1900 spectrophotometer in each Ca^2+^-containing and Ca^2+^-free pH buffer. The used excitation wavelength was 504 nm, both the excitation and emission slits were set to 5 nm. The spectra were recorded at a scan speed of 2000 nm/min by recording datapoints every in 0.5. nm. The instrument was used in low sensitivity mode. The emission intensity values were plotted against for pH in both the Ca^2+^-containing and Ca^2+^-free solutions shown in Fig. S16.

**Figure S16.**
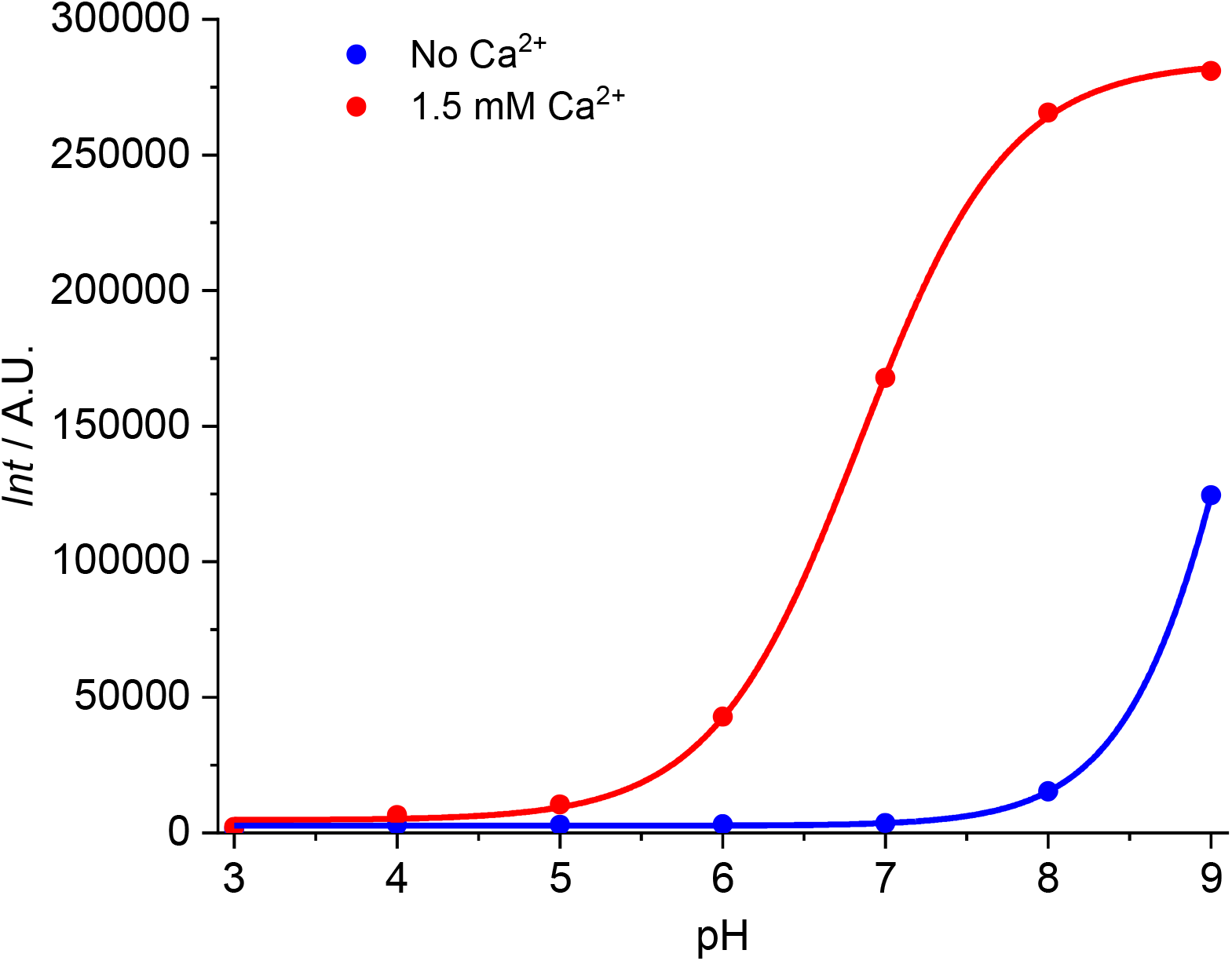
**Emission intensity of probe BEEF-CP** measured in solutions containing no Ca^2+^ or 1.5 mM Ca^2+^ at different pH values.

#### Two-photon cross-section measurements

**The two-photon absorption cross section (TPCS)** was determined with the two-photon excited fluorescence (TPEF) method^5^. The measurements were performed using an inverted two-photon microscope (FemtoSmart2D, Femtonics), equipped with a XLUMPFLN20XW Olympus objective (numerical aperture; NA = 1.0) and a tunable high-power Ti:Sapphire laser (Coherent Chameleon Discovery COM5, wavelength of the excitation light is between 700 nm and 1040 nm). The incident light source was focused into at capillary filled with either the sample or the reference solution (Rhodamine 6G in MeOH^6^) and integrated fluorescence emission was detected in a wavelength window from 475 to 575 nm (green channel of the microscope). The power of the laser source was kept constant at 15 mW. TPCS at each excitation wavelength was calculated according to the following equation:

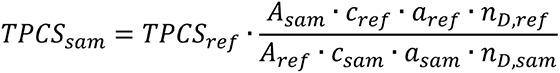

where *A* is the mean TPEF emission intensity, *c* is the concentration of the dye, *n* is the refractive index of the solvent measured at the sodium D-line; *a* is a ratio derived from one-photon emission measurements calculated as the integral of one-photon emission spectrum from 475 to 575 nm divided by the total one-photon emission spectrum integral, *ref* is reference, *sam* is sample.

**Figure S17.**
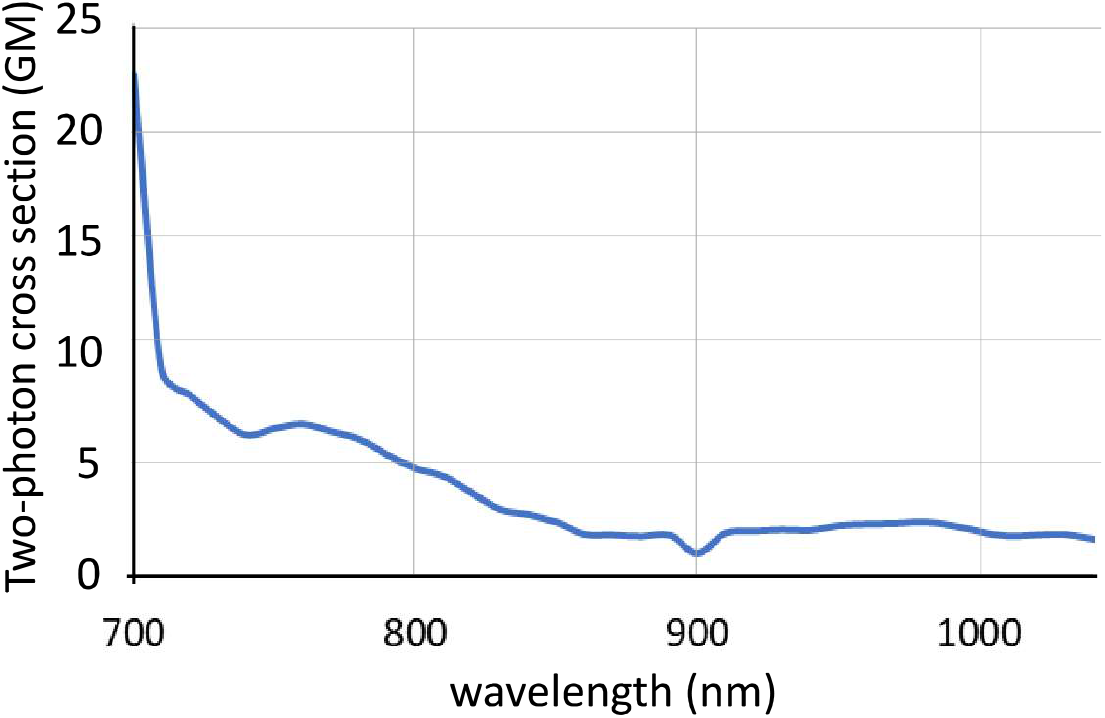
**Two-photon cross section of BEEF-CP** in HEPES buffer at pH = 7.2. GM: Goeppert-Mayer unit.

### NMR spectra of the synthesized novel compounds

**Figure S18.**
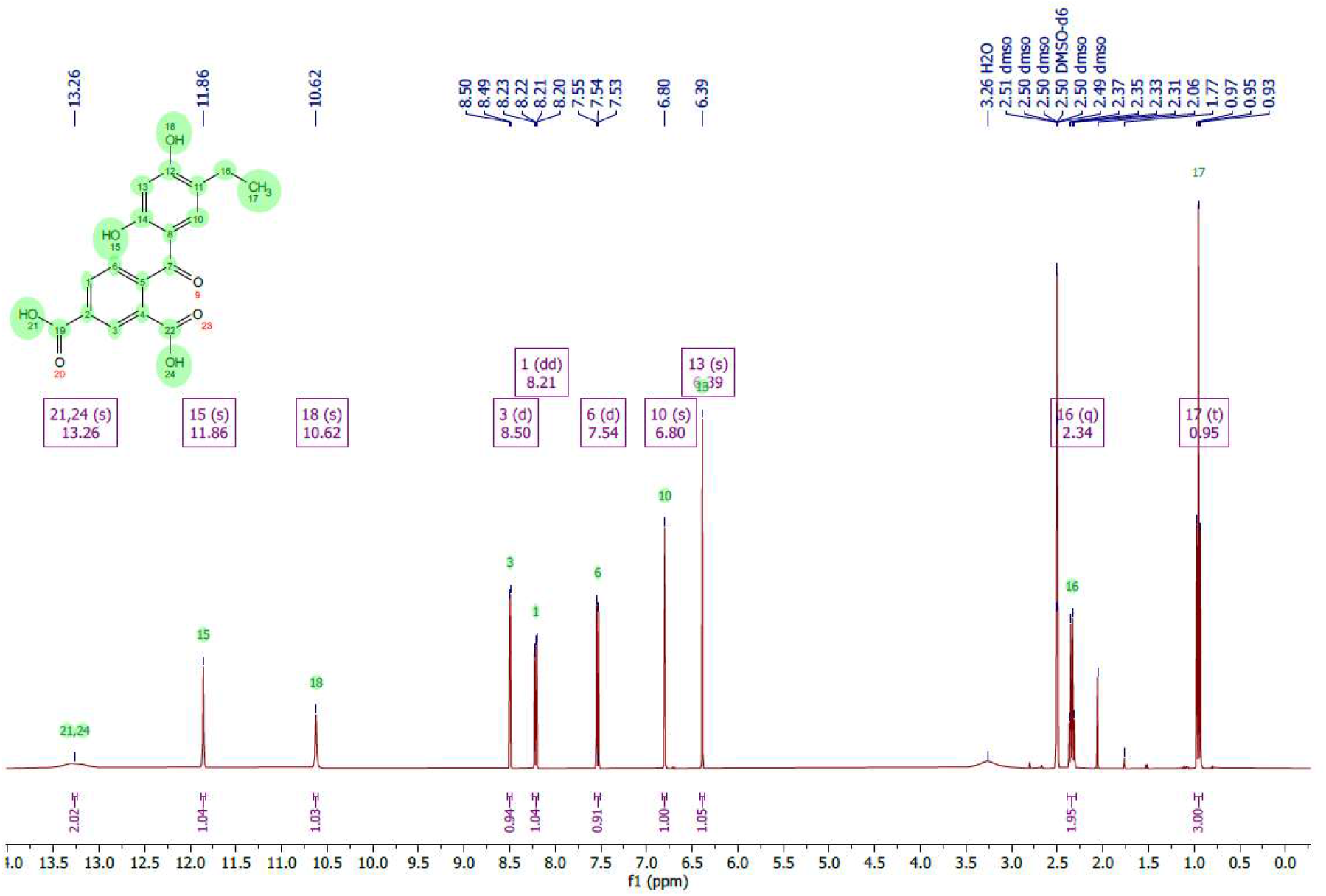
^1^H NMR spectrum of **5** recorded at 400 MHz in DMSO-d_6_.

**Figure S19.**
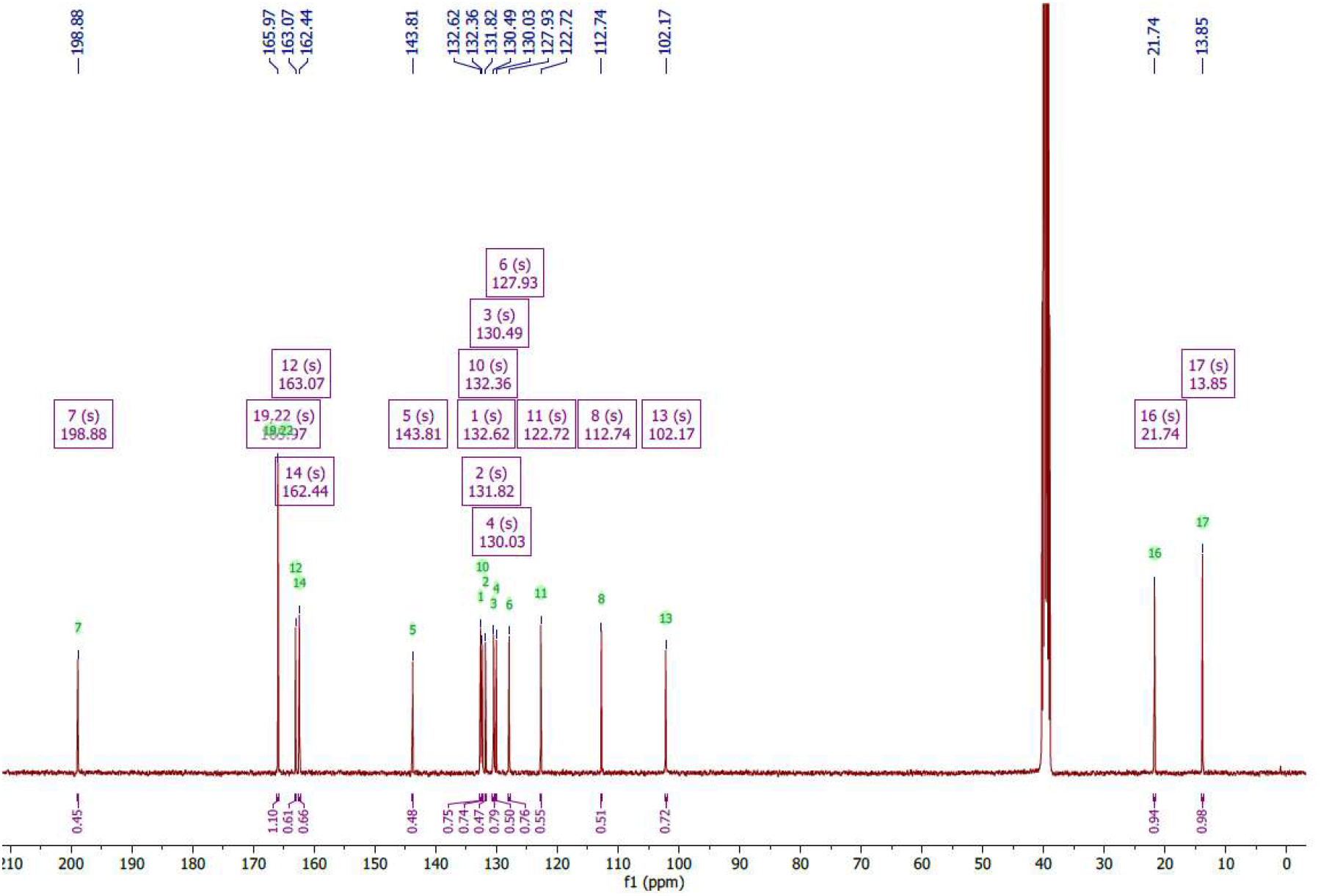
^13^C NMR spectrum of **5** recorded at 101 MHz in DMSO-d_6_.

**Figure 20.**
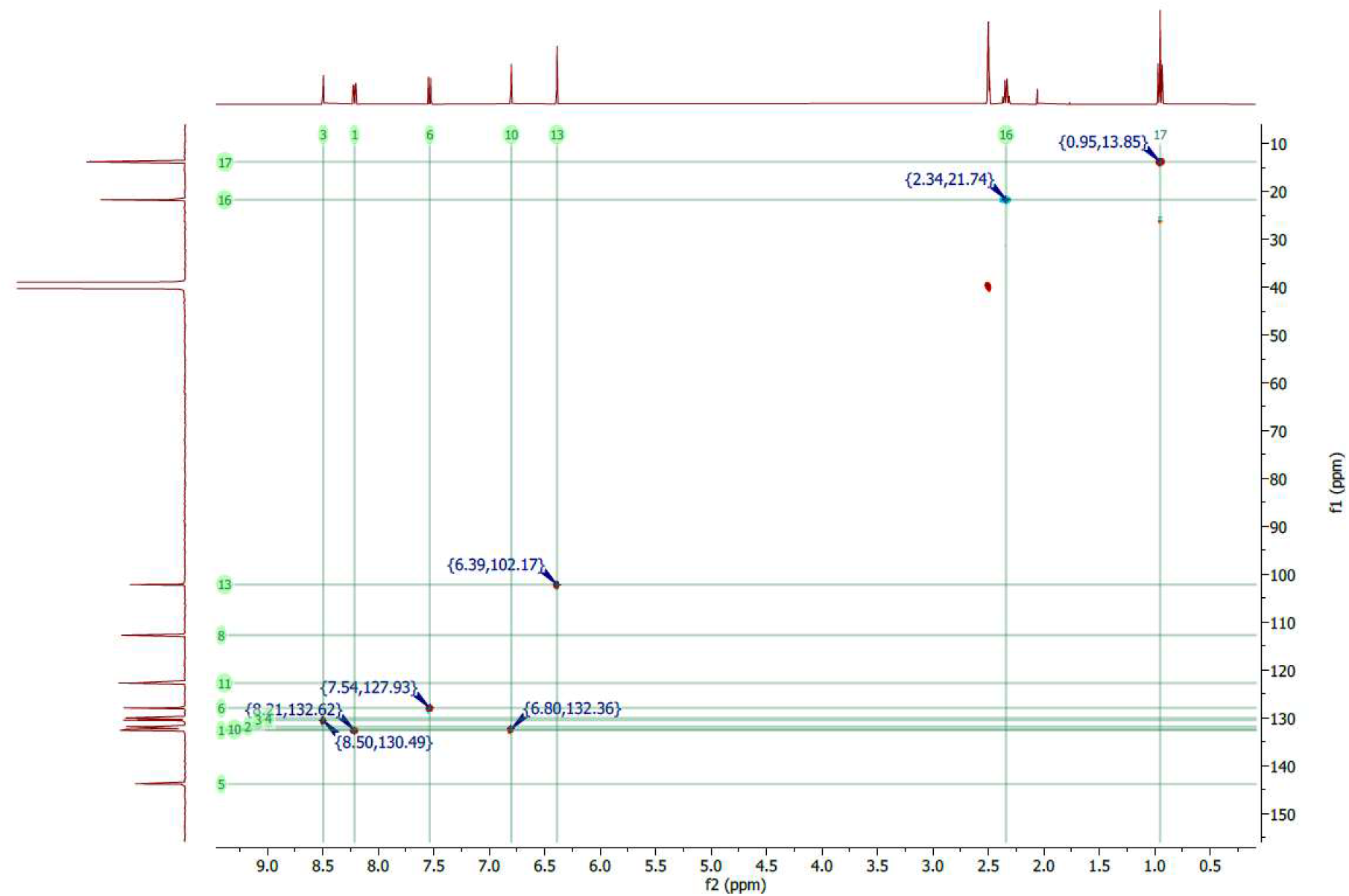
HSQC spectrum of **5** recorded in DMSO-d_6_.

**Figure S21.**
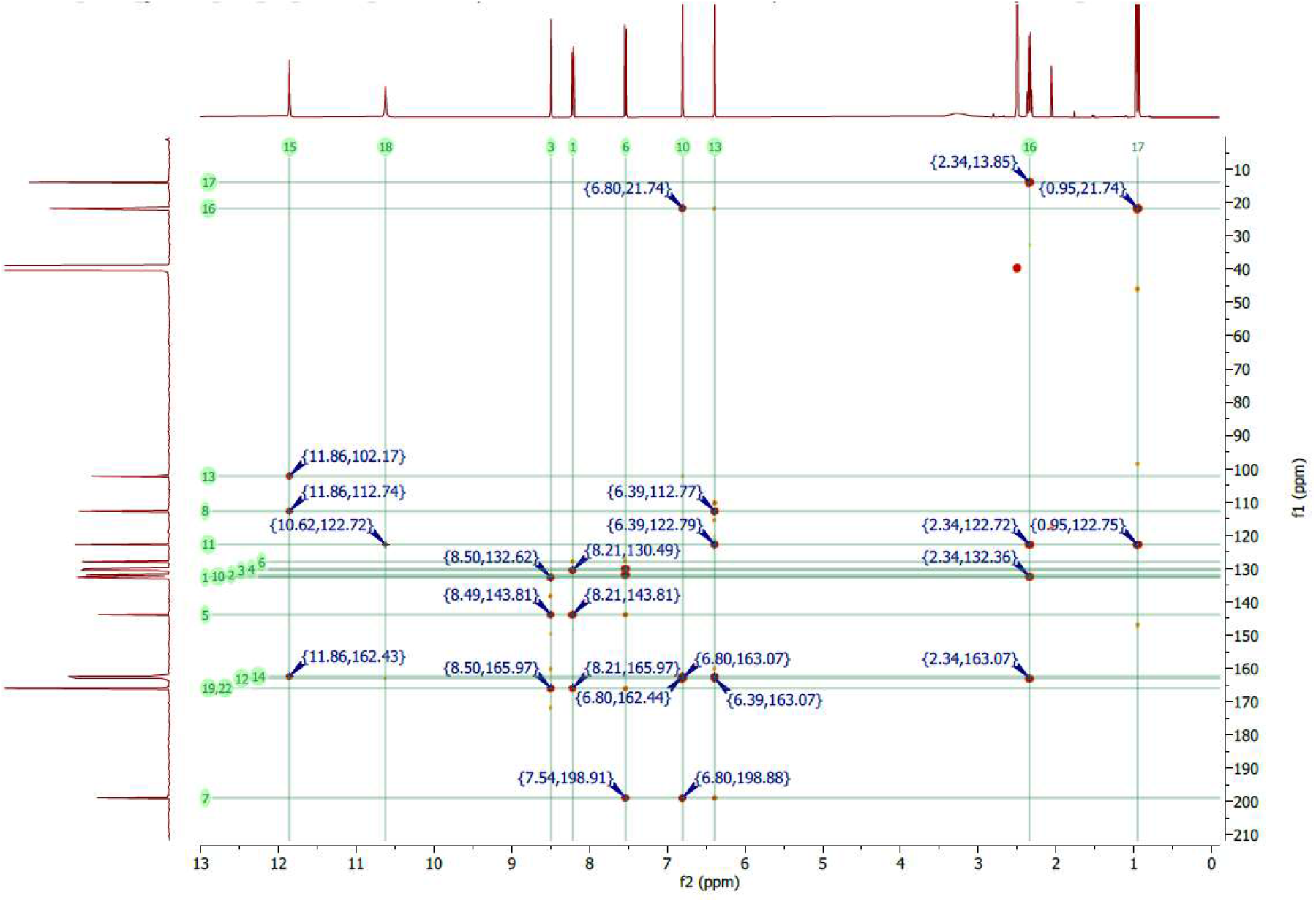
HMBC spectrum of **5** recorded in DMSO-d_6_.

**Figure S22.**
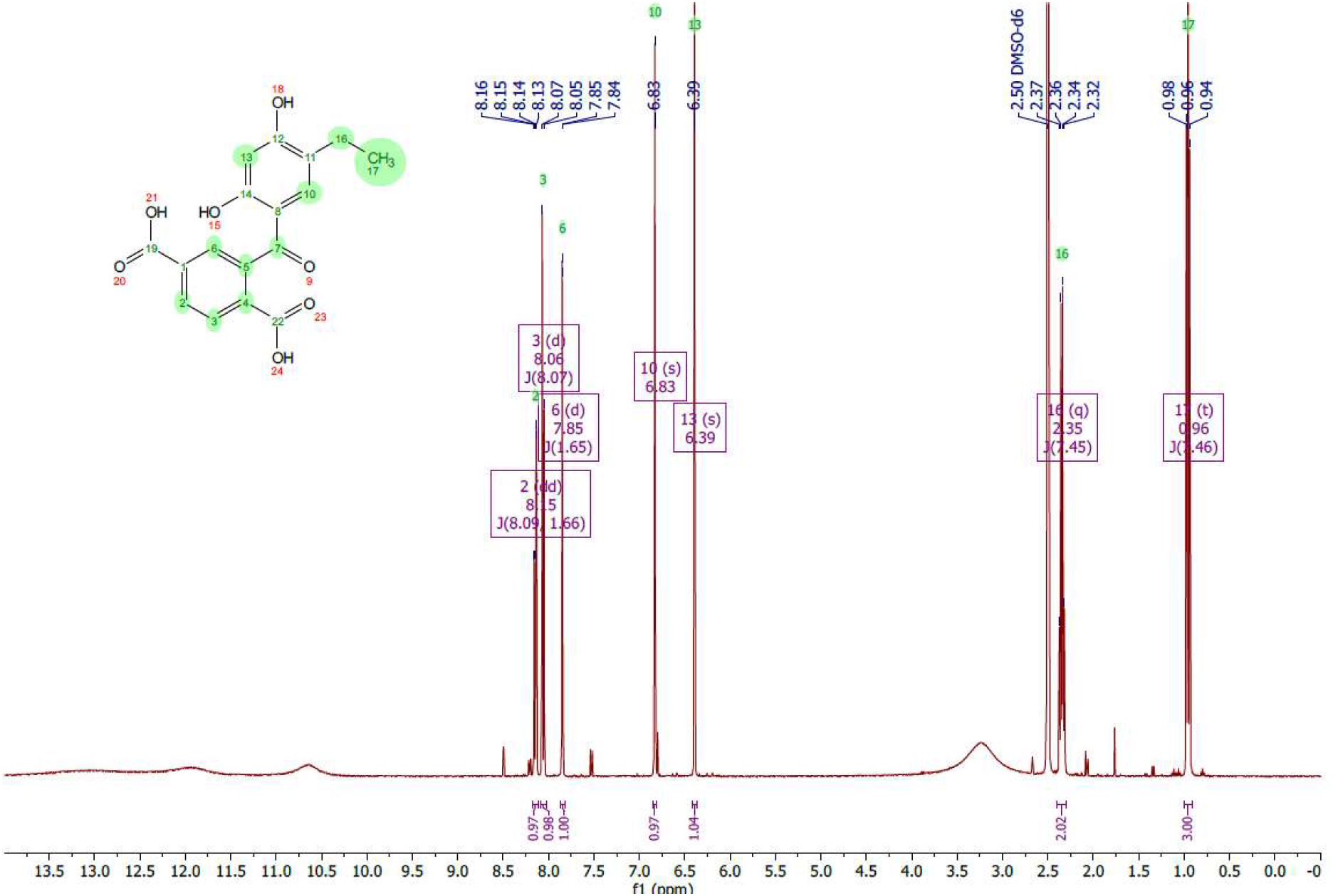
^1^H NMR spectrum of **5_co** recorded at 400 MHz in DMSO-d_6_.

**Figure S23.**
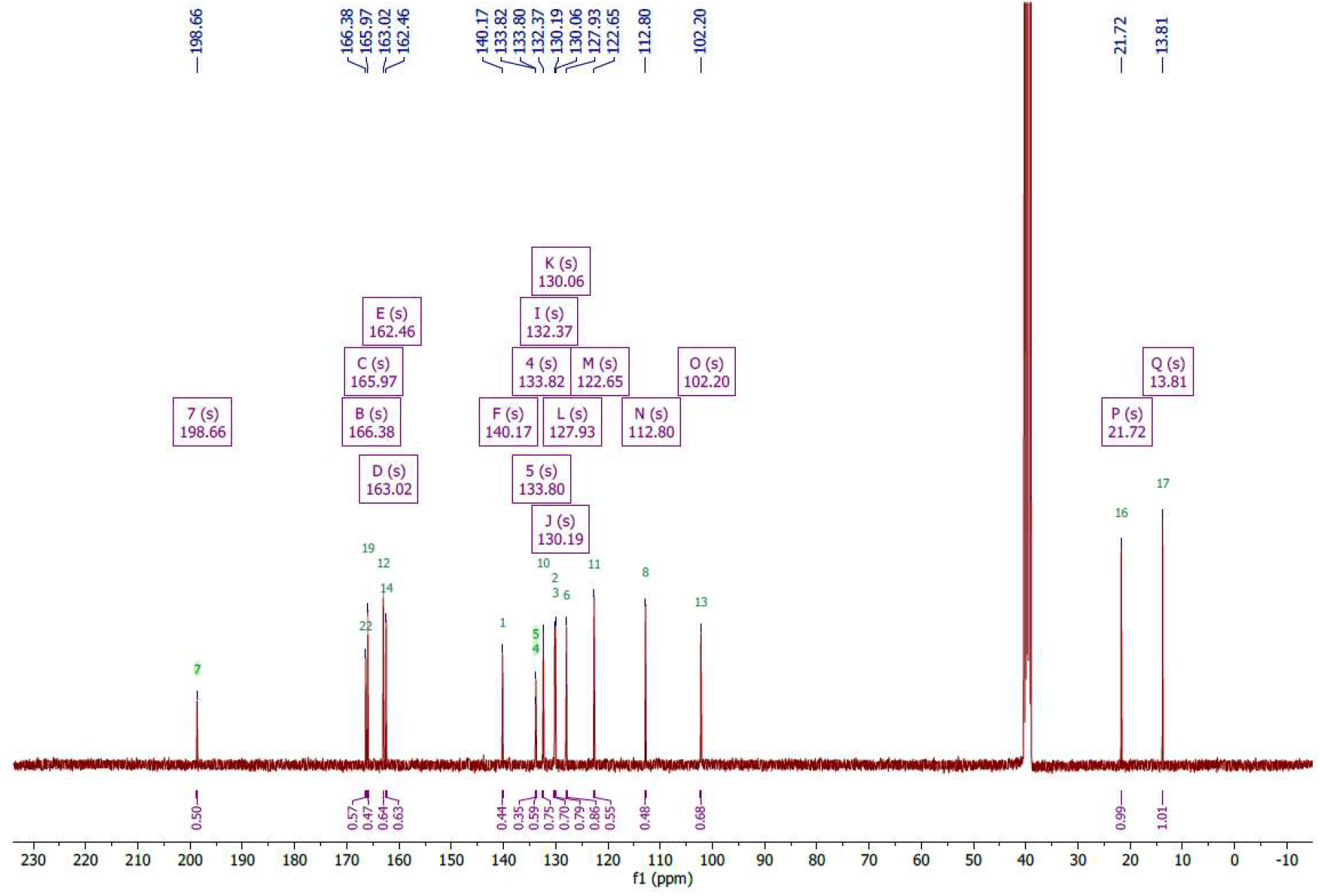
^13^C NMR spectrum of **5_co** recorded at 101 MHz in DMSO-d_6_.

**Figure S24.**
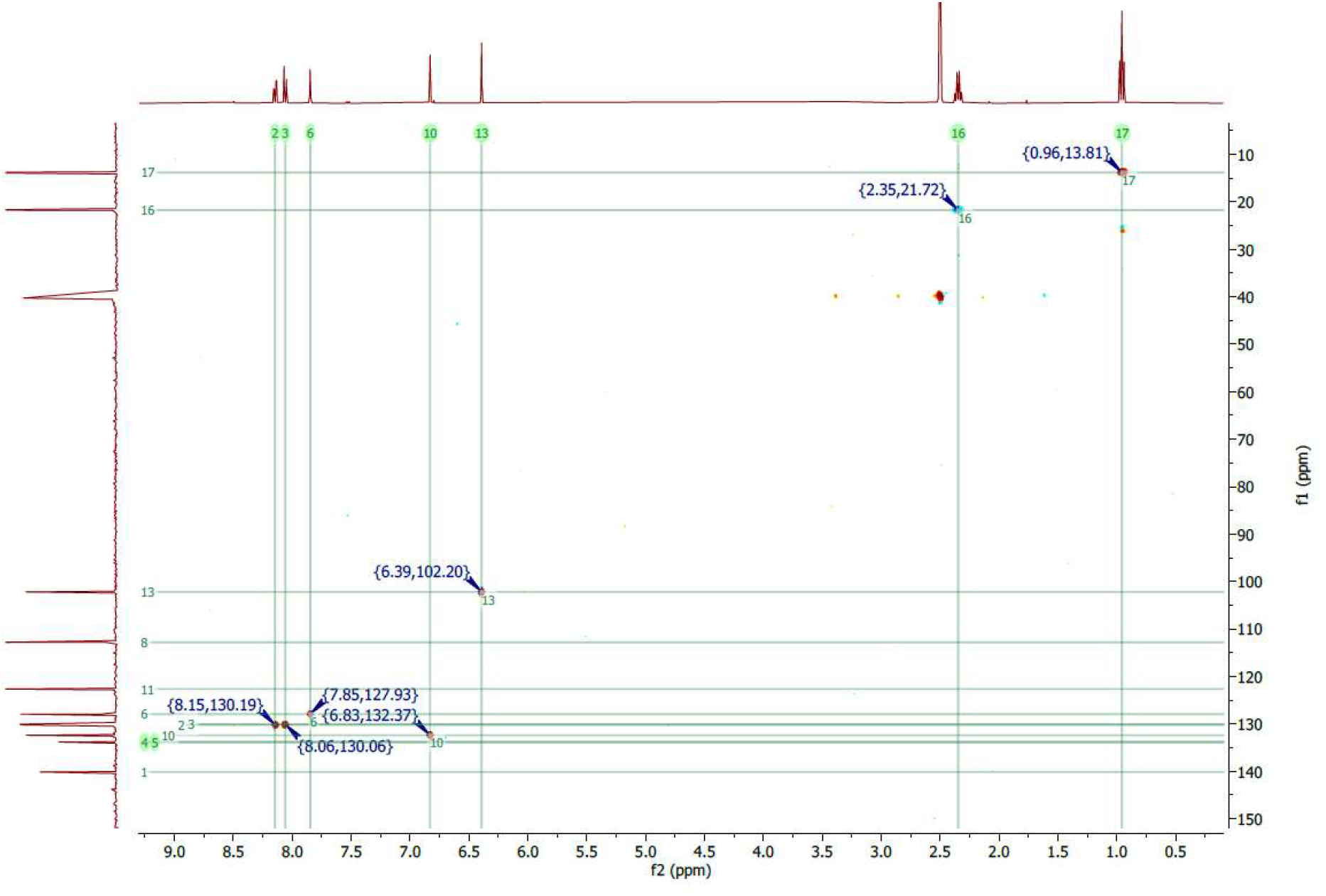
HSQC spectrum of **5_co** in DMSO-d_6_.

**Figure S25.**
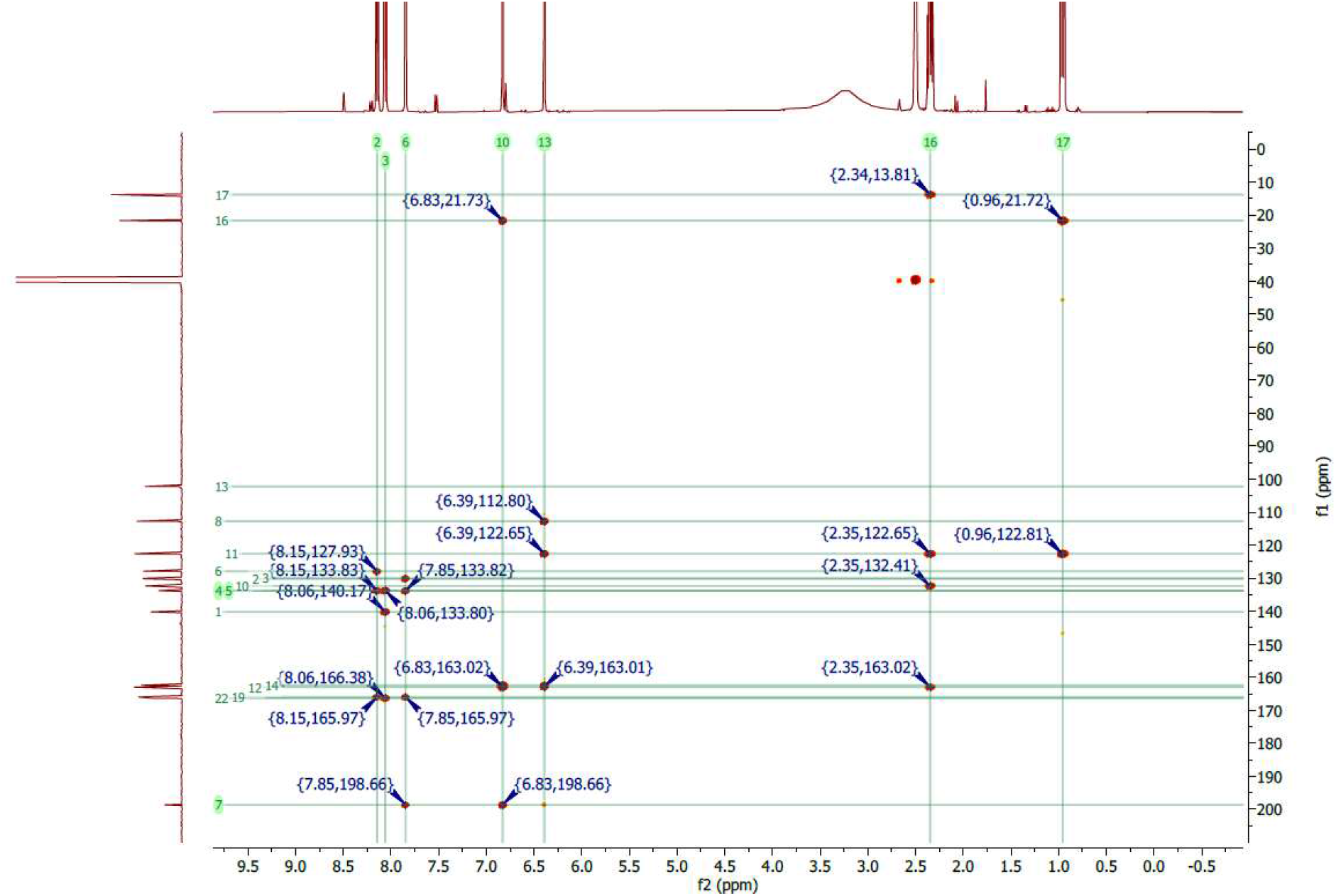
HMBC spectrum of **5_co** in DMSO-d_6_.

**Figure S26.**
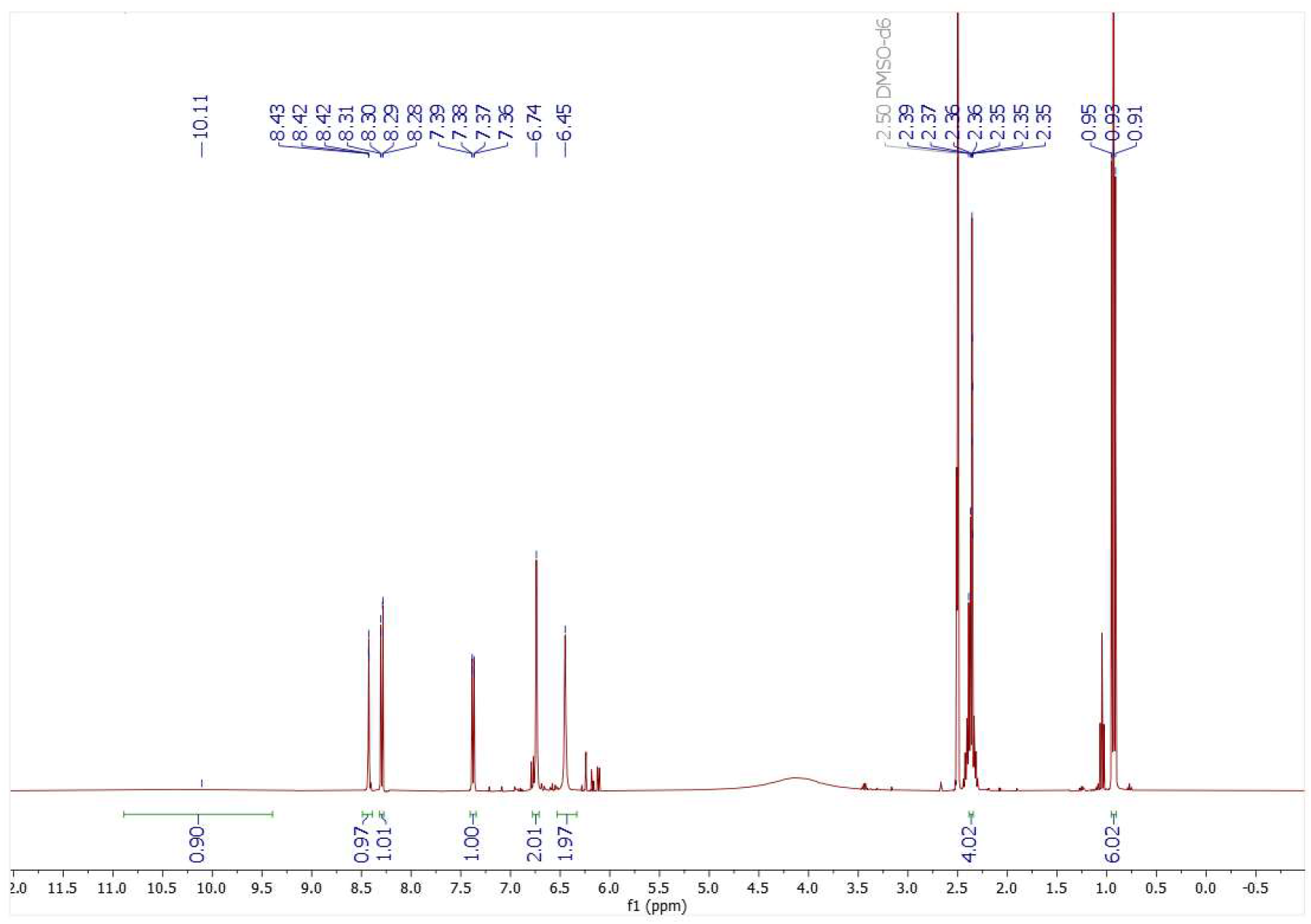
^1^H NMR spectrum of **6** recorded at 400 MHz in DMSO-d_6_.

**Figure S27.**
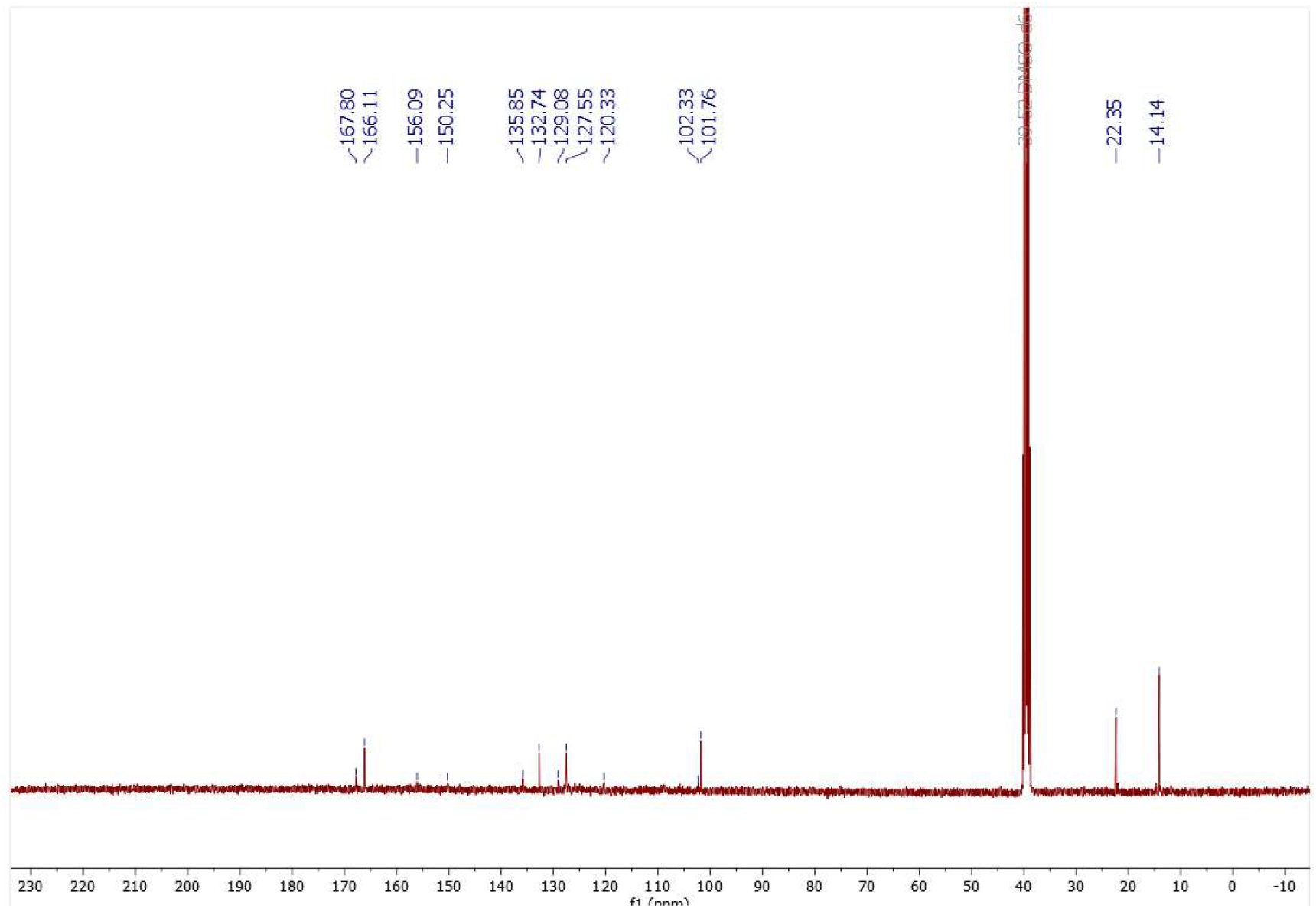
^13^C NMR spectrum of **6** recorded at 101 MHz in DMSO-d_6_.

**Figure S28.**
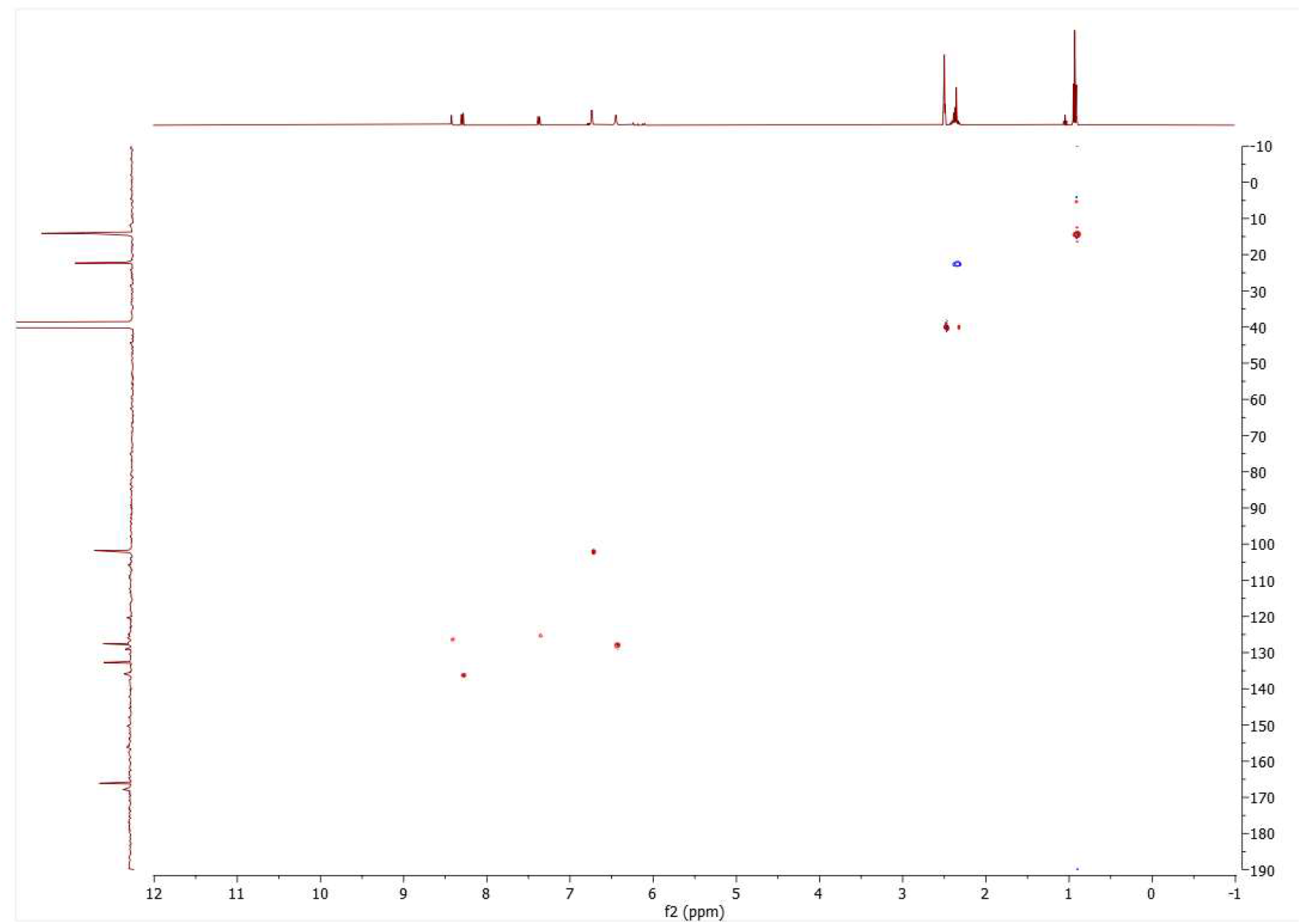
HSQC spectrum of **6** recorded in DMSO-d_6_.

**Figure S29.**
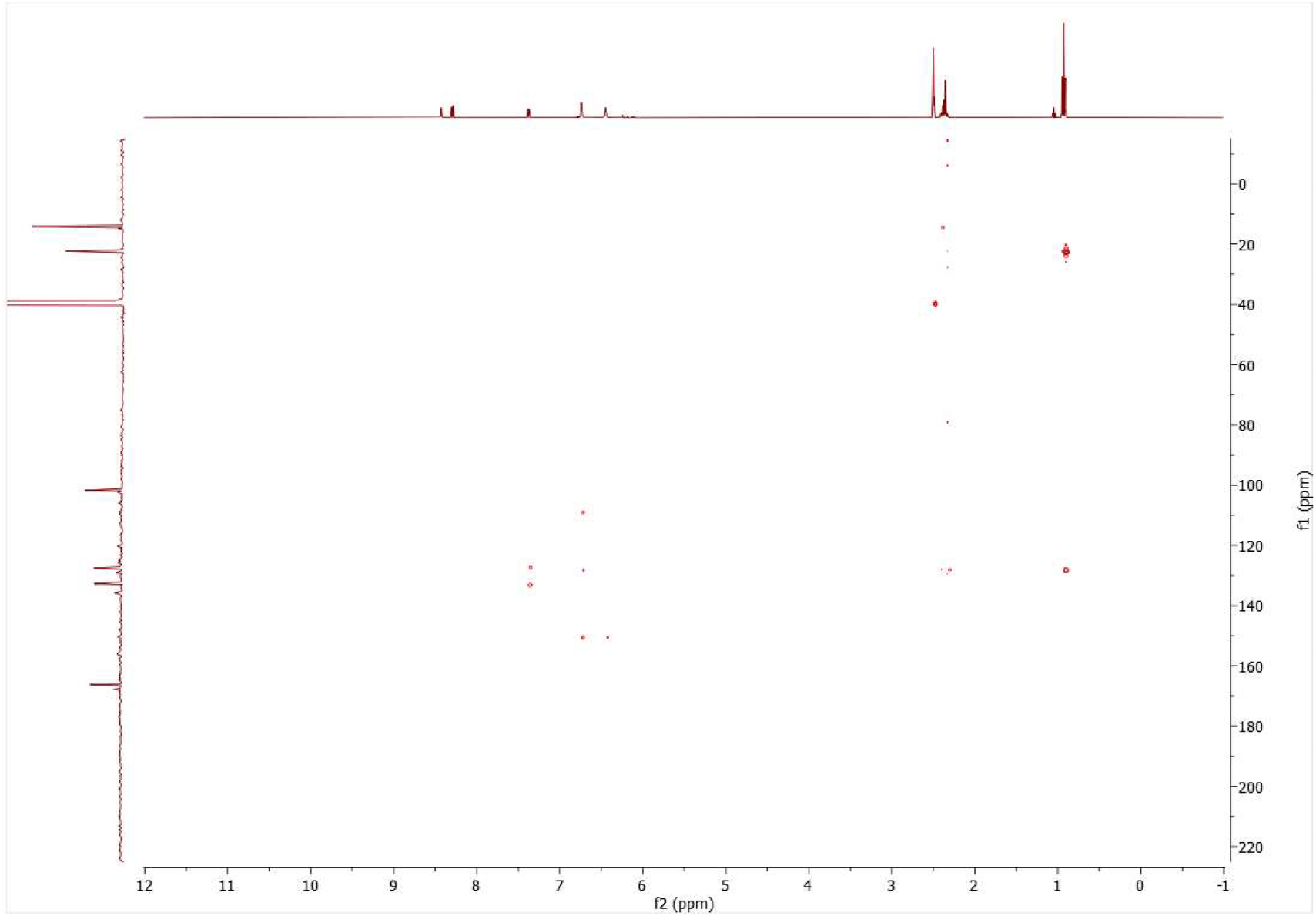
HMBC spectrum of **6** recorded in DMSO-d_6_.

**Figure S30.**
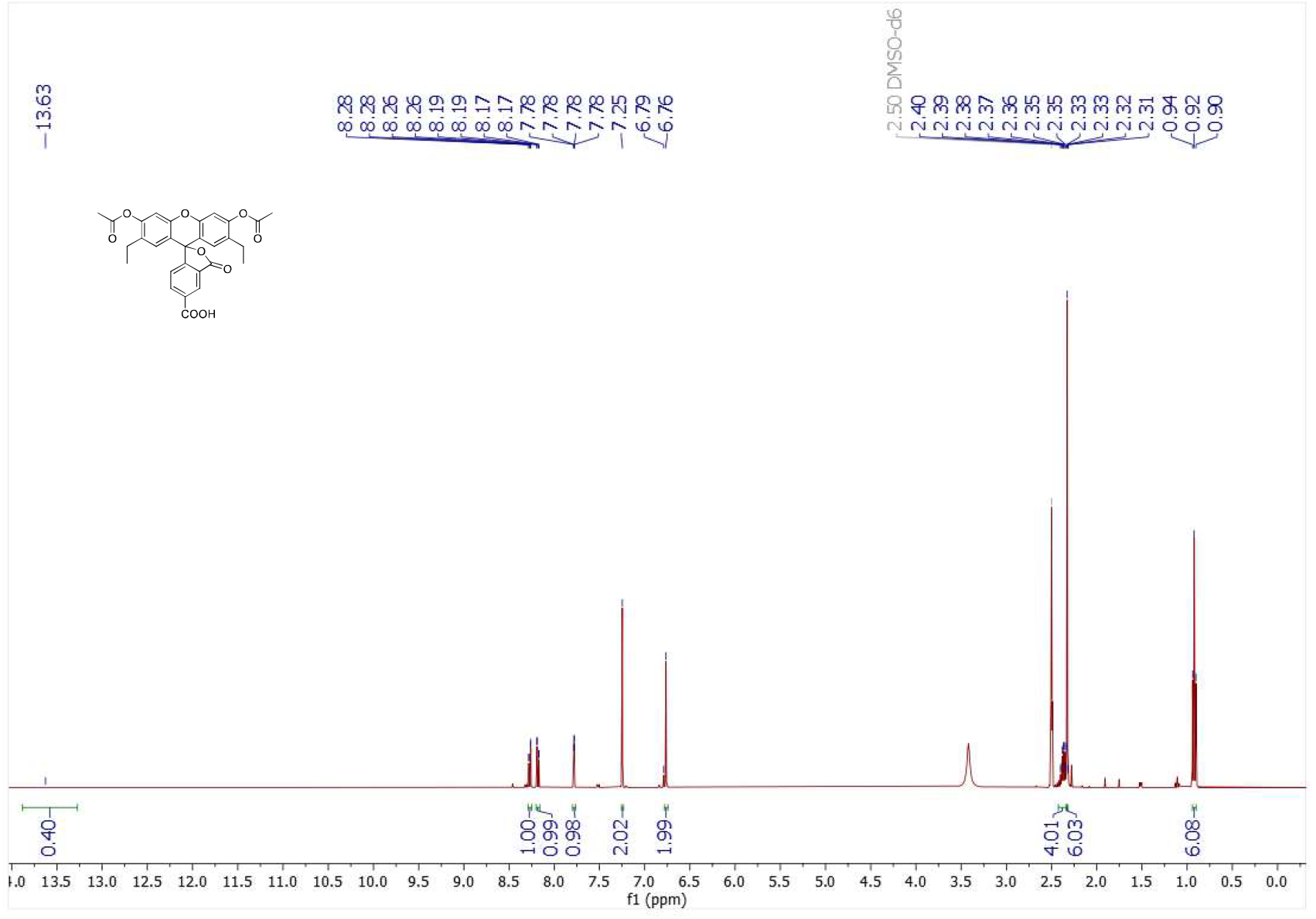
^1^H NMR spectrum of **3** recorded at 400 MHz in DMSO-d_6_.

**Figure S31.**
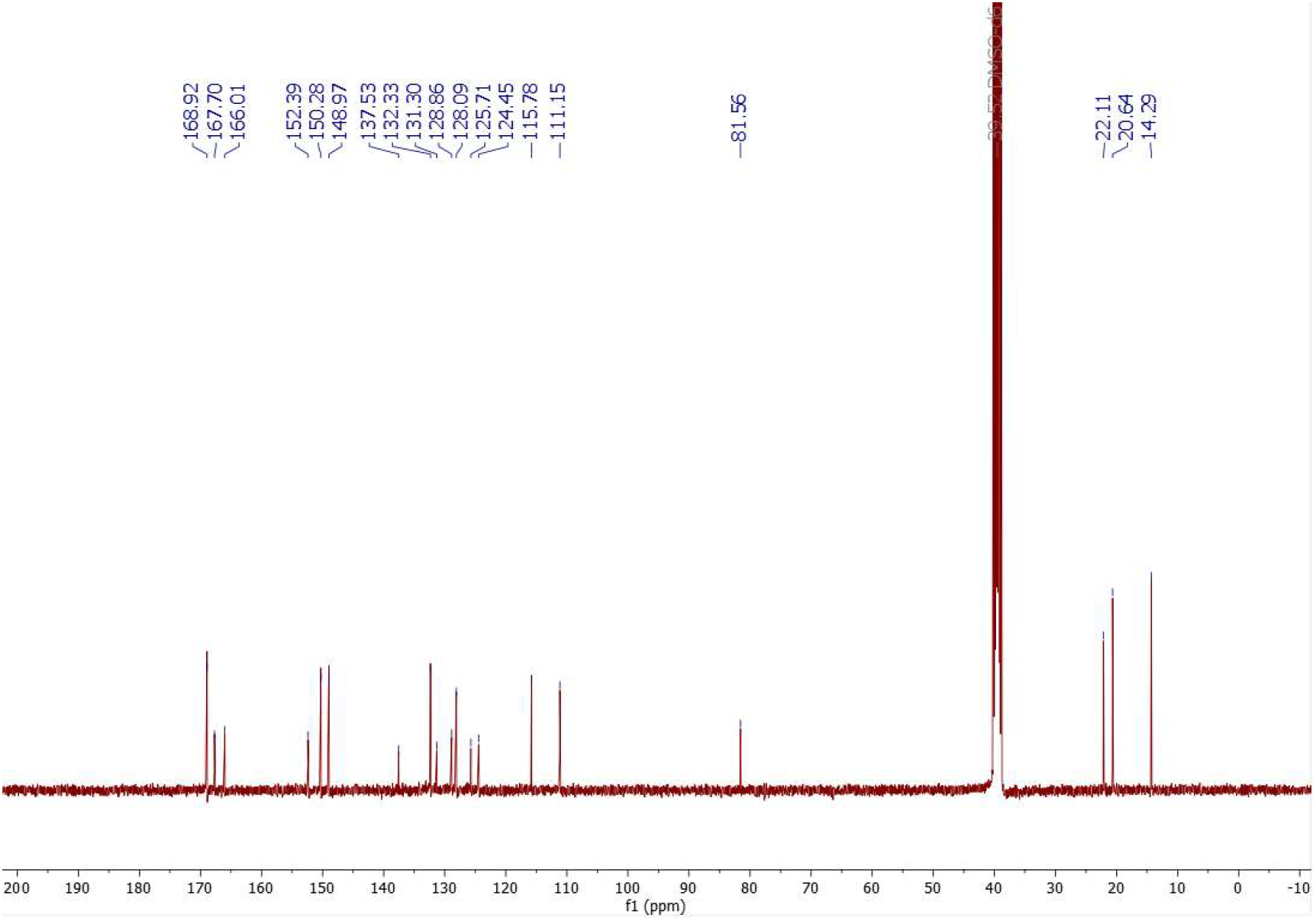
^13^C NMR spectrum of **3** recorded at 101 MHz in DMSO-d_6_.

**Figure S32.**
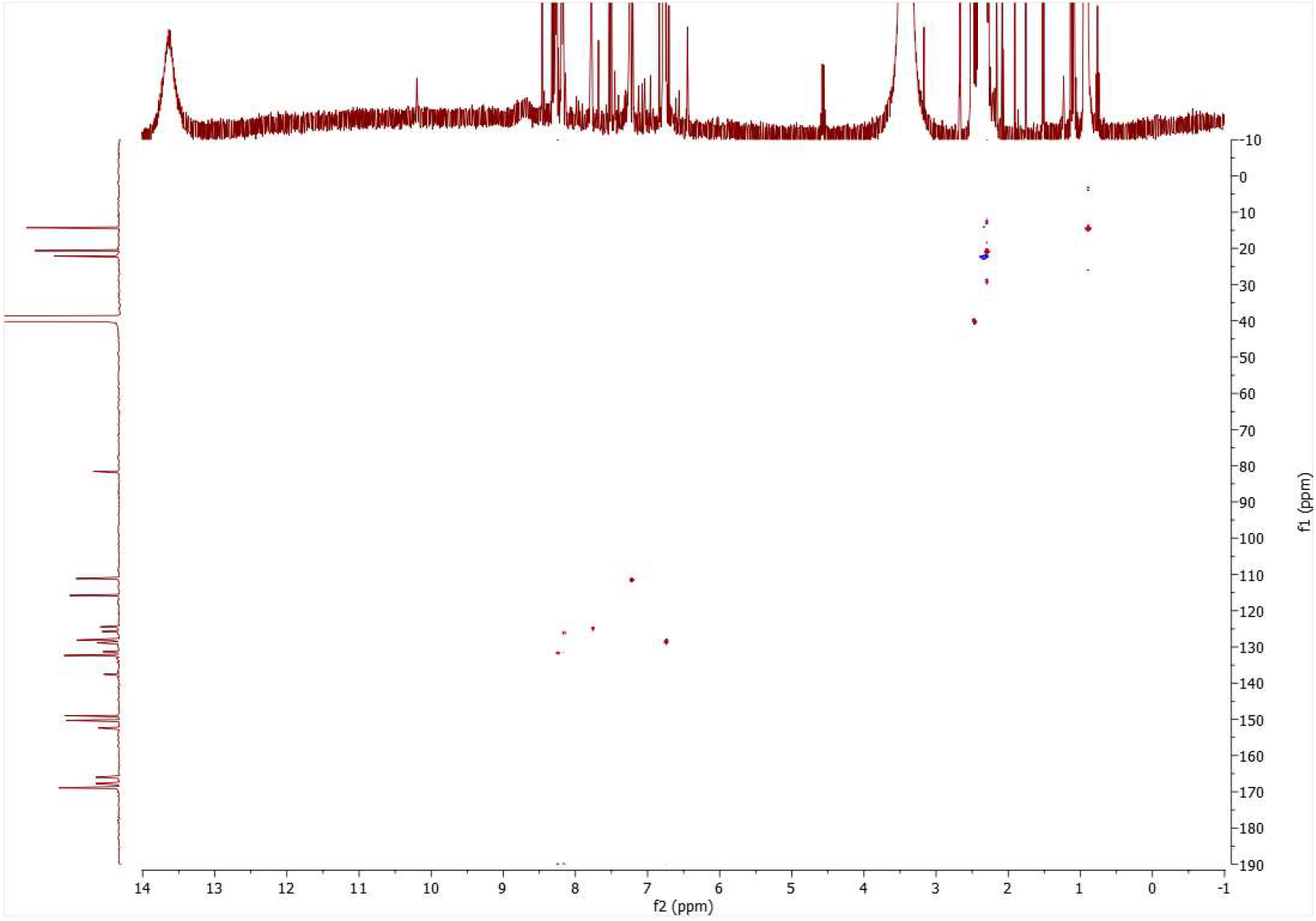
HMBC spectrum of **3** recorded in DMSO-d_6_.

**Figure S33.**
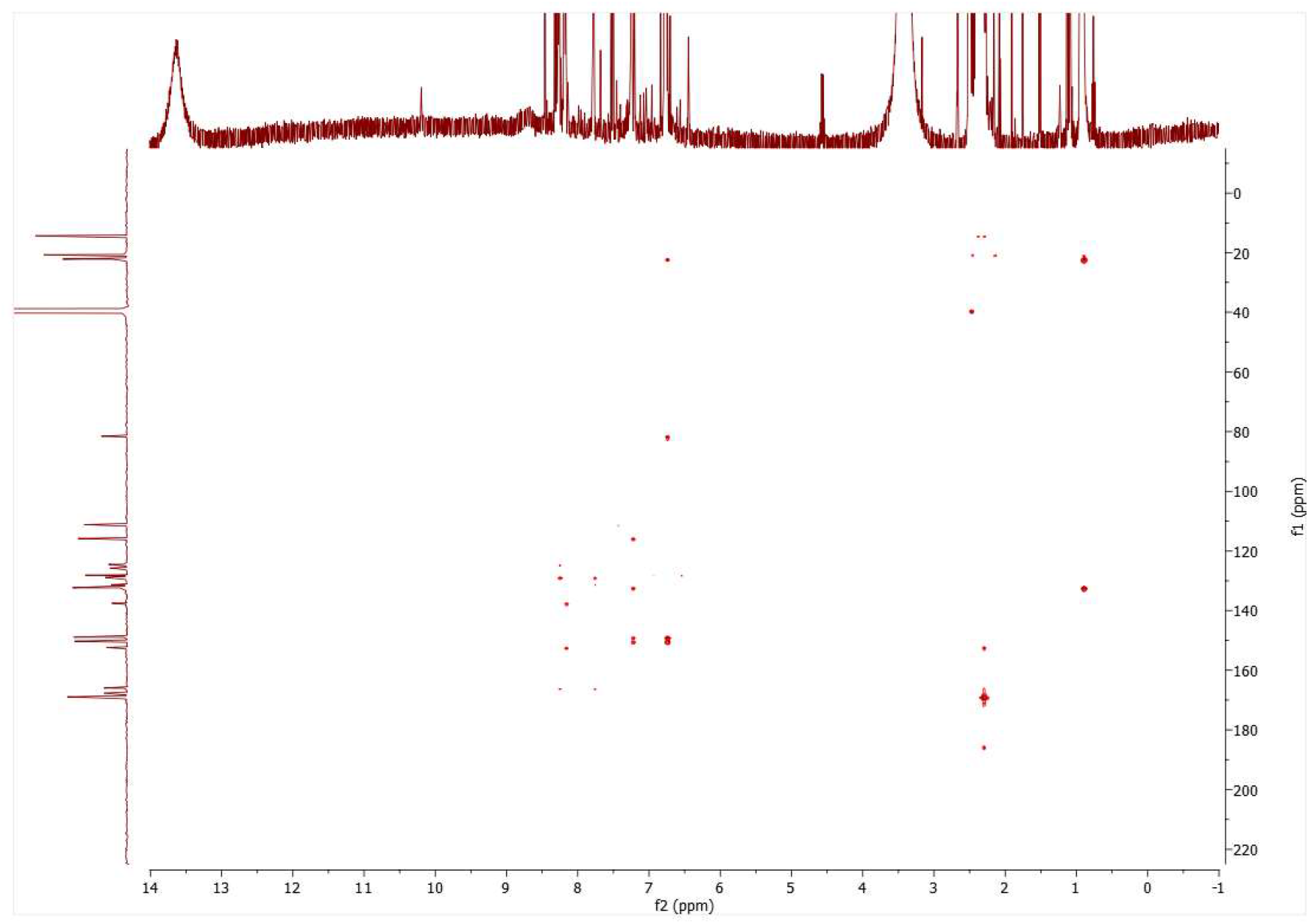
HSQC spectrum of **3** recorded in DMSO-d_6._

**Figure S34.**
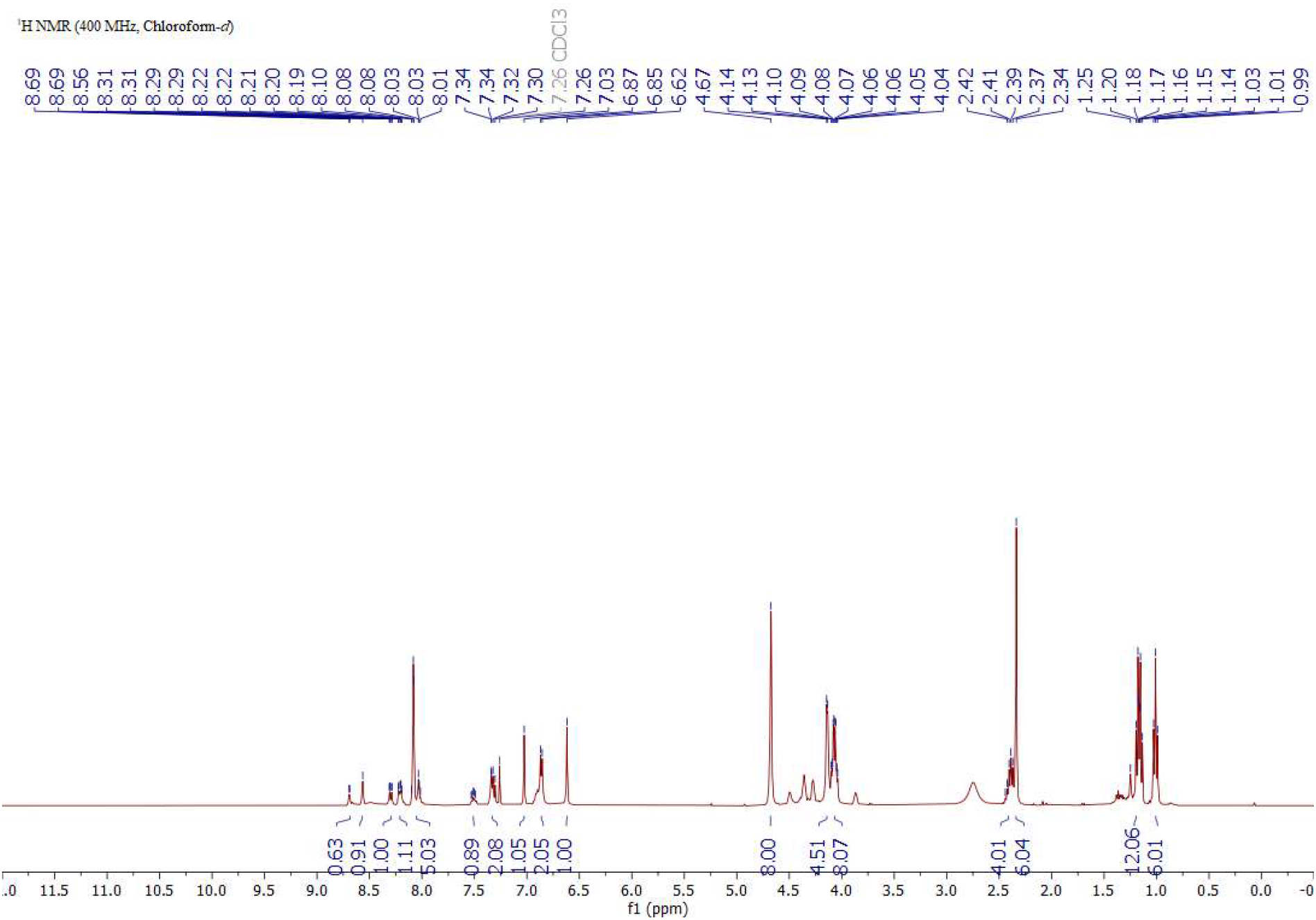
^1^H NMR spectrum of **7** recorded at 400 MHz in CDCl_3_.

**Figure S35.**
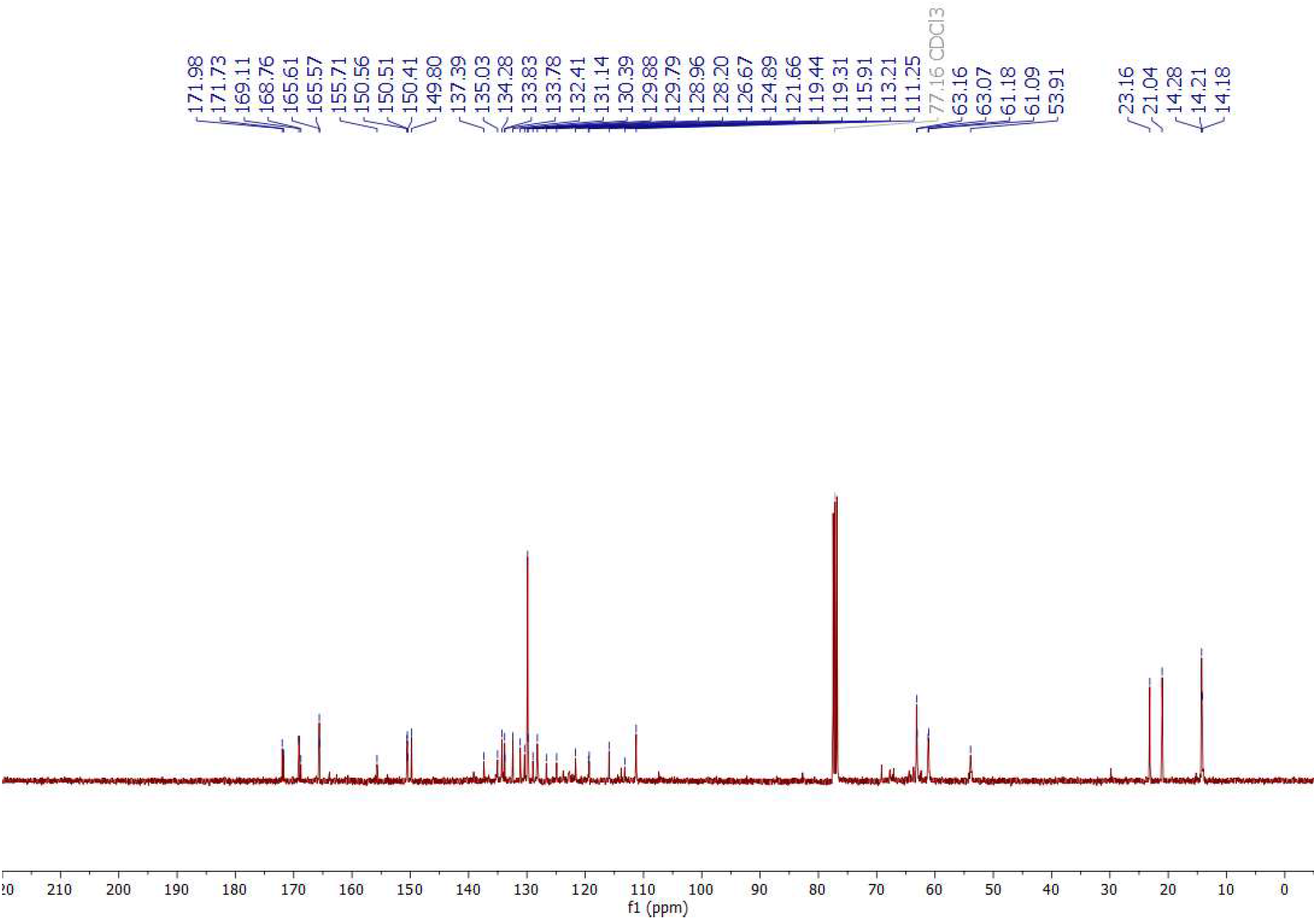
^13^C NMR spectrum of **7** recorded at 101 MHz in CDCl_3_.

**Figure S36.**
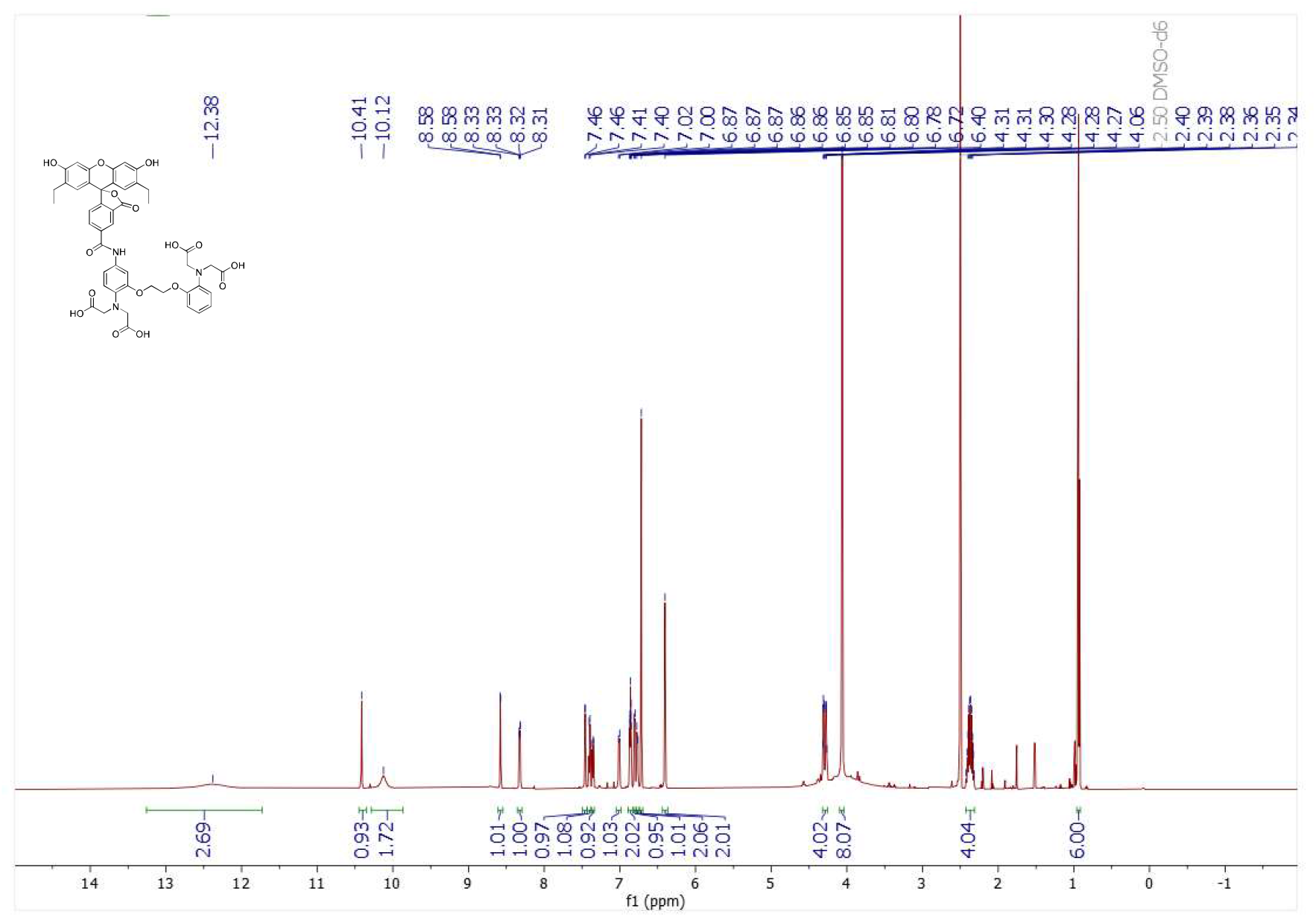
^1^H NMR spectrum of **BEEF-CP** recorded at 600 MHz in DMSO-d_6_.

**Figure S37.**
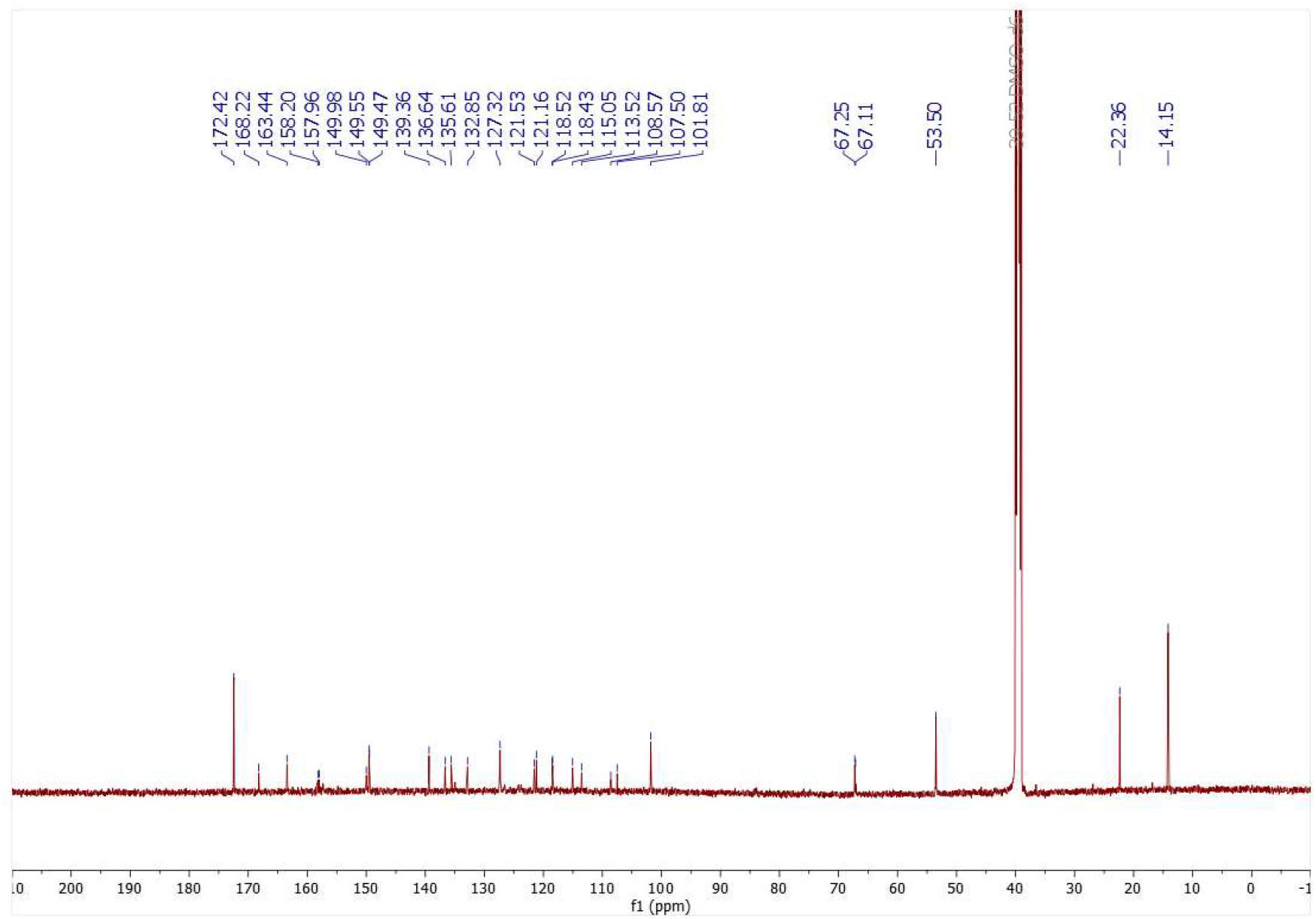
^13^C NMR spectrum of **BEEF-CP** recorded at 151 MHz in DMSO-d_6._

## References

1. Dong, E., Du, H. & Gardner, L. An interactive web-based dashboard to track COVID-19 in real time. Lancet Infect. Dis. 20, 533–534 (2020).

2. Coronavirus Disease (COVID-2019) Situation Reports. WHO (2022). Available at: https://covid19.who.int/. (Accessed: 2nd September 2022)

3. Planas, D. et al. Considerable escape of SARS-CoV-2 Omicron to antibody neutralization. Nature 602, 671–675 (2022).

4. Pacheco-Herrero, M. et al. Elucidating the Neuropathologic Mechanisms of SARS-CoV-2 Infection. Front. Neurol. 12:660087 (2021).

5. Ludwig, S. & Zarbock, A. Coronaviruses and SARS-CoV-2: A Brief Overview. Anesth. Analg. 131, 93–96 (2020).

6. Evans, J. P. & Liu, S.-L. Role of host factors in SARS-CoV-2 entry. J. Biol. Chem. 297, 100847 (2021).

7. Jackson, C. B., Farzan, M., Chen, B. & Choe, H. Mechanisms of SARS-CoV-2 entry into cells. Nat. Rev. Mol. Cell Biol. 23, 3–20 (2022).

8. Wang, K. et al. CD147-spike protein is a novel route for SARS-CoV-2 infection to host cells. Signal Transduct. Target. Ther. 5, 283 (2020).

9. Palacios-Rápalo, S. N. et al. Cholesterol-Rich Lipid Rafts as Platforms for SARS-CoV-2 Entry. Front. Immunol. 12, 12:796855 (2021).

10. Shen, X.-R. et al. ACE2-independent infection of T lymphocytes by SARS-CoV-2. Sig. Transduct. Target. Ther. 7, 83 (2022).

11. Khelashvili, G., Plante, A., Doktorova, M. & Weinstein, H. Ca^2+^-dependent mechanism of membrane insertion and destabilization by the SARS-CoV-2 fusion peptide. Biophys. J. 120, 1105–1119 (2021).

12. Hallaj, S. et al. Angiotensin-converting enzyme as a new immunologic target for the new SARS-CoV-2. Immunol. Cell Biol. 99, 192–205 (2021).

13. Nataraj, C. et al. Angiotensin II regulates cellular immune responses through a calcineurin-dependent pathway. J. Clin. Invest. 104, 1693–1701 (1999).

14. Nieto-Torres, J. L. et al. Severe acute respiratory syndrome coronavirus E protein transports calcium ions and activates the NLRP3 inflammasome. Virology 485, 330–339 (2015).

15. Hajnóczky, G., Davies, E. & Madesh, M. Calcium signaling and apoptosis. Biochem. Biophys. Res. Commun. 304, 445–54 (2003).

16. Chiovini, B. et al. Dendritic Spikes Induce Ripples in Parvalbumin Interneurons during Hippocampal Sharp Waves. Neuron 82, 908–924 (2014).

17. Verdiá-Báguena, C. et al. Coronavirus E protein forms ion channels with functionally and structurally-involved membrane lipids. Virology 432, 485–494 (2012).

18. Li, F. Structure, Function, and Evolution of Coronavirus Spike Proteins. Annu. Rev. Virol. 3, 237–261 (2016).

19. Tang, T., Bidon, M., Jaimes, J. A., Whittaker, G. R. & Daniel, S. Coronavirus membrane fusion mechanism offers a potential target for antiviral development. Antiviral Res. 178, 104792 (2020).

20. Lai, A. L., Millet, J. K., Daniel, S., Freed, J. H. & Whittaker, G. R. The SARS-CoV Fusion Peptide Forms an Extended Bipartite Fusion Platform that Perturbs Membrane Order in a Calcium-Dependent Manner. J. Mol. Biol. 429, 3875–3892 (2017).

21. He, C.-L. et al. Identification of bis-benzylisoquinoline alkaloids as SARS-CoV-2 entry inhibitors from a library of natural products. Sig. Transduct. Target. Ther. 6, 131 (2021).

22. Danta, C. C. SARS-CoV-2, Hypoxia, and Calcium Signaling: The Consequences and Therapeuti c Options. ACS Pharmacol. Transl. Sci. 4, 400–402 (2021).

23. Straus, M. R. et al. Inhibitors of L-Type Calcium Channels Show Therapeutic Potential for Treating SARS-CoV-2 Infections by Preventing Virus Entry and Spread. ACS Infect. Dis. 7, 2807–2815 (2021).

24. Danta, C. C. Calcium Channel Blockers: A Possible Potential Therapeutic Strategy for the Treatment of Alzheimer’s Dementia Patients with SARS-CoV-2 Infection. ACS Chem. Neurosci. 11, 2145–2148 (2020).

25. Alothaid, H., Aldughaim, M. S. K., El Bakkouri, K., AlMashhadi, S. & Al-Qahtani, A. A. Similarities between the effect of SARS-CoV-2 and HCV on the cellular level, and the possible role of ion channels in COVID19 progression: a review of potential targets for diagnosis and treatment. Channels 14, 403–412 (2020).

26. Chen, D. et al. ORF3a of SARS-CoV-2 promotes lysosomal exocytosis-mediated viral egress. Dev. Cell 56, 3250–3263.e5 (2021).

27. Putlyaeva, L. V & Lukyanov, K. A. Studying SARS-CoV-2 with Fluorescence Microscopy. International Journal of Molecular Sciences 22, (2021).

28. Drouillat, B., d’Aboville, E., Bourdreux, F. & Couty, F. Synthesis of 2-Phenyl- and 2,2-Diarylpyrrolidines through Stevens Rearrangement Performed on Azetidinium Ions. Eur. J. Org. Chem. 2014, 1105–1109 (2014).

29. Plaze, M. et al. Inhibition of the replication of SARS-CoV-2 in human cells by the FDA-approved drug chlorpromazine. Int. J. Antimicrob. Agents 57, 106274 (2021).

30. Rut, W. et al. SARS-CoV-2 Mpro inhibitors and activity-based probes for patient-sample imaging. Nat. Chem. Biol. 17, 222–228 (2021).

31. Ju, X. et al. A novel cell culture system modeling the SARS-CoV-2 life cycle. PLOS Pathog. 17, e1009439 (2021).

32. Csomos, A. et al. A Comprehensive Study of the Ca^2+^ Ion Binding of Fluorescently Labelled BAPTA Analogues. European J. Org. Chem. 2021, 5248–5261 (2021).

33. Cork, R. J. Problems with the application of quin-2-AM to measuring cytoplasmic free calcium in plant cells. Plant. Cell Environ. 9, 157–161 (1986).

34. Lavis, L. D. Live and Let Dye. Biochemistry 60, 3539–3546 (2021).

35. Tsien, R. & T. Pozzan, T. Measurement of cytosolic free Ca^2+^ with quin2. Methods Enzymol. 172, 230–262 (1989).

36. Chelators, Calibration Buffers, Ionophores and Cell-Loading Reagents. in The Molecular Probes® Handbook - A Guide to Fluorescent Probes and Labeling Technologies 11th Edition (2010).

37. Lin, L., Li, Q., Wang, Y. & Shi, Y. Syncytia formation during SARS-CoV-2 lung infection: a disastrous unity to eliminate lymphocytes. Cell Death Differ. 28, 2019–2021 (2021).

38. Schneider, C., Rasband, W. & Eliceiri, K. NIH Image to ImageJ: 25 years of image analysis. Nat. Methods. 9, 671–675 (2012).

39. Lien, A. & Scanziani, M. In vivo Labeling of Constellations of Functionally Identified Neurons for Targeted in vitro Recordings. Front. Neural Circuits 5:16 (2011).

40. Ma, H. et al. Wide-field in vivo neocortical calcium dye imaging using a convection-enhanced loading technique combined with simultaneous multiwavelength imaging of voltage-sensitive dyes and hemodynamic signals. Neurophotonics 1, 1–12 (2014).

41. Tada, M., Takeuchi, A., Hashizume, M., Kitamura, K. & Kano, M. A highly sensitive fluorescent indicator dye for calcium imaging of neural activity in vitro and in vivo. Eur. J. Neurosci. 39, 1720–1728 (2014).

42. Kerr, J. N. D., Greenberg, D. & Helmchen, F. Imaging input and output of neocortical networks in vivo. Proc. Natl. Acad. Sci. 102, 14063–14068 (2005).

43. Dicken, S. J. et al. Characterisation of B.1.1.7 and Pangolin coronavirus spike provides insights on the evolutionary trajectory of SARS-CoV-2. bioRxiv 2021.03.22.436468 (2021). doi:10.1101/2021.03.22.436468

44. Papalia, T. et al. Cell internalization of BODIPY-based fluorescent dyes bearing carbohydrate residues. Dye. Pigment. 110, 67–71 (2014).

45. Katona, G. et al. Fast two-photon in vivo imaging with three-dimensional random-access scanning in large tissue volumes. Nat. Methods 9, 201–208 (2012).

## References

1 A. Csomos, B. Kontra, A. Jancsó, G. Galbács, R. Deme, Z. Kele, B. J. Rózsa, E. Kovács and Z. Mucsi, A comprehensive study of the Ca^2+^ ion binding of fluorescently labelled BAPTA analogues, European J. Org. Chem., 2021, 2021, 5248–5261.

2 Lakowicz Joseph R., Effects of Solvents on Fluorescence Emission Spectra, in *Principles of Fluorescence Spectroscopy*, Springer, Boston, 2006, pp. 187–215.

3 R. Tsien and T. T. Pozzan, Measurement of cytosolic free Ca^2+^ with quin2, Methods Enzymol., 1989, 172, 230–262.

4 Chelators, Calibration Buffers, Ionophores and Cell-Loading Reagents, in The Molecular Probes® Handbook - A Guide to Fluorescent Probes and Labeling Technologies 11^th^ Edition, 2010.

5 C. Xu and W. W. Webb, Measurement of two-photon excitation cross sections of molecular fluorophores with data from 690 to 1050 nm, J. Opt. Soc. Am. B, 1996, 13, 481–491.

6 N. S. Makarov, M. Drobizhev and A. Rebane, Two-photon absorption standards in the 550-1600 nm excitation wavelength range, Opt. Express, 2008, 16, 4029–4047.

